# Nethobench: Forecasting design and conditioning shape the structural realism of generated neural dynamics

**DOI:** 10.64898/2026.10.07.757249

**Authors:** Blanche Marion, Andrés De Vicente, Mathias Mahn, Andreas Lüthi, Leon Luithlen, Catalin Mitelut

## Abstract

Which properties of neural population activity does a forecasting model reproduce? Neural forecasts are commonly evaluated by how closely they follow a held-out recording. Yet in a stochastic, partially observed system, several future trajectories are consistent with the same neural history. A forecast that averages over these possibilities can achieve low pointwise error while suppressing fluctuations that characterize individual recordings. We introduce Nethobench, a benchmark that measures structural agreement between generated and recorded neural activity. Its five metric families compare activity distributions, temporal and spectral structure, dependencies among neurons or brain regions, latent geometry, and transitions between population states. We validate these measures in stochastic systems with known dynamics and apply them to mouse and human recordings to examine how model design and conditioning on sensory and behavioral inputs shape the dynamics that forecasts reproduce. Reporting these structural profiles alongside predictive accuracy can allow researchers to compare forecasters on the properties of neural activity most relevant to their scientific question.

## 1 Introduction

Forecasting models are increasingly used as generative surrogates of neural population activity. (Azabou et al. (2023); Calcagno et al. (2024); Lueckmann et al. (2025); Duan et al. (2025)). Predicting future activity from past observations provides a fundamental test of our understanding of complex systems, including the brain, by testing whether learned patterns generalize to unseen activity. Such a test requires distinguishing three evaluation questions. Does a forecast match the observed realization? Does a predictive distribution assign appropriate probabilities under the conditioning context? And which population-level properties does generated activity preserve? Pointwise error addresses the first question. For the second, proper scoring rules assess predictive distributions (Gneiting & Raftery (2007)). The third remains less systematically addressed by existing benchmarks and requires explicit *structural diagnostics* to identify which aspects of population activity a model reproduces or distorts.

We define *structural agreement* as agreement on specified summaries under a fixed observation transform, support, and horizon. In stochastic, partially observed systems, different futures consistent with the same observed history can share these summary properties. Realizations of the same process can thus differ point by point while sharing statistical properties such as their marginal distributions and temporal dependence (Koppe et al. (2019)). Conversely, conditional-mean forecasters produce low pointwise errors, while suppressing the variability of the original realizations (Green et al. (2026)). We thus argue that structural evaluation should complement predictive-distribution evaluation, when using generative models to emulate dynamical processes.

Neuro-ethological recordings make these questions especially relevant: animals engage in spontaneous, only partially observed, behavior, where neural dynamics are non-stationary (Mazor & Laurent (2005); Sussillo & Barak (2013); Cunningham & Yu (2014); Musall et al. (2019); Stringer et al. (2019); Dennis et al. (2021)). Building generative models of these processes introduces choices that can favor different aspects of the recorded structure. One-step and multistep objectives impose different long-horizons constraints (Ben Taieb & Hyndman (2012); Benechehab et al. (2024)) ; teacher forcing and self-feedback change the histories encountered during training (Bengio et al. (2015)); and losses prioritize different statistics (Le Guen & Thome (2019); Jiang et al. (2024)). Evaluating these choices motivates the need for evaluation benchmarks that reveal which structural properties are preserved.

Existing benchmarks evaluate neural forecasts using pointwise or likelihood-based scores, including mean absolute error for calcium activity in ZAPBench (Lueckmann et al., 2025) and bits per spike for spiking activity in BrainWideBench (Andre et al., 2026). Related work on simulated neural populations shows that models can accurately reconstruct observed activity while mischaracterizing its underlying dynamics (Versteeg et al., 2025). WeatherBench 2 illustrates why structural diagnostics are useful alongside forecast error: spatial power spectra reveal the loss of fine-scale variability even when RMSE is low (Rasp et al., 2024).

Our contributions are threefold. **First**, we introduce Nethobench, a framework for comparing the statistical and dynamical properties of generated and recorded neural activity across five metric families, with empirical references to contextualize the scores. **Second**, we use controlled perturbations to test each family’s sensitivity to changes in its target properties and its robustness to changes in other properties. **Third**, we demonstrate Nethobench’s diagnostic and scientific utility by identifying which properties neural forecasts preserve and testing how model selection affects the recovery of stimulus-side decodability.

## 2 Method and Metrics

We formalize Nethobench as an evaluation framework for comparing reference and predicted neural activity, designed to complement pointwise fidelity or probability distribution metrics. It reports structural agreement in five metric families (Figure 1), each addressing a different question about neural population activity: which activity levels occur, over what timescales they fluctuate, which signals covary, what geometry the population occupies, and how it moves between recurrent states. They each reflect established approaches to characterizing neural population activity in systems and computational neuroscience (Cohen & Kohn (2011); Murray et al. (2014); Buzsáki & Mizuseki (2014); Cunningham & Yu (2014); Mazzucato et al. (2015); Elsayed & Cunningham (2017)).

**Figure 1:**
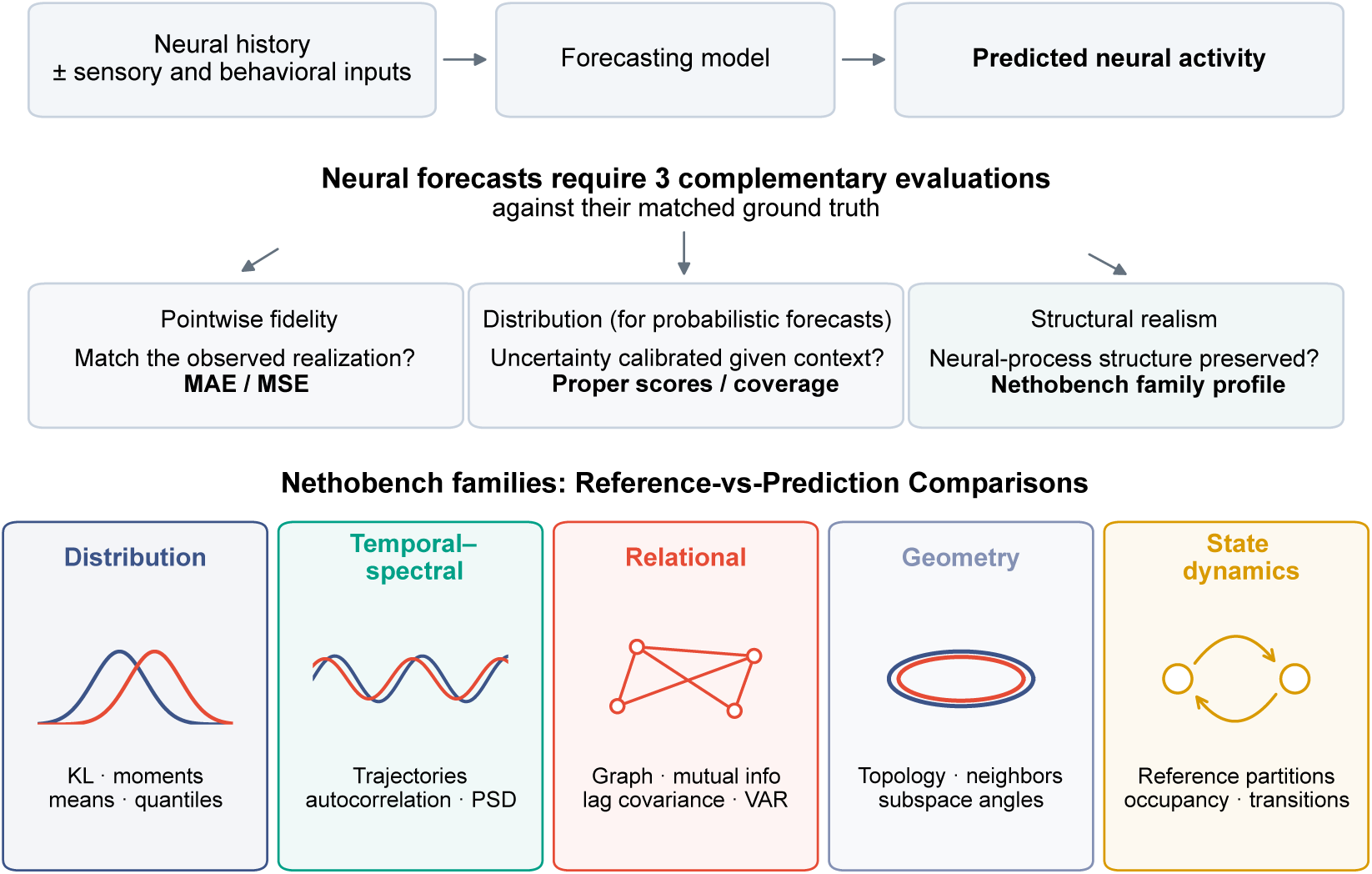
Overview of neural forecasting challenges and Nethobench framework. Five metric families assess distributional, temporal-spectral, relational, geometric, and state-dynamical agreement with reference activity. These structural profiles complement pointwise accuracy and predictive-distribution evaluation.

Let *X, Y* R*^M^*^×^*^T^* ^×^*^R^* denote reference and predicted activity, respectively, with *X_mtr_* and *Y_mtr_* indexed by the same sequence *m*, time point *t*, and recorded region or unit *r*. We compare *X* and *Y* using the following families:

**Distribution.** This family tests whether forecasts preserve the marginal statistics of recorded activity. For example, let *p_mr_* and *q_mr_* be the reference and forecast histograms for region *r* in sequence *m*, computed on the same bins. We compare them through the symmetric divergence 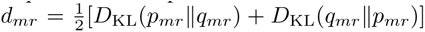 and map it to agreement as (1 + *d_mr_*)^−1^. Separate comparisons of means, moments, and tail quantiles identify shifts in baseline activity, amplitude compression, and loss of rare events.

**Temporal-spectral.** This family tests whether forecasts preserve how activity evolves over time. For a given sequence and region, let *p_f_* and *q_f_* denote normalized reference and forecast power spectra over nonzero frequencies, and let *P_X_* and *P_Y_* denote their total powers. For positive total powers, spectral agreement is

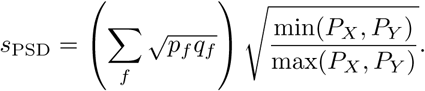

The first factor measures agreement in spectral shape, while the second detects a change in overall power even when the shape is preserved. The family also compares latent trajectories and the autocorrelation of both activity and its increments.

**Relational.** This family tests whether forecasts preserve dependencies among regions. Let *E_X_* and *E_Y_* contain the strongest 20% of edges by absolute correlation in the reference and forecast networks. Their overlap, *E_X_ E_Y_ / E_X_ E_Y_*, measures whether the same pairs of regions remain strongly coupled. The graph component also compares degree and clustering. Mutual information, lagged covariance, and a fitted one-step VAR response provide complementary summaries of population dependence. The lagged covariance and VAR comparisons pool sequences and retain one matrix triangle; Appendix C.6 examines a within-sequence, directed variant.

**Geometry.** This family tests whether forecasts preserve the geometry of population activity. If *θ*_1_*, . . ., θ_k_* are the principal angles between the dominant reference and forecast covariance subspaces at their common retained rank *k*, subspace agreement is 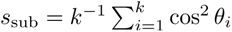. A complementary manifold component compares persistence lifetimes and local neighborhood distances, which can differ even when the dominant subspaces align.

**State dynamics.** This family tests whether forecasts visit and move between latent states with the same frequencies as recorded activity. We assign both datasets to clusters fitted on the reference embedding. For state labels *z_t_*, we compare occupancy frequencies *p*^*_i_* = Pr(*z_t_* = *i*) and joint transition frequencies 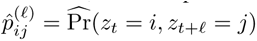 at several lags *ℓ*, counting pairs within sequences.

**Aggregation and reporting.** Together, these five families comprise 18 components, each mapped to an agreement score in [0, 1]. We average these scores within each family, then combine the five family scores, *F_D_, F_T_, F_R_, F_G_, F_S_*, into a composite:

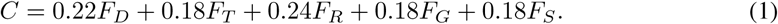

These weights are design choices, whose influence we assessed through 5,000 weight draws and explicit tests of equal weighting, omission of each family, and doubling of each family’s weight. Across all settings, three checkpoints retained the same ranking by mean composite across 15 seeds (Figure S1, Appendix C.3). We nevertheless consider the family profiles to be the primary output of Nethobench, since the same composite can arise from different combinations of preserved and lost structure. We return to this point in Section 3.

When a component cannot be calculated, we omit it and renormalize the remaining weights within its family. All omissions are reported, with component definitions, hyperparameters, and missingness rules detailed in Appendix C.1.

Although Nethobench applies across modalities, its scores should primarily be interpreted within the data and evaluation setting from which they arise. We thus use real-real and shuffled comparisons to contextualize agreement for each dataset and window length. These provide empirical references for higher and lower agreement, rather than strict upper and lower bounds (Figure S2).

**Metric sensitivity and robustness.** To assess whether the metrics capture their intended properties, we first examine signals whose differences are known. Independent realizations of the same stochastic process receive high structural scores despite poor pointwise agreement (Figure 2a). Targeted perturbations produce distinct family profiles, with gain and bias changes mainly affecting distributional and state-dynamical scores, and coupling rewiring most strongly affecting relational agreement (Figure 2b). Time shuffling preserves distributional agreement while reducing temporal agreement, and variance contraction lowers distributional, temporal-spectral, and state-dynamical scores (Figure 2c-d). The temporal–spectral and state-dynamical families also broadly agree with angular DSA on model rankings, with differences in sensitivity to controlled perturbations (Appendix C.2.1). Additional temporal controls and component audits appear in Figure S4 and Fig. S5, respectively. Sensitivity analyses examine how scores and rankings depend on PCA dimensionality, state counts, and lags (Figures S1 and S7).

**Figure 2:**
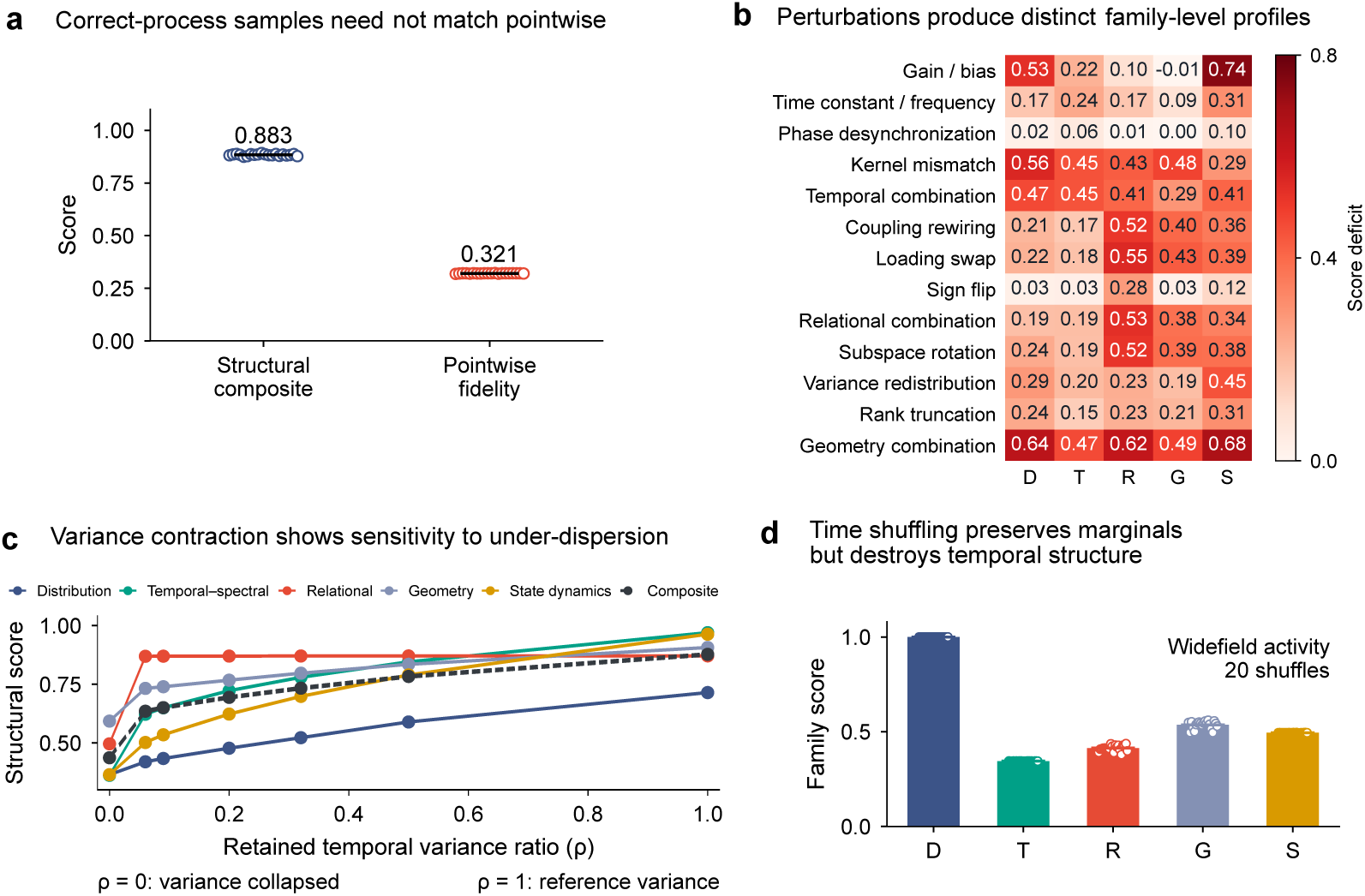
Controlled validation of structural diagnostics. (a) 20 oracle realizations scored with structural agreement (Nethobench) and pointwise fidelity. Produced by a controlled generator using a latent first-order autoregressive process and a calcium observation transform. (b) Mean oracle score minus the maximum-dose perturbed score. The perturbation inventory changes gain/bias; time constants/frequency; phase synchronization; observation kernels; combinations of temporal parameters; coupling; channel loadings; signs; relational combinations; subspace orientation; variance allocation; rank; and geometric combination. (c) Effect of mean-preserving temporal-variance contraction across the five families. 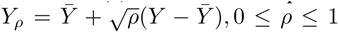 where means are taken over time within region/window. (d) Family scores after regional time shuffling across 20 shuffles. Appendix C.2 provides the full definitions and implementation details behind (a-d).

## 3 Validation and Results

Our validation experiments focus primarily on establishing what Nethobench adds to the evaluation of neural forecasts, with less emphasis on the relative performance of individual models. Model comparisons serve this aim when they help explain which properties distinguish the forecasts and why those differences matter.

We thus consider that the utility of Nethobench should serve a dual purpose: one *diagnostic*, and the other *(neuro)scientific*. *Diagnostic* because it should help one identify what is gained or lost through forecasting model design choices. *Scientific* because its intended use is for biological systems, where the value of a mathematical approximation ultimately depends on how faithfully it reproduces the phenomenon under study. Identifying which statistical properties of neural activity a generator preserves is a first step ; but preserving these properties should ideally translate into improved capture of condition-specific neural activity.

### 3.1 On their diagnostic utility

A preliminary experiment towards justifying the diagnostic utility of Nethobench can ask what its metrics reveal when pointwise error is held fixed. We thus applied five different distortions to some widefield calcium recordings (Appendix B.1) and matched the resulting sequences in MSE. The distortions include time-order change, channel shifts, smoothing, a PCA shortcut and state cycling (Appendix C.2). As hypothesized, different distortions produced different structural profiles (Figure 3a). For example, temporal reordering yielded a low temporal score (0.70) and a high relational score (0.97), whereas the PCA shortcut showed the reverse pattern (0.93 and 0.77, respectively). Omitting the temporal-spectral family reduced the composite’s sensitivity to temporal reordering, whereas omitting the relational family reduced its sensitivity to the PCA shortcut (Figure S6).

**Figure 3:**
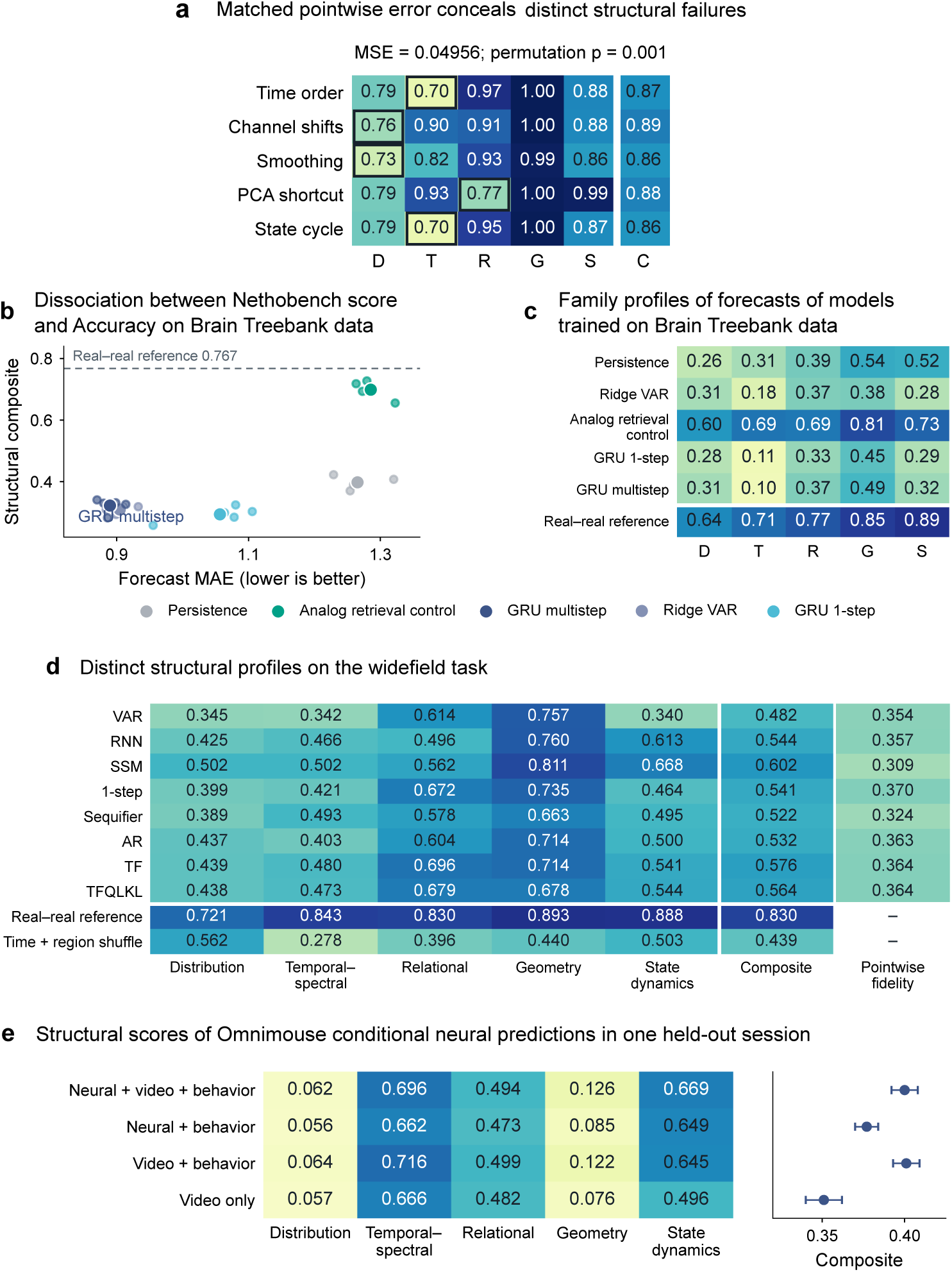
Validation of structural diagnostics. (a) Effect of five amplitude-matched widefield transformations on metric families. The family-profile differences are tested with 999 blocked permutations, yielding p = 0.001. (b) MAE and Nethobench composite score of forecasts across five Brain Treebank participants (small points) and equally weighted method means (large points) (c) Family scores averaged across the five Brain Treebank participants. The dashed line in (b) and last row in (c) show the empirical real-real reference: symmetric comparisons of alternating 25-window halves, averaged within participants. (d) Nethobench family and composite means of the widefield model suite. Real-real and time-and-region shuffle references use the same targets. (e) Nethobench family and composite means across four sequence-level splits in one held-out session of the Omnimouse dataset.

Up to now, we knew by construction how two signals differ. The next step is to ask whether the same metrics could identify meaningful differences between forecasts when the source of those differences is unknown. We thus assess forecasters using structural and pointwise scores on generated samples - as design choices are often described to result in different learned dynamics (Ben Taieb & Hyndman (2012); Benechehab et al. (2024); Bengio et al. (2015); Jiang et al. (2024); Le Guen & Thome (2019)).

This experiment includes forecasting experiments on different data modalities, serving different questions:

1. **When MAE and structural agreement favor different forecasters, which properties distinguish their forecasts?** Using electropysiological recordings from intracranial probes in humans (Wang et al. (2024))
2. **At similar pointwise fidelity, which properties change with the training regime, and how does that affect model selection?** Using widefield calcium imaging recordings in head-fixed mice (Saxena et al. (2020); Mitelut et al. (2022))
3. **With the model fixed, which properties change when conditioning inputs are added?** Using single-neuron spiking activity inferred from two-photon calcium recordings, captured alongside behavioral and stimulus input (Fahey et al. (2026); Willeke et al. (2026))
4. **4. Does the evaluation criterion change model rankings across datasets?** All of above and electrophysiological recordings from Neuropixel probes in different labs (International Brain Laboratory (2024))

**When MAE and structural agreement favor different forecasters, which properties distinguish their forecasts?** We evaluated forecasters on high-gamma activity from 64 intracranial contacts on 5 participants of Brain Treebank dataset. For each of them, we held out one of three movie runs in turn, trained on the other two, and predicted the next 20s of activity from 5s of neural history. We chose simple forecasters whose predictions are easy to interpret: persistence, ridge VAR, GRUs trained with one-step or multistep objectives, and analog retrieval (the implementation details are provided in Appendix B.3). MAE and the Nethobench composite ranked two forecasters at opposite extremes

1. The multistep GRU, trained to minimize error over the forecast window, scores lowest on both MAE and Nethobench Composite.
2. Analog retrieval, which returns the recorded continuation of a similar neural history, had the highest MAE yet scored highest on each of Nethobench five metric families (Figure 3bc).

The structural aspect of neural data Analog retrieval learns that multistep GRU does not is principally its temporal structure (T of 0.69 against 0.10). This comparison shows a concrete use case where Nethobench complements MAE by identifying which properties of recorded population activity a forecast reproduces, even when it diverges from the observed trajectory.

**At similar pointwise fidelity, which properties change with the training regime, and how does that affect model selection?** After Brain Treebank experiment, we were naturally interested in asking whether structural differences also emerge when two models share an architecture and achieve similar pointwise fidelity. We evaluated eight forecasters on 101 hours of widefield calcium recordings from 16 cortical regions in six mice. Given 3s of recorded activity, each model generated a 24s rollout using its own predictions or latent states. The suite contains a linear autoregressive baseline (**VAR**), a recurrent neural network (**RNN**), a stochastic latent state-space model (**SSM**), and five Transformers. These include two next-token models (**1-step** and **Sequifier**), two sequential models trained with teacher forcing (**TF**) or self-feedback (**AR**), and a teacher-forced variant with additional tail-quantile and symmetric-KL losses (**TF QL KL**). The dataset and models specifications are detailed in Appendices B.1 and D.

Many of those provide good use cases for one-to-one comparisons (Figure 3d, Tables 9 and 10); notably **TF** and **AR** models, that differ only in their training regime. Strikingly, their mean pointwise fidelity was nearly identical, yet **TF** scored higher on both relational (0.696 vs 0.604) and temporal-spectral structure (0.480 vs 0.403) while both having the same mean geometry score. The family profiles also let us ask whether the model with the highest composite is the best choice for a particular property. Across the full suite, sampled **SSM** forecasts achieved the highest mean composite and led on four of the five families. However, the relational score favored **TF** (0.696 vs 0.562). For an analysis focused on inter-regional dependencies, the relational score would therefore favor **TF**, whereas the composite would favor the **SSM**. Within this suite and task formulation, family profiles show which aspects of a forecast change with the training regime even when pointwise fidelity barely changes, and which model best preserves a property of interest.

**With the model fixed, which properties change when conditioning inputs are added?** These comparisons give some empirical intuition on the impact of model and training choices on the structure a forecast preserves. What if we now held the model fixed and varied only the information available at inference? We evaluated pretrained OmniMouse predictions for 128 neurons in one held-out naturalistic video session. We divided the session into 60 sequences of 3 s each and scored only the 2s forecast window. The experimental set up is further described in Appendix B.2.

Adding behavior to video increased mean scores across all five families and raised the composite from 0.351 to 0.401. The largest gain was in state dynamics (0.496 to 0.645). Adding video to neural history and behavior also increased all five family means, raising the composite from 0.377 to 0.400 (Figure 3e and Table 12). Adding neural history to video and behavior had a different effect. The composite barely changed (0.401 to 0.400), yet the mean state-dynamics score rose (0.645 to 0.669) and the mean temporal-spectral score fell (0.716 to 0.696).

**Does the evaluation criterion change model rankings across datasets?** Ideally, the same model comparisons should recur across recordings. To test that hypothesis, we fit three published architectures (**POCO** (Duan et al. (2025)), **TSMixer** (Chen et al. (2023)), and **NetFormer** (Lu et al. (2025))) and two baselines (per-channel ridge, denoted **Linear**, and a **GRU**) on IBL, widefield, Brain Treebank, and OmniMouse recordings. Within each dataset, all models predicted the same held-out activity from neural history alone. We ranked the five models by mean MAE and by mean Nethobench composite, then averaged their ranks across datasets (Table 1). POCO and TSMixer had the same mean MAE rank (2.50), yet the composite ranked POCO above TSMixer and NetFormer on all four datasets. We also observed a ranking reversal across individual animals: in 15 of 17 held-out IBL mice, POCO achieved lower MAE than a Linear baseline, while the Linear baseline achieved higher structural agreement. We report the implementation details and scores across all components in Appendix B.7.

**Table 1:** P**o**intwise **error and structural agreement across four neural forecasting datasets.** Within each dataset, models were evaluated on the same held-out targets using neural history alone. Comp. denotes the Nethobench composite. Mean ranks give equal weight to each dataset and rank the five models shown (1 = best). Bold marks the best fitted value in each column. RR, and TR denote the real-real, and time-and-region shuffle references, respectively.

|  | IBL |  | Widefield |  | Brain Treebank |  | OmniMouse |  | Mean rank |  |
|---|---|---|---|---|---|---|---|---|---|---|
| Model | MAE↓ | Comp.↑ | MAE↓ | Comp.↑ | MAE↓ | Comp.↑ | MAE↓ | Comp.↑ | MAE↓ | Comp.↓ |
| POCO | **0.686** | 0.460 | 0.7215 | **0.494** | 0.832 | **0.4157** | 0.343 | **0.367** | 2.50 | **1.25** |
| TSMixer | 0.781 | 0.431 | **0.7207** | 0.488 | 0.894 | 0.3951 | 0.331 | 0.337 | 2.50 | 2.50 |
| NetFormer | 1.130 | 0.308 | 0.7480 | 0.420 | 1.774 | 0.3486 | 0.604 | 0.334 | 5.00 | 4.75 |
| Linear | 0.710 | **0.470** | 0.7212 | 0.467 | 0.868 | 0.3913 | **0.310** | 0.360 | **2.00** | 2.50 |
| GRU | 0.826 | 0.409 | 0.7214 | 0.474 | **0.820** | 0.3907 | 0.359 | 0.319 | 3.00 | 4.00 |
| RR | 0.798 | 0.734 | 1.0650 | 0.754 | 1.114 | 0.7070 | 0.444 | 0.507 | — | — |
| TR | 1.202 | 0.468 | 1.0391 | 0.471 | 1.077 | 0.6031 | 0.428 | 0.354 | — | — |

**Table 2:** Stimulus-side decoding from recorded and forecast hippocampal activity across 23 IBL sessions. Separate decoders were fitted on the same held-out trials. The final column reports the absolute difference between forecast and recorded balanced accuracy, computed within each session and then <u>averaged across sessions.</u>

| Population activity | Mean balanced accuracy (%) | Mean absolute difference from recordings (pp) |
|---|---|---|
| Recorded responses | 61.6 | — |
| Nethobench-selected forecasts | 75.8 | **14.7** |
| MAE-selected forecasts | 83.4 | 21.9 |
| Poisson-selected forecasts | 96.4 | 34.9 |

### 3.2 On their (neuro)scientific utility

In OmniMouse, access to biologically relevant information improved structural agreement. This naturally leads us to ask whether the opposite is true. If we select a forecaster for its structural agreement, does its output more faithfully reproduce a stimulus-related property of the recorded population? If this is the case, we have empirical examples of transfer from more accurate conditioning to a high score, and of transfer from a high score to better downstream performances.

We tested this hypothesis in hippocampal recordings from the International Brain Laboratory, where population activity carries information about whether a visual stimulus appeared on the left or right (International Brain Laboratory (2024)). We identified 23 sessions from multiple laboratories in which stimulus side could be decoded from recorded spiking activity. For each session, we trained forecasters to predict the hippocampal response given the stimulus and recent neural activity. From the same candidates, we selected models by validation Nethobench composite, MAE, or Poisson log loss. We then decoded stimulus side from recorded and forecast activity between 100 and 400 ms after stimulus onset, fitting a separate decoder to each on the same held-out trials. We report further experimental details on the training task and model suite in Appendix B.4.1.

Mean balanced accuracy for decoding stimulus side was 61.6% in the recordings and higher in all three sets of forecasts: 75.8% for models selected by Nethobench, 83.4% for MAE, and 96.4% for Poisson log loss. To assess agreement with the recordings, we computed the absolute difference in decoding accuracy within each session. We obtain the the smallest mean discrepancy with Nethobench (14.7 percentage points, compared with 21.9 for MAE and 34.9 for Poisson log loss). The criteria also consistently favors different model families. In all sessions, Nethobench selects a stimulus-conditional mean corrected using the trial’s neural history, while MAE and Poisson select GRUs.

Because all three selections overestimate mean decodability, even forecasts unrelated to the trial’s stimulus side might come closer to the recorded accuracy. To test this, we reassigned entire forecasts among trials within the same session and at the same absolute stimulus contrast. Decoding falls to near chance for all three criteria (46.5-48.1%), indicating that the original forecasts carried information about the target trial’s stimulus side. Only for Nethobench did the original pairing yield a smaller mean discrepancy than reassignment, and the difference is just 0.97 percentage points. Thus, Nethobench-selected forecasts come closest to the recorded level of decodability, although preserving the trial pairing improved that agreement only slightly.

Across calcium imaging and electrophysiology, the five metric families identified which properties of recorded population activity were best preserved in forecasts. Further controls include a small two-photon dataset of the kind often collected in systems neuroscience, where forecasts of amygdala activity showed stronger agreement with the recordings in population geometry than in activity distributions (Fig. S23). In IBL, structural agreement declined when forecasters were evaluated on recordings from mice in held-out laboratories (Figs. S21-S22). Shuffling synthetic BOLD signals generated from known dynamics also reduced structural agreement (Fig. S24). We interpret each profile within its recording setting because the observation process determines which fluctuations in neural activity the metrics can capture (Wei et al. (2020)).

## 4 Discussion

Our results gather empirical evidence that Nethobench effectively rewards forecasts that preserve the structure of the neural activity they are meant to emulate and identifies which properties are lost. We observed this sensitivity across recording modalities, animals, and datasets, in models ranging from standard baselines to established neural forecasting architectures. Nethobench thus complements measures of pointwise fidelity and proper scores for predictive distributions. Together, these evaluations give a fuller account of forecast quality by asking how closely a forecast matches the observed realization, how well the model characterizes uncertainty given the context, and which population-level properties its forecasts preserve.

One should however interpret these scores within their scope: given our definition of structural agreement - defined as concerning selected summaries under a specified observation transform and evaluation window -, preserving it through a high Nethobench score does not establish recovery of a generating process, appropriate uncertainty or causal mechanisms. A control performed on Brain Treebank retrieval illustrates this distinction: recorded continuations can preserve population structure without being appropriate futures for a given context (Appendix B.6 and FigureS18). Proper probabilistic scores, conditional calibration and task-specific validation therefore remain necessary.

Additionally, some mthodological choices further delimit the sensitivity of the scores. PCA-based components can miss weak but behaviorally relevant signals, and clustered states should not be equated with cognitive states. Comparisons across neural populations require additional alignment assumptions, while dependence on preprocessing and window length calls for experiment-specific real-real and shuffled references. Families are also partly correlated, but their differing perturbation responses support diagnostic complementarity.

An immediate next step for NeuroAI would be a large-scale evaluation of recently proposed forecasting architectures and pretrained models across neural datasets. Scoring their forecasts on the same held-out recordings with MAE, RMSE, and Nethobench would show how often pointwise and structural criteria favor different models and which properties of population activity distinguish their forecasts. In systems neuroscience, Nethobench could help evaluate models intended as experimental surrogates by testing whether their forecasts reproduce measured changes in population activity across behavioral or intervention conditions.

## AI USE STATEMENT

Generative AI tools assisted with analysis and plotting code, numerical consistency checks, figure preparation, and manuscript editing. Quantitative figures are based on recorded or simulated data; conceptual schematics are illustrative. AI assistance did not replace verification against the experimental records, and the authors are responsible for the final text, methods, and claims.

## Ethics statement

The benchmark evaluates processed neural recordings and secondary analyses of public datasets. Human analyses use the de-identified Brain Treebank release and do not attempt participant identification. Structural scores are not measures of clinical validity, cognition, or causal mechanism, and the evaluated models are not validated for clinical decision-making. Dataset access and reuse remain subject to the originating datasets’ terms and governance. Reporting empirical references, component availability, and failure cases reduces the risk of treating a single benchmark score as evidence of biological equivalence.

## Reproducibility statement

Appendices B–E specify dataset splits, preprocessing, model objectives, score transforms, component availability, and uncertainty units. The figures retain model- and seed-level controls alongside aggregate scores. We identify the evaluation implementation and distinguish fixed-checkpoint comparisons, training replications, corruption realizations, and participant-level inference; these sources of variation are not pooled into a single uncertainty estimate.

## Appendix

- Appendix A: Related Work
- Appendix B: Experimental Set Up and Datasets
- Appendix C: Validation and robustness analyses
- Appendix D: Models
- Appendix E: Code, Data and Reproducibility
- Appendix F: Supplementary Figures

## A Related Work

**Predictive modeling and neural forecasting.** Predictive modeling is central to systems and computational neuroscience. In encoding models, neural responses are predicted from external variables such as stimuli, actions, or task variables; in decoding models, stimuli, behavior, or latent variables are predicted from neural population activity (Kriegeskorte & Douglas (2019); Glaser et al. (2020); Mathis et al. (2024)). These approaches test what information neural populations carry, but predictive success does not imply recovery of the mechanisms or dynamics that generated the activity. Recent work has extended this paradigm in scale and scope, from large-scale neural population decoders to foundation-style models that predict brain responses under naturalistic multimodal stimulation (Azabou et al. (2023); Wang et al. (2025); d’Ascoli et al. (2026)).

Neural forecasting is a narrower predictive setting in which the target is future neural activity itself, generated from recent neural history, and potentially, stimulus and behavioral variables (Antoniades et al. (2024); Calcagno et al. (2024); Duan et al. (2025)). When recordings are viewed as samples from a stochastic dynamical process, forecasting models can therefore serve as generative surrogates of neural dynamics.

**Neural benchmarks and the scope of Nethobench.** Existing neural benchmarks - including Brain-Score, Sensorium, Dynamic Sensorium, the Neural Latents Benchmark, and NeuralBench - standardize the evaluation of models predicting held-out neural, behavioral, or stimulus variables (Schrimpf et al. (2020); Turishcheva et al. (2024a;b); Pei et al. (2022); Banville et al. (2026)), but they are not designed to assess the long-horizon structure of generated neural activity. Recent forecasting benchmarks - such as ZAPBench, PoCo, and BrainWideBench (Lueckmann et al. (2025); Duan et al. (2025); Andre et al. (2026)) - move closer to this setting, while emphasizing pointwise measures such as correlation and mean squared error, or probabilistic measures such as likelihood. Pointwise metrics require trajectory alignment, whereas probabilistic measures evaluate the predicted distribution. But neither provides a family-resolved account of which structural properties are preserved.

Complementary methods treat neural activity as a representation or dynamical system. RSA, CKA, and generalized shape metrics compare representational geometry under different invariances (Kriegeskorte et al. (2008); Kornblith et al. (2019); Williams et al. (2022)). Dynamical Similarity Analysis more directly compares the temporal organization of computation across neural systems (Ostrow et al. (2023)), while optimal-transport approaches compare noisy, stochastic neural population dynamics without reducing them to deterministic trajectories (Nejatbakhsh et al. (2024)). This system-level perspective has also begun to enter benchmarking. The Computation-through-Dynamics Benchmark similarly shows in controlled settings that reconstruction accuracy need not imply recovery of the correct dynamics (Versteeg et al. (2025)).

Nethobench is designed for this complementary evaluation problem. Its scope is not all neural modeling, and not all predictive modeling. It specifically targets generative neural forecasting models, evaluating them over long rollout horizons. Within this scope, Nethobench asks which structural properties of the reference neural process are preserved by the generated activity. This emphasis parallels evaluation practices in other domains where forecasts are assessed on structural or physical properties alongside predictive accuracy, notably WeatherBench 2 in weather forecasting (Rasp et al., 2024) and PDEBench in modeling physical systems governed by partial differential equations (Takamoto et al., 2024).

## B Experimental Set Up and Datasets

### B.1 Widefield calcium forecasting

The processed widefield dataset contains 287 sessions of approximately 21 minutes from six GCaMP6 transgenic male mice (Ai93 and Ai94), totaling approximately 101 hours Mitelut et al. (2022); Saxena et al. (2020). During the recordings, the head-fixed mice performed an uncued, self-initiated lever-pull task for a water reward, with a 3-second unrewarded lockout period imposed after each successful pull. Neural activity was captured via widefield calcium imaging through a chronic transcranial window at 30 Hz with a spatial resolution of 68 *µ*m/pixel.

Following rigid alignment to a 2D projection of the Allen Common Coordinate Framework v3, locaNMF was applied to decompose the imaging data into anatomical components. Consequently, activity is summarized over 16 cortical regions at 30 Hz. We use these regional calcium time series rather than raw images; the benchmark does not test image segmentation, anatomical registration, or spike inference. The processed release is identified as `Nethobench/nethobench-widefield-v1` and distributed under CC-BY-4.0. Recording identifiers define the data partitions.

We split the data at the original recording-sequence level: all subsequences derived from a given recording are assigned to exactly one split. Formally, letting *s*(*i*) denote the parent recording sequence of example *i*, we enforce

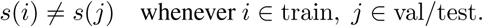

This prevents temporally adjacent subsequences from the same recording from appearing in different splits.

Training examples pair 90 context samples with a 90-sample target block. Non-overlapping blocks are constructed within recordings, and longer evaluation targets concatenate consecutive blocks without crossing recording boundaries.Channel normalization uses training-context mean and standard deviation, with numerical stabilization 10^−6^, and is applied unchanged to evaluation data. The principal comparison scores only the 720 generated samples following 90 observed samples: 24 s of forecast from 3 s of context. For each model and forecast horizon, scores are computed separately on four disjoint groups of evaluation sequences and averaged within each training seed. We report the mean and sample standard deviation across three training-seed means. This SD describes training variability on the fixed evaluation set, rather than variability across animals or evaluation groups. Horizon analyses score the first 3, 12, and 24 seconds of generated activity, excluding the observed context.

Widefield references use the same 222 evaluation targets as the model comparison, excluding context and retaining the full 720-bin forecast window. Each of 20 random draws is scored jointly within four fixed evaluation groups of 56/56/55/55 sequences, using the same evaluator as the model comparisons. Real-real companions are finite 720-bin windows sampled from the same recording without overlapping their paired target window. Temporal shuffling independently permutes time within each channel, whereas pooled shuffling permutes all time-channel entries within each sequence. Scores are averaged equally across the four groups and then across draws (Figure S2). Their values calibrate this observation scale and support; they are not attainable upper and lower bounds on all forecasts. A separate five-checkpoint robustness panel uses 100 sequences of 630 bins and includes context in its evaluation tensor (Appendix C.3).

**Evaluation records and provenance.** The evaluation pipeline saves component, family, composite, and pointwise-fidelity scores for each model, training seed, and evaluation split, with separate records for each forecast horizon. Structural and pointwise scores use the same prediction–reference arrays and evaluation groups. JSON caches and CSV exports retain the individual scores, selected sequence indices, and aggregation summaries. Reported variability is the sample SD across training-seed means after averaging the four evaluation groups within each seed; within-seed split variability is exported separately. Family-level sensitivity to context length, number of regions, and training duration is shown in Figure S12; component-level and model-specific diagnostics are shown in Figures S11, S13, and S14.

### B.2 OmniMouse conditional neural prediction

We evaluate the public OmniMouse-80M checkpoint (Willeke et al., 2026) on one held-out naturalistic-video session containing 128 evaluated neurons. Four configurations retain neural history, video and behavior; neural history (single-neuron spiking activity inferred from two-photon calcium fluorescence) and behavior; video and behavior; or video alone. Behavioral variables condition predicted neural activity; behavioral outputs are not scored. All configurations use the same targets, neuron panel, preprocessing and four non-overlapping sequence-level evaluation splits.

**Input timing and prediction task.** We reconstruct the inference protocol from the published repository. Let *t* = 0 denote the beginning of the neural target interval. Each model call use a two-second input window, nominally spanning [1, 1) s, with neural responses and stimulus video sampled at 30 Hz and behavioral covariates at 20 Hz. Neural targets are input-window samples 30-59, corresponding to [0, 1) s. For the two configurations using neural history, only samples 0-24 are encoded, with sample times from 1 to 0.2 s. The remaining configurations receive no neural response values as context. These timings describe the configured sampling grid.

When behavior conditioning is enabled, all 40 behavioral samples from the two-second window are encoded, including observations during the neural target interval. When video conditioning is enabled, the complete 60-frame clip enters the video encoder. The `causal_response_video_ behavior` strategy subsequently selects video features using the configured frame range 30-59, whereas `video_behavior` and `stimulus_driven` retains features for the full clip (Table 3). Because selection occurs after video encoding, the configured range does not restrict the encoder’s raw input to target-interval frames. Accordingly, “causal’ refers to the neural response prefix and does not imply past-only availability of every conditioning modality.

**Table 3:** **OmniMouse modality-conditioning configurations.** Inputs refer to the first model call.

| **Manuscript configuration** | **Saved strategy** | **Neural input** | **Video processing** | **Behavior input** |
|---|---|---|---|---|
| Neural history + video + behavior | causal_response_video_behavior | Samples 0–24 | Full 60-frame clip encoded;<br>subsequent feature selection<br>configured as frames 30–59 | All 40 samples |
| Neural history + behavior | causal_response_behavior | Samples 0–24 | None | All 40 samples |
| Video + behavior | video_behavior | None | Full 60-frame clip encoded;<br>full-clip features retained | All 40 samples |
| Video only | stimulus_driven | None | Full 60-frame clip encoded;<br>full-clip features retained | None |

**Two-step prediction.** The primary protocol (Figure S20b) uses a three-second source window nominally spanning [1, 2) s, through two successive model calls. The first call predicts neural responses over [0, 1) s. The second shifts the input window forward by one second and predicts [1, 2) s. For neural-history configurations, the selected population’s neural context is replaced with first-call predictions, of which the first 25 samples form the encoded prefix. Video and behavioral inputs remain recorded observations in each shifted window. Thus, prediction is recursive in neural responses only for the neural-history configurations, while enabled covariates remain observed throughout the predicted interval.

**Scoring support.** The primary protocol scores the concatenated 60 predicted samples against the corresponding recorded activity, excluding the initial recorded context. The separate native-horizon protocol (Figure S20a) concatenates 30 recorded context samples with one 30-sample prediction and compares this with the same recorded context followed by the recorded target. Both protocols therefore score 60-sample sequences. This support does not meet the default tail-quantile component’s 80-sample requirement, so available-component aggregation must be interpreted with this limitation.

Tables 11 and 12 retain the supplied aggregate means and SEMs across the four splits. The realreal reference uses distinct recorded sequences; temporal, time-and-region and factorized controls retain different amounts of structure. They cannot be interpreted as interchangeable null hypotheses. Generalization across sessions, mice, behavioral variables or conditioning protocols is not established by this single-session comparison.

### B.3 Brain Treebank human intracranial recordings

**Cohort and observation transform.** We use Brain Treebank (Wang et al., 2024), NEMAR `nm000253`, version 1.0.0 (DOI: `10.82901/nemar.nm000253`). The selection includes participants with at least three available runs, taking the first three in acquisition order: participants 01–04, runs 01/02/03, and participant 06, runs 01/02/05. These are 15 participant–movie runs totaling 32.1 hours, not a claim of 15 distinct movies.

Contacts must have BIDS-good status in all three runs. Before common-average referencing, gross quality control excludes contacts whose pooled first-difference median absolute deviation is above 20 times or below 1/20 of the within-participant candidate median. The calculation samples differences at a stride of 2048 original samples. We retain 64 contacts by evenly spaced acquisition-order indices. This contact panel and gross QC use all three runs, including the held-out run; they are not a completely held-out-blind preprocessing pipeline. They do not use forecasting errors or retrieval outcomes.

The 2048 Hz signals undergo common-average referencing, fourth-order zero-phase Butterworth filtering in 70–115 and 125–150 Hz bands, and Hilbert-power extraction. Band powers are averaged, resampled to 16 Hz, and log-transformed. Each contact is normalized by its training-movie median and 1.4826 times its training-movie MAD, and clipped to [8, 8]. A sample with log power below 20 in more than 25% of selected contacts marks a dropout; any context-plus-future window containing it is excluded. Normalization and training PCA use unflagged training samples. Zero-phase filtering is an offline observation transform, so the forecasts are not a prospective causal signal-processing benchmark.

**Splits, retrieval and prediction.** Within each participant, one movie is held out and the other two provide training data. Each held-out run contributes 50 non-overlapping windows after excluding 60 s at each edge. A window contains 80 context samples (5 s) and 320 future samples (20 s), giving 750 evaluation windows overall. Persistence repeats the last observed vector. Ridge VAR uses 16 lags, ridge coefficients from 0.1, 1, 10, and a maximum fitted spectral radius of 0.995. Analog retrieval projects observed contexts into a 16-component whitened PCA representation fitted on training movies, retrieves the nearest training context, and outputs its recorded continuation. Both the 5-s context and the 20-s continuation in the library come exclusively from the two training movies. The held-out movie and future do not enter the library or nearest-context selection. This control tests whether a training continuation passes structural checks; it is not evidence of conditional correctness.

The GRUs have two layers, 128 hidden units and dropout 0.1. Training uses seeds 17, 29 and 43, batch size 64, learning rate 5 10^−4^, weight decay 10^−4^, gradient clipping at 1, a maximum of 60 epochs, and early-stopping patience 8. Candidate windows advance by 16 samples. Each epoch uses 4096 training windows; validation uses 512 windows from a 10% portion of each training movie. Training/validation window boundaries are not purged of all potential overlap; this affects the independence of checkpoint selection within training movies, while the third movie remains excluded from fitting. The multistep objective supervises horizons 1, 4, 16 and 64; the one-step model uses the next observation. Both generate autonomously at evaluation.

**Scoring and inference.** Temporal metrics use 16 Hz. Each deterministic method contributes one evaluation per fold, while each GRU contributes three training-seed evaluations: 135 model/fold/seed rows. Scores average seeds within movie, movies within participant, and then the five participants equally. Persistence produces an undefined ACF score in 15 rows; removing ACF from every method preserves model ordering and gives a multistep-minus-one-step composite difference of approximately 0.0280. The real–real reference averages the two directions of comparing alternating halves of the 50 evaluation windows (25 versus 25); its mean composite is 0.767.

For the primary GRU contrast, the participant is the independent unit. The paired hierarchical bootstrap resamples participants and then their movies, after averaging seeds within movie and keeping methods paired (10,000 replicates, seed 20260917). It yields Δ*C* = 0.0285, 95% interval [0.0180, 0.0415]. Enumerating all 2^5^ = 32 participant sign assignments gives two-sided *p* = 0.0625. These summaries are both reported: a positive bootstrap interval does not override the resolution of the five-participant randomization test. Leave-one-participant-out analyses test dependence on individual participants, and the 45 movie/seed contrasts display within-participant variation.

**Human corruption controls.** For each held-out fold, four transformations are evaluated at *α* 0, 0.25, 0.5, 0.75, 1, giving 15 4 5 = 300 evaluations. Additive noise uses *Y* = *X* + *αɛ*, with standard-normal *ɛ*. Temporal block permutation divides each 320-bin future into ten 32-bin (2 s) blocks, selects a nested random-priority prefix of round(10*α*) blocks, and cyclically remaps it jointly across contacts. Channel-wise circular shifts use fixed contact-specific base shifts scaled by *α*. Lower-variance attenuation uses training-fitted PCA and scales components 33–64 by 1 *α*, retaining the other components; PCA is fitted on at most 100,000 valid training samples. The removed half accounts for 21.3–40.3% of training variance. It is not infinitesimal variance or independently identified task-relevant information. Severity zero is identity. One nested corruption realization per fold and type is held fixed across doses. Participant-mean dose correlations are descriptive associations across five doses, not independent tests at each point. The principal Brain Treebank results, participant/component robustness, and controlled-corruption calibration are shown in Figures S15, S16, and S17, respectively.

### B.4 IBL Neuropixels Repeated Site Experiment

The data is published in International Brain Laboratory (2024)

#### B.4.1 Stimulus-side decoding in recorded and forecast hippocampal activity

**Sessions and forecasting task.** We used hippocampal spiking activity recorded with Neuropixels probes during the International Brain Laboratory visual decision-making task. We focused on 23 sessions in which stimulus side could be decoded from the recorded population response after correction for multiple comparisons (*q <* 0.05). Across these sessions, balanced accuracy in the initial analysis ranged from 60.7% to 89.4%. For the forecast comparison, we estimated recorded accuracy again using only the forecasting test trials. This gave each set of predictions a recorded reference based on exactly the same trials (Table 4).

**Table 4:** Stimulus-side decoding from hippocampal population activity. The initial analysis reports balanced accuracy across the trials used to identify sessions with decodable stimulus side. The final four columns report balanced accuracy (%) on the same forecasting test trials, using either recorded responses <u>or forecasts from the model selected by each validation criterion.</u>

|  | Initial | Forecasting test trials |  |  |  |  |
|---|---|---|---|---|---|---|
| Session | BA (%) | Trials | Recorded | Nethobench | MAE | Poisson |
| NR_0024 | 81.5 | 55 | 72.8 | 100.0 | 90.9 | 98.2 |
| NR_0027 | 76.0 | 84 | 67.7 | 73.4 | 88.0 | 96.3 |
| NR_0019 | 74.0 | 53 | 55.8 | 69.2 | 100.0 | 100.0 |
| CSHL059 | 71.9 | 67 | 68.7 | 73.1 | 82.1 | 98.5 |
| UCLA037 | 64.8 | 55 | 49.3 | 79.3 | 76.0 | 90.3 |
| CSH_ZAD_022 | 62.2 | 57 | 54.3 | 66.7 | 89.6 | 96.4 |
| DY_018 | 89.4 | 57 | 85.6 | 98.5 | 95.8 | 100.0 |
| CSH_ZAD_026 | 80.6 | 78 | 74.2 | 85.9 | 83.7 | 82.5 |
| KS084 | 80.4 | 45 | 80.1 | 97.8 | 93.4 | 100.0 |
| DY_010 | 77.5 | 34 | 73.6 | 76.0 | 73.6 | 100.0 |
| KS074 | 72.5 | 56 | 61.0 | 56.4 | 100.0 | 100.0 |
| NYU-21 | 71.6 | 74 | 70.4 | 94.6 | 79.6 | 79.6 |
| NYU-26 | 70.2 | 102 | 52.5 | 77.9 | 72.2 | 95.8 |
| CSHL051 | 67.8 | 47 | 65.7 | 87.2 | 80.7 | 100.0 |
| DY_009 | 67.2 | 35 | 54.1 | 71.1 | 82.7 | 100.0 |
| UCLA011 | 66.5 | 69 | 50.7 | 85.1 | 93.6 | 98.3 |
| KS044 | 64.7 | 43 | 54.4 | 53.4 | 97.9 | 100.0 |
| DY_016 | 64.6 | 36 | 57.5 | 68.1 | 62.5 | 100.0 |
| KS023 | 63.1 | 59 | 50.6 | 55.5 | 67.1 | 100.0 |
| NYU-30 | 62.2 | 69 | 62.4 | 66.0 | 75.1 | 83.9 |
| ZFM-01936 | 62.1 | 53 | 55.3 | 76.8 | 80.0 | 98.3 |
| NYU-37 | 61.0 | 38 | 57.9 | 65.8 | 76.3 | 100.0 |
| ibl_witten_13 | 60.7 | 74 | 41.7 | 64.9 | 78.5 | 100.0 |

We fitted forecasters separately for each session. The neural context extended from 100 ms before to 100 ms after stimulus onset, and the forecast extended from 100 to 1,000 ms after onset. Candidates included GRUs with different hidden sizes and training checkpoints, ridge vector autoregressive models, a stimulus-conditional mean, and a stimulus-conditional mean with a neural-history correction. The conditional mean predicted the average training response for the presented stimulus. Its history-dependent variant added a regularized linear correction estimated from the spiking activity observed on the current trial. For each session, we selected models by validation MAE, Poisson log loss, or Nethobench composite and froze their predictions before decoding.

**Decoding and comparison with recorded activity.** We restricted decoding to the first 300 ms of the forecast, from 100 to 400 ms after stimulus onset. Within each session, we fitted four separate left-versus-right decoders, one on recorded population responses and one on the forecasts selected by each criterion. We used the same five cross-validation folds across all four representations. Thus, each decoder was trained and evaluated within its own representation, and all decoding folds consisted exclusively of trials held out from forecaster training and selection.

We measured decodability by balanced accuracy, the mean of the accuracies on left- and right-stimulus trials. For each criterion, we took the absolute difference between forecast and recorded balanced accuracy separately for each session, then averaged those differences with equal weight across sessions. Taking the absolute value within sessions prevents overestimation in one session from cancelling underestimation in another. The session-level comparison also shows that the ordering varied. For example, in KS044, Nethobench-selected forecasts closely reproduced recorded accuracy (53.4% vs. 54.4%), while in NR 0024 they yielded perfect decoding against a recorded accuracy of 72.8%.

**Stimulus-side negative control.** We asked how much of the observed decodability depended on pairing forecasts with the stimulus side of their target trials. For each of 2,000 permutations, we reassigned entire predicted population sequences among forecasting test trials within the same session and absolute stimulus contrast. Each target trial retained its original side label. This preserved the temporal and population structure of each forecast and the contrast of its assigned trial, while randomizing the association between the forecast and the target trial’s stimulus side. We refitted the decoders for each permutation using the original five folds and fixed decoding hyperparameters. We kept the recorded reference accuracy fixed throughout.

Reassignment reduced mean balanced accuracy to 46.5–48.1% across criteria (Table 5). Its effect on agreement with the recordings depended on the selection criterion. For Nethobench, the original pairing reduced the mean absolute discrepancy by 0.97 percentage points relative to reassignment. For MAE and Poisson log loss, the original pairing increased it by 5.67 and 18.01 points, respectively. These comparisons are descriptive. Because the forecasters received the stimulus as input, the control establishes that their outputs carried information about the target trial’s stimulus side. Whether the same neurons or population patterns carried that information in forecasts and recordings remains open.

**Table 5:** Effect of trial reassignment on stimulus-side decoding. Balanced accuracy is averaged across sessions. Discrepancy is the mean absolute within-session difference from recorded balanced accuracy, in per<u>centage points. Reassigned values are averaged over 2,000 permuta</u>tions. Balanced accuracy (%) Discrepancy (points) Selection criterion Original Reassigned Original Reassigned

|  | Balanced accuracy (%) |  | Discrepancy (points) |  |
|---|---|---|---|---|
| Selection criterion | Original | Reassigned | Original | Reassigned |
| Nethobench | 75.8 | 48.1 | 14.68 | 15.65 |
| MAE | 83.4 | 47.3 | 21.86 | 16.19 |
| Poisson log loss | 96.4 | 46.5 | 34.85 | 16.84 |

#### B.4.2 Population Forecasts

**Data and support.** The repeated-site IBL cohort (International Brain Laboratory, 2024) comprises 17 sessions, four laboratories and 2176 stimulus-aligned trials. Twelve regional population-rate channels are binned at 20 ms, transformed as log(1 + rate in Hz), and normalized using training-context statistics. Each sequence contains 25 context bins and 75 future bins (0.5 and 1.5 s). A region requires at least five quality-filtered units for the anatomical-coverage count. Missing regional channels are zero-filled before normalization in the stored tensors, so a common 12-channel shape does not imply equal regional coverage. The within-session, held-out-session and held-out-laboratory splits contain 436, 512 and 128 evaluated target contexts, respectively. They are fixed partitions, not independent draws of laboratory splits.

**Forecast models and temporal controls.** The population examples compare PSTH, VAR, latent-dynamical VAR and a regularized Transformer. The latter adds mean, 90th-quantile and lag-product penalties to MSE; the stored model identifier does not denote a negative-binomial likelihood. The GRU and latent SSM-VAE transfer comparisons retain training seeds 7, 11 and 19. Connected summary points score their ensemble-averaged predictions, rather than averaging the seed-level scores. These structural transfer panels score all 100 bins, including copied context. Pearson horizon comparisons and all probabilistic analyses instead exclude context. Their absolute scores are not directly comparable.

Order controls operate on the 436 within-session target tensors of shape 75 12 at 50 Hz. A partial shuffle selects round(75*α*) bins per trial from a nested random order and permutes them jointly across channels. Selected positions may remain fixed by chance. Jitter applies contact-specific circular shifts bounded by *M* (*α*) = round(*α* 75*/*8); shifts are round(*UM*) for fixed *U* [1, 1] across doses. Maximum jitter is nine bins (180 ms). Error bars summarize three corruption RNG seeds (7, 11, 19), not three independently sampled animals or model fits.

**Probabilistic evaluation.** We evaluate *K* = 18 predictive draws per context on the target only. For target vector *x* and draws *y_k_*, the empirical energy score is

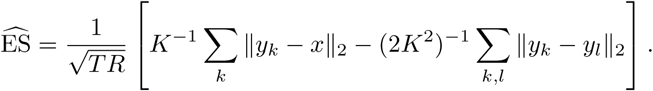

The temporal variogram score (Scheuerer & Hamill, 2015) compares target increment magnitudes with ensemble-expected increment magnitudes over the specified within-channel lags (Appendix C.5). Lower values are better. An empirical marginal reference consists of 18 other real target sequences selected by distinct nonzero offsets, equally spaced as integer indices from 1 to *N* 1, and added modulo *N* to the target index. A target never supplies its own reference, but reference sequences are reused between contexts. This is neither a deployable conditional predictor nor a mathematical lower bound.

Central-interval coverage and ranks among the 18 draws (19 possible ranks) pool dependent context/time/channel entries. No independent-sample confidence intervals are attached. The variance ratio divides ensemble variance by pooled target variance, which includes between-context differences; unity is not a conditional-calibration target. Structural composites use 18 Monte Carlo replicates, each taking one draw per corresponding context; points and sample SD quantify draw variability, not training-seed uncertainty. The within-session spread control multiplies centered deviations by 0.5, 1, 2, 4; energy and variogram values retain their own units and should only be compared within each metric.

**Conditioning control.** A separate ten-seed matched-capacity GRU experiment compares correct versus shuffled covariates: signed contrast, stimulus contrast, probability-left, previous choice and previous feedback. Current-trial choice, movement, feedback and reward are excluded. Complete covariate rows are shuffled within session and training/validation split; true held-out covariates are used for evaluation. The mean condition-stratified composite contrast is 0.000358 (paired 95% bootstrap interval [0.001922, 0.001317]; exact two-sided sign-flip *p* = 0.6914, Holm-adjusted *p* = 1; four of ten seeds positive). This does not establish a benefit for this structural endpoint, nor a general absence of behavioral or functional information. Population-transfer and order-control results, together with stochastic predictive calibration and the conditioning control, are shown in Figures S21 and S22, respectively.

### B.5 Two-photon calcium example

A separate single-mouse example uses two-photon calcium imaging through an implanted GRIN lens in a head-fixed mouse during exposure to appetitive rewards. The recording comprises calcium Δ*F/F* traces from 88 basolateral-amygdala principal neurons sampled at 10Hz for approximately 66 minutes, following CaImAn motion correction and source extraction (Giovannucci et al., 2019). These traces are not inferred spike counts. The population-conditioned MLP combines per-neuron temporal features and a population context embedding, with a residual output and learned persistence skip. The illustrated inference protocol applies one-step predictions autoregressively within chunks, then resets context to recorded activity (Figure S23). It therefore receives intermittent observations and is not comparable to a continuously autonomous rollout of the same total duration. The example illustrates a metric profile in another observation regime; it is not an independent multi-animal replication or a controlled architecture comparison with the widefield models.

### B.6 Brain Treebank contact-selection and retrieval audit

**Training-movie-only contact selection.** We repeated the original gross-QC algorithm using only the two training movies in each held-out fold. The BIDS-good intersection, pooled first-difference MAD thresholds, and evenly spaced selection were recomputed independently for each fold. Sparse raw-signal differences used exactly the original stride of 2048 samples, resolutions, and acquisition-order indices; they were retained for reproduction. As a check on the implementation, pooling all three movies exactly reproduced the original 64 selected contacts for every participant.

Training-only QC selected the same 64 contacts in 14 of 15 folds. For participant P01 with movie 02 held out, the retained pool changed from 154 to 153 contacts, and the evenly spaced 64-contact panels shared 46 contacts. The other 14 selections were identical. The main analysis therefore retains its original offline participant-level panel: these checks do not retroactively make its preprocessing test-blind. A complete refit with the changed contact panel is required to replace the affected fold. In particular, changing contacts also changes common-average referencing and the fitted normalization and models; deleting a few output channels is not an equivalent correction.

Excluding the changed-QC fold and reaveraging movies within participants preserves the main dissociation: the analog retrieval control has the highest composite (0.7034), and GRU multistep has the lowest MAE (0.8871). The GRU multistep-minus-one-step contrast is 0.0255, positive in all five participants. This is an omission sensitivity using existing forecasts, not a contact-panel refit.

**Programmatic retrieval exclusions.** For all 15 folds we verified that the held-out movie is absent from the two training-run identifiers, every candidate context/continuation lies wholly within a permitted training run, and every recorded selected candidate belongs to that pool. All 750 stored analog predictions exactly equal their selected 320-bin training continuation. The selector’s inputs are training runs, training quality masks, training indices, and observed contexts; neither the held-out future nor a held-out-movie library enters its PCA fit or nearest-context search. This audit checks stored matches, candidate bounds and the executed selector’s data contract; it does not substitute a new nearest-neighbor search or demonstrate prospective preprocessing.

**Offline observation transform.** The Butterworth filters are applied in both temporal directions and the Hilbert transform has nonlocal support. Consequently, a fixed finite onset exclusion cannot certify causal signal construction. We retain this limitation even when examining forecasts away from their boundary. Prospective claims require a one-sided transform, fold-specific contact selection and model refitting under that transform. The present experiment supports an offline comparison between realization accuracy and structural agreement, not a validated online forecasting system.

**Forecast-onset sensitivity on fixed predictions.** We rescored all 135 saved model/seed/fold arrays after excluding the first 32 forecast bins (2 s), leaving 50 sequences of 288 bins and 64 contacts per array. The same pinned component definitions and 16-Hz sampling rate were used. Native errors were recomputed on this identical 18-s support, then training seeds were averaged within movie, movies within participant, and the five participants equally. Analog retrieval remains the structural-composite leader (0.6936; MAE 1.2870), while GRU multistep retains the lowest MAE (0.8923; composite 0.3125). These are descriptive within-support comparisons; the values are not directly comparable to full 20-s scores. Removing onset samples does not remove nonlocal preprocessing dependence and is not a substitute for a causal-transform refit. The training-continuation provenance and context-stratified comparisons are shown in Figure S18; composite stability under sequence-safe directed relational rescoring is shown in Figure S19.

### B.7 Forecasting comparison across four datasets

**Forecasting task.** We fit each architecture on the four datasets described in Appendices B.1–B.4. Within each dataset, all models received the same neural history and predicted the same held-out activity. Table 6 gives the history, prediction block, and full forecast horizon. When the forecast extended beyond one block, we fed the predicted activity back as input to the next block. We scored only the forecast interval.

**Table 6:** **Forecast windows and test coverage.** Test counts are pooled across splits. The prediction block is the duration generated in one model call.

| Dataset | History | Prediction block | Forecast horizon | Held-out data |
|---|---|---|---|---|
| IBL | 0.5 s | 1.5 s | 1.5 s | 17 mice, 2,176 windows |
| Widefield | 3 s | 3 s | 24 s | 287 recordings, one window each |
| Brain Treebank | 5 s | 5 s | 20 s | 15 movie runs, 360 windows |
| OmniMouse | 1 s | 1 s | 2 s | 3 scans, 120 windows |

**Recordings and splits.** For IBL, we used regional firing rates sampled at 50 Hz and transformed as log(1 + *x*). For each held-out mouse, we scored the regions recorded in that mouse and represented in training. For widefield, we used the released 16-region traces at 30 Hz. We divided IBL mice into five groups and widefield recordings into six groups. Each fold used one group for testing, the next for validation, and the remaining groups for training. The widefield split was by recording, so recordings from the same mouse could occur in different folds.

For Brain Treebank, we used 10-minute crops of log high-gamma power (70–115 and 125–150 Hz) sampled at 16 Hz from 16 contacts marked good across all three runs of each participant. We fit models separately for each of the five participants, holding out each movie run in turn. The final 20% of each of the other two runs served as validation, with a 25 s gap between training and validation. For OmniMouse, we fit models separately within three naturalistic-video scans. We used a 10-minute crop per scan and selected 16 neurons using the training portion. We sampled responses at 30 Hz using the released acquisition offset for each neuron. Each scan was split chronologically into training, validation, and test portions (60/20/20), with a 3 s gap before each validation and test block.

We estimated normalization parameters separately for each channel from the observed histories of training windows. For Brain Treebank, we centered each channel by its median, divided by its scaled median absolute deviation, and clipped normalized values to [8, 8]. For the other three datasets, we used the training mean and standard deviation.

**Models and fitting.** Table 1 compares POCO, TSMixer, NetFormer, a two-layer GRU, and per-channel ridge regression (**Linear**). Tables 14 and 13 further report family and component scores.

**Table 7:** Principal widefield model configurations. *H* is the number of supervised future steps per training example. Teacher forcing supplies only preceding target observations; all evaluation rollouts use self-generated history. MLP widths list consecutive layer dimensions.

| Model | Architecture | Training inputs | *H* | Objective |
|---|---|---|---|---|
| VAR | Affine map, 1440 → 16 | Teacher forcing | 90 | MAE |
| RNN | GRU: 8 layers, width 64; layer-normalized readout | Teacher forcing | 90 | MAE |
| 1_step | Causal Transformer: width 64, 8 layers, 8 heads, FFN 128 | Context only | 1 | MAE |
| Sequifier | Causal Transformer: width 64, 8 layers, 8 heads, FFN 128 | Context only | 1 | MAE |
| TF | Same backbone as 1_step | Teacher forcing | 90 | MAE |
| AR | Same backbone as TF | Self-feedback | 90 | MAE |
| TF<sub>QL_KL</sub> | Same backbone as TF | Teacher forcing | 90 | MAE + histogram and quantile losses |
| SSM | Latent width 12; context/posterior GRUs 96; transition MLP 12→64→12; decoder 12→32→32→16 | Target-informed posterior (training only) | 90 | Regularized variational objective |

**Optimization and model selection.** On the same train-val-test partition, models were trained with seeds 101, 102, and 103. Shared AdamW (Loshchilov & Hutter, 2019) settings are collected in Table 8. For the reported widefield comparison, each run is represented by the checkpoint with the lowest validation mean-forecast MAE, independently of the training objective. For deterministic models this is the MAE of the model output; for the SSM it is the MAE after averaging decoder means over 16 sampled latent trajectories, with fixed validation seeds across epochs. Training-progress analyses instead use the specified intermediate checkpoints.

**Table 8:** Optimization settings for the principal widefield comparison. The neural-model column covers the Transformers, RNN, and SSM. Accumulation gives 2048 examples per full optimizer update. SSM-specific regularization is defined in Section D.4.

| Setting | Neural models | VAR |
|---|---|---|
| Optimizer | AdamW | AdamW |
| Learning rate | OneCycle, maximum 3 × 10<sup>−4</sup> | Constant 10<sup>−3</sup> |
| Schedule | Cosine; warm-up fraction 0.3; division factors 25 and 100 | None |
| Weight decay | 10<sup>−2</sup> | 10<sup>−5</sup> |
| Batch size / accumulation | 1024 / 2 | 2048 / 1 |
| Maximum epochs | 150 | 150 |
| Minimum validation improvement | 10<sup>−6</sup> | 10<sup>−6</sup> |
| Dropout | 0.05 (Transformers and RNN); none (SSM) | None |
| Gradient clipping / precision | Global norm 1; mixed precision on CUDA | No clipping; float32 |

**Table 9:** F**a**mily**-level Nethobench and pointwise-fidelity scores.** Model values are mean sample standard deviation across three training seeds, after averaging four evaluation-group scores within each seed. Empirical reference values are mean Monte Carlo sample SD across 20 random draws, after averaging four fixed evaluation groups of 56/56/55/55 sequences within each draw; all draws use the same 222 targets and 720-bin forecast window. Pointwise fidelity is computed on the same forecasts and evaluation groups as the structural scores and is reported separately from the structural composite.

|  | Model | Nethobench scores |  |  |  |  |  |  |
|---|---|---|---|---|---|---|---|---|
|  |  | D | T | R | G | S | Composite | Pointwise fidelity |
| Models | VAR | 0.345<br>± 0.005 | 0.342<br>± 0.008 | 0.614<br>± 0.014 | 0.757<br>± 0.020 | 0.340<br>± 0.006 | 0.482<br>± 0.006 | 0.354<br>± 0.004 |
|  | RNN | 0.425<br>± 0.010 | 0.466<br>± 0.011 | 0.496<br>± 0.012 | 0.760<br>± 0.012 | 0.613<br>± 0.018 | 0.544<br>± 0.006 | 0.357<br>± 0.003 |
|  | SSM | 0.502<br>± 0.020 | 0.502<br>± 0.013 | 0.562<br>± 0.014 | 0.811<br>± 0.024 | 0.668<br>± 0.028 | 0.602<br>± 0.018 | 0.309<br>± 0.003 |
|  | 1-step | 0.399<br>± 0.011 | 0.421<br>± 0.010 | 0.672<br>± 0.011 | 0.735<br>± 0.014 | 0.464<br>± 0.029 | 0.541<br>± 0.007 | 0.370<br>± 0.003 |
|  | Sequifier | 0.389<br>± 0.010 | 0.493<br>± 0.010 | 0.578<br>± 0.031 | 0.663<br>± 0.073 | 0.495<br>± 0.064 | 0.522<br>± 0.030 | 0.324<br>± 0.023 |
|  | AR | 0.437<br>± 0.007 | 0.403<br>± 0.014 | 0.604<br>± 0.024 | 0.714<br>± 0.016 | 0.500<br>± 0.005 | 0.532<br>± 0.009 | 0.363<br>± 0.003 |
|  | TF | 0.439<br>± 0.013 | 0.480<br>± 0.009 | 0.696<br>± 0.006 | 0.714<br>± 0.009 | 0.541<br>± 0.012 | 0.576<br>± 0.006 | 0.364<br>± 0.001 |
|  | TFQLKL | 0.438<br>± 0.010 | 0.473<br>± 0.007 | 0.679<br>± 0.009 | 0.678<br>± 0.028 | 0.544<br>± 0.018 | 0.564<br>± 0.009 | 0.364<br>± 0.002 |
| Refs. | Real–real reference | 0.721<br>± 0.038 | 0.843<br>± 0.013 | 0.830<br>± 0.012 | 0.893<br>± 0.041 | 0.888<br>± 0.017 | 0.830<br>± 0.012 | – |
|  | Time+region shuffle | 0.562<br>± 0.021 | 0.278<br>± 0.008 | 0.396<br>± 0.013 | 0.440<br>± 0.037 | 0.503<br>± 0.039 | 0.439<br>± 0.007 | – |

**Table 10:**
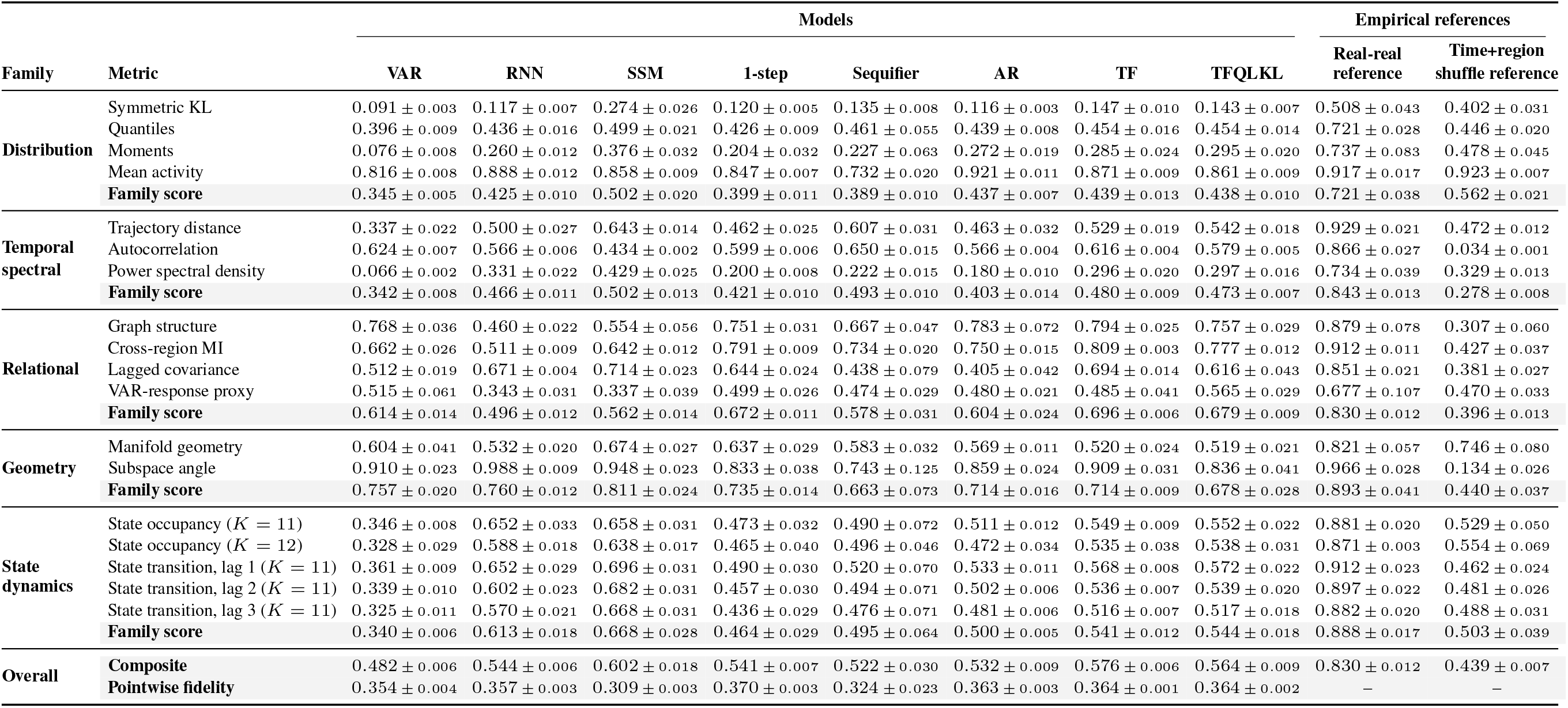
M**e**tric**-level Nethobench scores by family and model.** Model values are mean sample standard deviation across three training seeds, after averaging four evaluation-group scores within each seed. Empirical reference values are mean Monte Carlo sample SD across 20 random draws, after averaging four fixed evaluation groups of 56/56/55/55 sequences within each draw; all draws use the same 222 targets and 720-bin forecast window. Pointwise fidelity is computed on the same forecasts and evaluation groups as the structural scores and is reported separately from the structural composite.

| **Family** | **Metric** | **Models** |  |  |  |  |  |  |  | **Empirical references** |  |
|---|---|---|---|---|---|---|---|---|---|---|---|
|  |  | **VAR** | **RNN** | **SSM** | **1-step** | **Sequifier** | **AR** | **TF** | **TFQLKL** | **Real-real reference** | **Time+region shuffle reference** |
| **Distribution** | Symmetric KL | 0.091 ± 0.003 | 0.117 ± 0.007 | 0.274 ± 0.026 | 0.120 ± 0.005 | 0.135 ± 0.008 | 0.116 ± 0.003 | 0.147 ± 0.010 | 0.143 ± 0.007 | 0.508 ± 0.043 | 0.402 ± 0.031 |
|  | Quantiles | 0.396 ± 0.009 | 0.436 ± 0.016 | 0.499 ± 0.021 | 0.426 ± 0.009 | 0.461 ± 0.055 | 0.439 ± 0.008 | 0.454 ± 0.016 | 0.454 ± 0.014 | 0.721 ± 0.028 | 0.446 ± 0.020 |
|  | Moments | 0.076 ± 0.008 | 0.260 ± 0.012 | 0.376 ± 0.032 | 0.204 ± 0.032 | 0.227 ± 0.063 | 0.272 ± 0.019 | 0.285 ± 0.024 | 0.295 ± 0.020 | 0.737 ± 0.083 | 0.478 ± 0.045 |
|  | Mean activity | 0.816 ± 0.008 | 0.888 ± 0.012 | 0.858 ± 0.009 | 0.847 ± 0.007 | 0.732 ± 0.020 | 0.921 ± 0.011 | 0.871 ± 0.009 | 0.861 ± 0.009 | 0.917 ± 0.017 | 0.923 ± 0.007 |
|  | **Family score** | 0.345 ± 0.005 | 0.425 ± 0.010 | 0.502 ± 0.020 | 0.399 ± 0.011 | 0.389 ± 0.010 | 0.437 ± 0.007 | 0.439 ± 0.013 | 0.438 ± 0.010 | 0.721 ± 0.038 | 0.562 ± 0.021 |
| **Temporal spectral** | Trajectory distance | 0.337 ± 0.022 | 0.500 ± 0.027 | 0.643 ± 0.014 | 0.462 ± 0.025 | 0.607 ± 0.031 | 0.463 ± 0.032 | 0.529 ± 0.019 | 0.542 ± 0.018 | 0.929 ± 0.021 | 0.472 ± 0.012 |
|  | Autocorrelation | 0.624 ± 0.007 | 0.566 ± 0.006 | 0.434 ± 0.002 | 0.599 ± 0.006 | 0.650 ± 0.015 | 0.566 ± 0.004 | 0.616 ± 0.004 | 0.579 ± 0.005 | 0.866 ± 0.027 | 0.034 ± 0.001 |
|  | Power spectral density | 0.066 ± 0.002 | 0.331 ± 0.022 | 0.429 ± 0.025 | 0.200 ± 0.008 | 0.222 ± 0.015 | 0.180 ± 0.010 | 0.296 ± 0.020 | 0.297 ± 0.016 | 0.734 ± 0.039 | 0.329 ± 0.013 |
|  | **Family score** | 0.342 ± 0.008 | 0.466 ± 0.011 | 0.502 ± 0.013 | 0.421 ± 0.010 | 0.493 ± 0.010 | 0.403 ± 0.014 | 0.480 ± 0.009 | 0.473 ± 0.007 | 0.843 ± 0.013 | 0.278 ± 0.008 |
| **Relational** | Graph structure | 0.768 ± 0.036 | 0.460 ± 0.022 | 0.554 ± 0.056 | 0.751 ± 0.031 | 0.667 ± 0.047 | 0.783 ± 0.072 | 0.794 ± 0.025 | 0.757 ± 0.029 | 0.879 ± 0.078 | 0.307 ± 0.060 |
|  | Cross-region MI | 0.662 ± 0.026 | 0.511 ± 0.009 | 0.642 ± 0.012 | 0.791 ± 0.009 | 0.734 ± 0.020 | 0.750 ± 0.015 | 0.809 ± 0.003 | 0.777 ± 0.012 | 0.912 ± 0.011 | 0.427 ± 0.037 |
|  | Lagged covariance | 0.512 ± 0.019 | 0.671 ± 0.004 | 0.714 ± 0.023 | 0.644 ± 0.024 | 0.438 ± 0.079 | 0.405 ± 0.042 | 0.694 ± 0.014 | 0.616 ± 0.043 | 0.851 ± 0.021 | 0.381 ± 0.027 |
|  | VAR-response proxy | 0.515 ± 0.061 | 0.343 ± 0.031 | 0.337 ± 0.039 | 0.499 ± 0.026 | 0.474 ± 0.029 | 0.480 ± 0.021 | 0.485 ± 0.041 | 0.565 ± 0.029 | 0.677 ± 0.107 | 0.470 ± 0.033 |
|  | **Family score** | 0.614 ± 0.014 | 0.496 ± 0.012 | 0.562 ± 0.014 | 0.672 ± 0.011 | 0.578 ± 0.031 | 0.604 ± 0.024 | 0.696 ± 0.006 | 0.679 ± 0.009 | 0.830 ± 0.012 | 0.396 ± 0.013 |
| **Geometry** | Manifold geometry | 0.604 ± 0.041 | 0.532 ± 0.020 | 0.674 ± 0.027 | 0.637 ± 0.029 | 0.583 ± 0.032 | 0.569 ± 0.011 | 0.520 ± 0.024 | 0.519 ± 0.021 | 0.821 ± 0.057 | 0.746 ± 0.080 |
|  | Subspace angle | 0.910 ± 0.023 | 0.988 ± 0.009 | 0.948 ± 0.023 | 0.833 ± 0.038 | 0.743 ± 0.125 | 0.859 ± 0.024 | 0.909 ± 0.031 | 0.836 ± 0.041 | 0.966 ± 0.028 | 0.134 ± 0.026 |
|  | **Family score** | 0.757 ± 0.020 | 0.760 ± 0.012 | 0.811 ± 0.024 | 0.735 ± 0.014 | 0.663 ± 0.073 | 0.714 ± 0.016 | 0.714 ± 0.009 | 0.678 ± 0.028 | 0.893 ± 0.041 | 0.440 ± 0.037 |
| **State dynamics** | State occupancy (*K* = 11) | 0.346 ± 0.008 | 0.652 ± 0.033 | 0.658 ± 0.031 | 0.473 ± 0.032 | 0.490 ± 0.072 | 0.511 ± 0.012 | 0.549 ± 0.009 | 0.552 ± 0.022 | 0.881 ± 0.020 | 0.529 ± 0.050 |
|  | State occupancy (*K* = 12) | 0.328 ± 0.029 | 0.588 ± 0.018 | 0.638 ± 0.017 | 0.465 ± 0.040 | 0.496 ± 0.046 | 0.472 ± 0.034 | 0.535 ± 0.038 | 0.538 ± 0.031 | 0.871 ± 0.003 | 0.554 ± 0.069 |
|  | State transition, lag 1 (*K* = 11) | 0.361 ± 0.009 | 0.652 ± 0.029 | 0.696 ± 0.031 | 0.490 ± 0.030 | 0.520 ± 0.070 | 0.533 ± 0.011 | 0.568 ± 0.008 | 0.572 ± 0.022 | 0.912 ± 0.023 | 0.462 ± 0.024 |
|  | State transition, lag 2 (*K* = 11) | 0.339 ± 0.010 | 0.602 ± 0.023 | 0.682 ± 0.031 | 0.457 ± 0.030 | 0.494 ± 0.071 | 0.502 ± 0.006 | 0.536 ± 0.007 | 0.539 ± 0.020 | 0.897 ± 0.022 | 0.481 ± 0.026 |
|  | State transition, lag 3 (*K* = 11) | 0.325 ± 0.011 | 0.570 ± 0.021 | 0.668 ± 0.031 | 0.436 ± 0.029 | 0.476 ± 0.071 | 0.481 ± 0.006 | 0.516 ± 0.007 | 0.517 ± 0.018 | 0.882 ± 0.020 | 0.488 ± 0.031 |
|  | **Family score** | 0.340 ± 0.006 | 0.613 ± 0.018 | 0.668 ± 0.028 | 0.464 ± 0.029 | 0.495 ± 0.064 | 0.500 ± 0.005 | 0.541 ± 0.012 | 0.544 ± 0.018 | 0.888 ± 0.017 | 0.503 ± 0.039 |
| **Overall** | **Composite** | 0.482 ± 0.006 | 0.544 ± 0.006 | 0.602 ± 0.018 | 0.541 ± 0.007 | 0.522 ± 0.030 | 0.532 ± 0.009 | 0.576 ± 0.006 | 0.564 ± 0.009 | 0.830 ± 0.012 | 0.439 ± 0.007 |
|  | **Pointwise fidelity** | 0.354 ± 0.004 | 0.357 ± 0.003 | 0.309 ± 0.003 | 0.370 ± 0.003 | 0.324 ± 0.023 | 0.363 ± 0.003 | 0.364 ± 0.001 | 0.364 ± 0.002 | – | – |

**Table 11:**
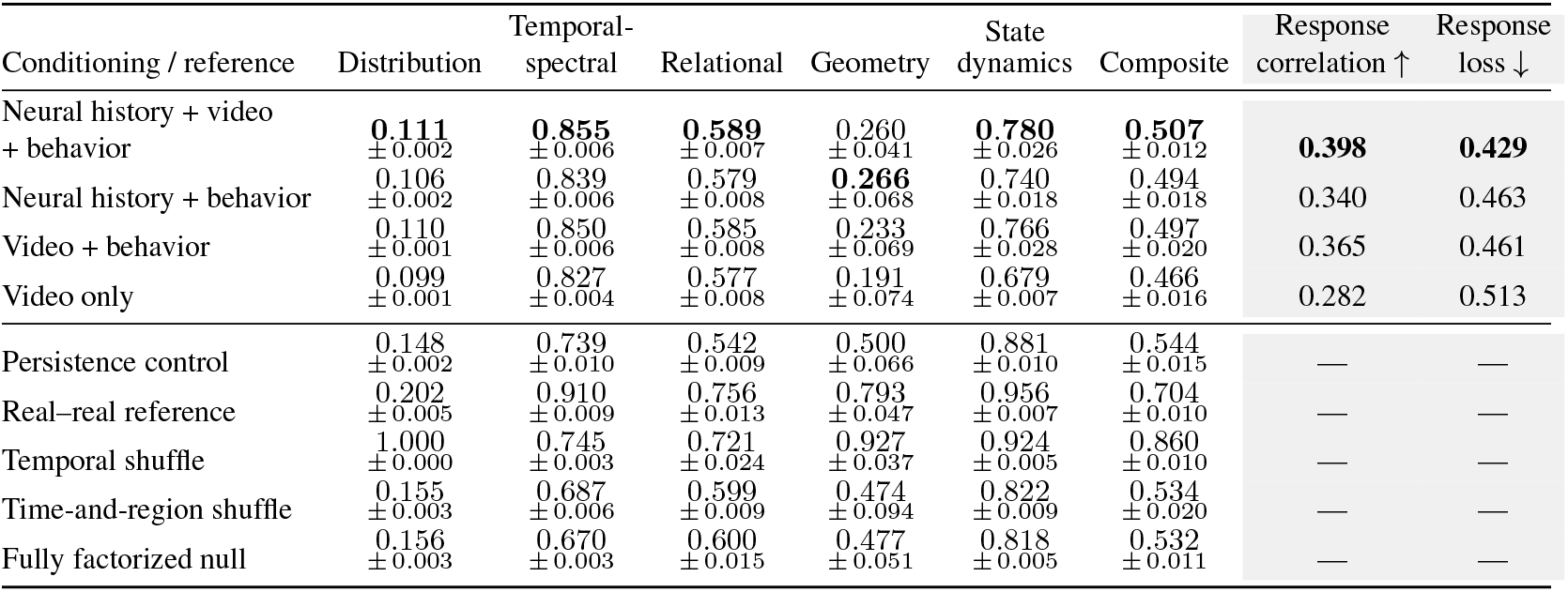
O**m**niMouse **evaluation at the native horizon with shared context.** Nethobench compares 30 recorded context samples followed by 30 predicted samples against the same context followed by the recorded future. Nethobench values are mean SEM across four sequence-level splits from one held-out session. The two grey columns reproduce response correlation (higher is better) and response loss (lower is better) reported by the OmniMouse authors under their evaluation protocol. Bold indicates the best mean or reported value among the four conditioning configurations in each column, excluding controls and references. The shared context contributes to absolute agreement. Temporal shuffling is reported as a control, not a universal score floor.

**Table 12:** O**m**niMouse **evaluation on two-block forecast-only sequences.** Nethobench compares two successively generated 30-sample blocks (2 s at 30 Hz) against the corresponding recorded future, excluding the initial context from scoring. Enabled video and behavioral covariates remain recorded observations spanning both predicted intervals. Values are mean SEM across four sequence-level splits from one held-out session. Bold indicates the highest mean among the four conditioning configurations in each column, excluding references and controls. Temporal shuffling is reported as a control, not a universal score floor.

| Conditioning / reference | Distribution | Temporal-spectral | Relational | Geometry | State dynamics | Composite |
|---|---|---|---|---|---|---|
| Neural history + video + behavior | 0.062<br>± 0.002 | 0.696<br>± 0.007 | 0.494<br>± 0.006 | **0.126**<br>± 0.063 | **0.669**<br>± 0.055 | 0.400<br>± 0.008 |
| Neural history + behavior | 0.056<br>± 0.002 | 0.662<br>± 0.006 | 0.473<br>± 0.002 | 0.085<br>± 0.031 | 0.649<br>± 0.058 | 0.377<br>± 0.007 |
| Video + behavior | **0.064**<br>± 0.001 | **0.716**<br>± 0.003 | **0.499**<br>± 0.008 | 0.122<br>± 0.061 | 0.645<br>± 0.059 | **0.401**<br>± 0.008 |
| Video only | 0.057<br>± 0.001 | 0.666<br>± 0.012 | 0.482<br>± 0.003 | 0.076<br>± 0.018 | 0.496<br>± 0.058 | 0.351<br>± 0.011 |
| Real–real reference | 0.144<br>± 0.005 | 0.904<br>± 0.014 | 0.643<br>± 0.029 | 0.591<br>± 0.034 | 0.917<br>± 0.019 | 0.620<br>± 0.017 |
| Temporal shuffle | 1.000<br>± 0.000 | 0.630<br>± 0.005 | 0.619<br>± 0.016 | 0.873<br>± 0.044 | 0.861<br>± 0.010 | 0.794<br>± 0.009 |
| Time-and-region shuffle | 0.107<br>± 0.002 | 0.513<br>± 0.014 | 0.441<br>± 0.002 | 0.041<br>± 0.037 | 0.649<br>± 0.017 | 0.346<br>± 0.008 |
| Fully factorized null | 0.108<br>± 0.002 | 0.504<br>± 0.018 | 0.440<br>± 0.001 | 0.029<br>± 0.022 | 0.652<br>± 0.012 | 0.343<br>± 0.007 |

**Table 13:** P**o**intwise **error and structural agreement by family.** Within each dataset, fitted models are ordered by increasing test MAE. Linear denotes per-channel ridge. Bold marks the best fitted value in each score column, using unrounded means. D, T, R, G, and S denote distributional, temporal– spectral, relational, geometric, and state-dynamical agreement, respectively. RR, WC, and TR denote the real–real, within-channel shuffle, and time-and-region shuffle references, respectively. Further experimental details are provided in Appendix B.7. Structural agreement by family *↑* Test score

|  | Structural agreement by family ↑ |  |  |  |  | Test score |  |
|---|---|---|---|---|---|---|---|
| Model | D | T | R | G | S | MAE ↓ | Comp. ↑ |
| **IBL** |  |  |  |  |  |  |  |
| POCO | 0.289 | 0.320 | 0.599 | 0.720 | 0.366 | **0.6860** | 0.4601 |
| Linear | **0.290** | **0.342** | **0.607** | **0.720** | **0.384** | 0.7102 | **0.4698** |
| TSMixer | 0.265 | 0.334 | 0.535 | 0.666 | 0.355 | 0.7812 | 0.4305 |
| GRU | 0.250 | 0.334 | 0.500 | 0.642 | 0.323 | 0.8260 | 0.4089 |
| NetFormer | 0.215 | 0.192 | 0.382 | 0.482 | 0.267 | 1.1300 | 0.3084 |
| RR | 0.582 | 0.591 | 0.773 | 0.862 | 0.885 | 0.7980 | 0.7344 |
| WC | 1.000 | 0.620 | 0.717 | 0.897 | 0.901 | 0.7168 | 0.8274 |
| TR | 0.272 | 0.412 | 0.549 | 0.666 | 0.457 | 1.2021 | 0.4679 |
| **Widefield** |  |  |  |  |  |  |  |
| TSMixer | 0.381 | 0.309 | **0.641** | 0.716 | 0.366 | **0.7207** | 0.4880 |
| Linear | 0.367 | **0.324** | 0.602 | 0.667 | 0.351 | 0.7212 | 0.4668 |
| GRU | 0.381 | 0.302 | 0.559 | **0.749** | 0.370 | 0.7214 | 0.4737 |
| POCO | **0.384** | 0.314 | 0.630 | 0.744 | **0.376** | 0.7215 | **0.4938** |
| NetFormer | 0.383 | 0.311 | 0.369 | 0.692 | 0.368 | 0.7480 | 0.4195 |
| RR | 0.560 | 0.732 | 0.752 | 0.866 | 0.901 | 1.0650 | 0.7537 |
| WC | 1.000 | 0.351 | 0.421 | 0.508 | 0.513 | 1.0073 | 0.5680 |
| TR | 0.648 | 0.311 | 0.397 | 0.467 | 0.516 | 1.0391 | 0.4708 |
| **Brain Treebank** |  |  |  |  |  |  |  |
| GRU | **0.342** | 0.304 | 0.403 | 0.570 | 0.342 | **0.8195** | 0.3907 |
| POCO | 0.340 | **0.319** | 0.402 | **0.653** | **0.387** | 0.8316 | **0.4157** |
| Linear | 0.330 | 0.294 | **0.441** | 0.579 | 0.309 | 0.8680 | 0.3913 |
| TSMixer | 0.326 | 0.308 | 0.405 | 0.604 | 0.344 | 0.8939 | 0.3951 |
| NetFormer | 0.311 | 0.220 | 0.298 | 0.618 | 0.320 | 1.7738 | 0.3486 |
| RR | 0.595 | 0.667 | 0.672 | 0.773 | 0.863 | 1.1137 | 0.7070 |
| WC | 1.000 | 0.665 | 0.530 | 0.718 | 0.773 | 1.0509 | 0.7351 |
| TR | 0.666 | 0.643 | 0.419 | 0.603 | 0.732 | 1.0768 | 0.6031 |
| **OmniMouse** |  |  |  |  |  |  |  |
| Linear | **0.064** | 0.355 | **0.454** | **0.574** | 0.384 | **0.3095** | 0.3596 |
| TSMixer | 0.059 | 0.347 | 0.423 | 0.474 | 0.416 | 0.3313 | 0.3372 |
| POCO | 0.059 | **0.371** | 0.416 | 0.467 | **0.575** | 0.3428 | **0.3672** |
| GRU | 0.054 | 0.351 | 0.377 | 0.390 | 0.463 | 0.3587 | 0.3192 |
| NetFormer | 0.057 | 0.328 | 0.368 | 0.427 | 0.539 | 0.6036 | 0.3335 |
| RR | 0.115 | 0.486 | 0.514 | 0.634 | 0.868 | 0.4442 | 0.5065 |
| WC | 1.000 | 0.424 | 0.581 | 0.849 | 0.872 | 0.3140 | 0.7456 |
| TR | 0.069 | 0.242 | 0.409 | 0.489 | 0.604 | 0.4283 | 0.3537 |

**Table 14:** **Component-level structural agreement across four datasets.** Higher scores indicate closer agreement. Bold marks the highest available mean among the five fitted models for each component and dataset, before rounding. Dashes denote unavailable scores. Linear denotes per-channel ridge. RR, WC, and TR denote the real–real, within-channel shuffle, and time-and-region shuffle references, respectively. Transition scores use *K* = 11. Components are grouped by family in the order used in Table 13. Further experimental details are provided in Appendix B.7.

| Component | POCO | TSMixer | NetFormer | Linear | GRU | RR | WC | TR |
|---|---|---|---|---|---|---|---|---|
| **IBL** |  |  |  |  |  |  |  |  |
| Symmetric KL | 0.040 | 0.040 | **0.047** | 0.041 | 0.041 | 0.378 | 1.000 | 0.093 |
| Tail quantiles | — | — | — | — | — | — | — | — |
| Moments | 0.261 | 0.255 | 0.240 | **0.267** | 0.250 | 0.775 | 1.000 | 0.337 |
| Mean shift | **0.566** | 0.501 | 0.358 | 0.562 | 0.460 | 0.592 | 1.000 | 0.387 |
| Trajectory | 0.625 | 0.627 | 0.295 | **0.640** | 0.625 | 0.962 | 0.941 | 0.668 |
| Autocorrelation | 0.231 | 0.220 | 0.127 | **0.237** | 0.231 | 0.234 | 0.205 | 0.201 |
| Power spectrum | 0.105 | 0.154 | **0.156** | 0.149 | 0.147 | 0.576 | 0.712 | 0.368 |
| Correlation graph | 0.717 | 0.614 | 0.450 | **0.725** | 0.625 | 0.865 | 0.752 | 0.670 |
| Cross-region MI | 0.474 | 0.434 | 0.275 | **0.475** | 0.363 | 0.654 | 0.501 | 0.467 |
| Lagged covariance | 0.570 | 0.500 | 0.324 | **0.586** | 0.455 | 0.759 | 0.728 | 0.467 |
| VAR response proxy | 0.568 | 0.526 | 0.394 | **0.573** | 0.485 | 0.767 | 0.836 | 0.500 |
| Manifold geometry | 0.573 | 0.559 | 0.507 | **0.601** | 0.547 | 0.816 | 0.810 | 0.530 |
| Subspace angle | **0.866** | 0.774 | 0.457 | 0.840 | 0.737 | 0.908 | 0.984 | 0.803 |
| Occupancy (*K* = 11) | 0.444 | 0.420 | 0.311 | **0.455** | 0.380 | 0.905 | 0.952 | 0.478 |
| Occupancy (*K* = 12) | 0.402 | 0.376 | 0.297 | **0.427** | 0.346 | 0.906 | 0.952 | 0.476 |
| Transitions (lag 1) | 0.332 | 0.331 | 0.246 | **0.350** | 0.301 | 0.874 | 0.845 | 0.446 |
| Transitions (lag 2) | 0.328 | 0.327 | 0.244 | **0.346** | 0.296 | 0.871 | 0.870 | 0.444 |
| Transitions (lag 3) | 0.322 | 0.320 | 0.238 | **0.340** | 0.291 | 0.868 | 0.888 | 0.441 |
| **Widefield** |  |  |  |  |  |  |  |  |
| Symmetric KL | **0.097** | 0.094 | 0.080 | 0.084 | 0.095 | 0.300 | 1.000 | 0.501 |
| Tail quantiles | **0.406** | 0.401 | 0.392 | 0.390 | 0.396 | 0.462 | 1.000 | 0.603 |
| Moments | 0.089 | 0.084 | **0.141** | 0.052 | 0.098 | 0.568 | 1.000 | 0.559 |
| Mean shift | **0.945** | 0.945 | 0.920 | 0.944 | 0.933 | 0.911 | 1.000 | 0.930 |
| Trajectory | 0.391 | **0.417** | 0.405 | 0.314 | 0.377 | 0.932 | 0.512 | 0.502 |
| Autocorrelation | 0.448 | 0.425 | 0.393 | **0.610** | 0.439 | 0.774 | 0.034 | 0.034 |
| Power spectrum | 0.102 | 0.085 | **0.134** | 0.047 | 0.090 | 0.491 | 0.507 | 0.395 |
| Correlation graph | 0.812 | 0.828 | 0.386 | **0.834** | 0.714 | 0.795 | 0.304 | 0.281 |
| Cross-region MI | 0.705 | **0.755** | 0.415 | 0.723 | 0.723 | 0.894 | 0.522 | 0.443 |
| Lagged covariance | 0.490 | **0.501** | 0.350 | 0.425 | 0.465 | 0.743 | 0.409 | 0.390 |
| VAR response proxy | **0.512** | 0.480 | 0.326 | 0.425 | 0.332 | 0.578 | 0.448 | 0.474 |
| Manifold geometry | **0.592** | 0.579 | 0.406 | 0.592 | 0.591 | 0.776 | 0.671 | 0.679 |
| Subspace angle | 0.896 | 0.852 | **0.977** | 0.741 | 0.907 | 0.956 | 0.345 | 0.255 |
| Occupancy (*K* = 11) | **0.386** | 0.376 | 0.376 | 0.363 | 0.382 | 0.895 | 0.554 | 0.561 |
| Occupancy (*K* = 12) | **0.368** | 0.360 | 0.361 | 0.344 | 0.362 | 0.883 | 0.540 | 0.541 |
| Transitions (lag 1) | **0.396** | 0.386 | 0.387 | 0.370 | 0.390 | 0.918 | 0.486 | 0.488 |
| Transitions (lag 2) | **0.372** | 0.362 | 0.365 | 0.347 | 0.367 | 0.908 | 0.493 | 0.496 |
| Transitions (lag 3) | **0.356** | 0.346 | 0.350 | 0.330 | 0.350 | 0.900 | 0.492 | 0.496 |
| **Brain Treebank** |  |  |  |  |  |  |  |  |
| Symmetric KL | 0.091 | 0.089 | 0.091 | 0.090 | **0.101** | 0.502 | 1.000 | 0.598 |
| Tail quantiles | 0.426 | 0.407 | 0.371 | 0.414 | **0.433** | 0.598 | 1.000 | 0.658 |
| Moments | 0.126 | 0.121 | **0.184** | 0.129 | 0.122 | 0.587 | 1.000 | 0.618 |
| Mean shift | **0.716** | 0.688 | 0.598 | 0.687 | 0.710 | 0.694 | 1.000 | 0.790 |
| Trajectory | **0.486** | 0.458 | 0.310 | 0.456 | 0.482 | 0.909 | 0.860 | 0.858 |
| Autocorrelation | **0.403** | 0.399 | 0.146 | 0.375 | 0.368 | 0.531 | 0.462 | 0.459 |
| Power spectrum | 0.069 | 0.068 | **0.205** | 0.051 | 0.063 | 0.564 | 0.672 | 0.613 |
| Correlation graph | 0.350 | 0.349 | 0.237 | **0.499** | 0.373 | 0.733 | 0.450 | 0.304 |
| Cross-region MI | 0.387 | 0.386 | 0.319 | 0.339 | **0.397** | 0.660 | 0.482 | 0.463 |
| Lagged covariance | 0.441 | 0.425 | 0.321 | **0.461** | 0.426 | 0.659 | 0.601 | 0.441 |
| VAR response proxy | 0.429 | 0.458 | 0.316 | **0.467** | 0.416 | 0.638 | 0.586 | 0.468 |
| Manifold geometry | 0.508 | 0.485 | 0.433 | **0.523** | 0.456 | 0.740 | 0.717 | 0.722 |
| Subspace angle | 0.798 | 0.724 | **0.804** | 0.635 | 0.683 | 0.806 | 0.719 | 0.484 |
| Occupancy (*K* = 11) | **0.459** | 0.404 | 0.378 | 0.355 | 0.405 | 0.882 | 0.807 | 0.768 |
| Occupancy (*K* = 12) | **0.447** | 0.377 | 0.384 | 0.356 | 0.400 | 0.880 | 0.797 | 0.752 |
| Transitions (lag 1) | **0.347** | 0.315 | 0.277 | 0.280 | 0.303 | 0.853 | 0.746 | 0.708 |
| Transitions (lag 2) | **0.341** | 0.312 | 0.280 | 0.276 | 0.301 | 0.850 | 0.755 | 0.716 |
| Transitions (lag 3) | **0.342** | 0.312 | 0.282 | 0.277 | 0.300 | 0.851 | 0.760 | 0.715 |
| **OmniMouse** |  |  |  |  |  |  |  |  |
| Symmetric KL | 0.045 | 0.040 | **0.060** | 0.040 | 0.044 | 0.152 | 1.000 | 0.077 |
| Tail quantiles | — | — | — | — | — | — | — | — |
| Moments | 0.051 | 0.057 | 0.043 | **0.071** | 0.038 | 0.141 | 1.000 | 0.044 |
| Mean shift | **0.083** | 0.080 | 0.067 | 0.081 | 0.078 | 0.052 | 1.000 | 0.086 |
| Trajectory | **0.683** | 0.659 | 0.528 | 0.583 | 0.661 | 0.815 | 0.679 | 0.569 |
| Autocorrelation | 0.362 | 0.320 | 0.380 | **0.431** | 0.325 | 0.481 | 0.104 | 0.097 |
| Power spectrum | 0.068 | 0.063 | **0.076** | 0.052 | 0.067 | 0.162 | 0.489 | 0.062 |
| Correlation graph | 0.225 | 0.390 | 0.182 | **0.399** | 0.264 | 0.351 | 0.538 | 0.319 |
| Cross-region MI | **0.365** | 0.307 | 0.320 | 0.355 | 0.336 | 0.655 | 0.581 | 0.340 |
| Lagged covariance | 0.615 | 0.548 | 0.508 | **0.641** | 0.480 | 0.559 | 0.766 | 0.498 |
| VAR response proxy | 0.459 | 0.449 | **0.461** | 0.422 | 0.429 | 0.492 | 0.441 | 0.479 |
| Manifold geometry | 0.670 | 0.611 | **0.680** | 0.653 | 0.610 | 0.574 | 0.733 | 0.619 |
| Subspace angle | 0.265 | 0.336 | 0.173 | **0.496** | 0.171 | 0.694 | 0.965 | 0.358 |
| Occupancy (*K* = 11) | **0.612** | 0.415 | 0.572 | 0.385 | 0.467 | 0.870 | 0.936 | 0.661 |
| Occupancy (*K* = 12) | 0.485 | 0.427 | 0.476 | 0.383 | **0.498** | 0.871 | 0.927 | 0.623 |
| Transitions (lag 1) | **0.607** | 0.429 | 0.572 | 0.396 | 0.468 | 0.873 | 0.801 | 0.566 |
| Transitions (lag 2) | **0.590** | 0.413 | 0.547 | 0.384 | 0.451 | 0.866 | 0.836 | 0.582 |
| Transitions (lag 3) | **0.579** | 0.396 | 0.527 | 0.372 | 0.433 | 0.860 | 0.859 | 0.590 |

We trained each of the four neural models with three seeds per split, minimizing mean squared error over observed channels with AdamW and batches of 64. For each learning rate (10^−3^ or 3 10^−4^), we ran 400 updates on IBL, 1,200 on widefield, and 600 on both Brain Treebank and OmniMouse. We evaluated validation forecasts every 40, 150, and 100 updates, respectively. For the linear model, we tested penalties of 0.01, 0.1, and 1 and tested histories of 5, 10, 20, and 45 samples whenever they fit within the observed window. We selected neural checkpoints and ridge settings by the lowest validation RMSE over the full rollout.

**Scoring and ranking.** We evaluated the selected fits by test MAE and by the Nethobench composite, using the family definitions and component-availability rules in Appendix C.1. We first averaged scores over training seeds for the neural models. Dataset means gave equal weight to IBL mice, widefield folds, Brain Treebank participants, or OmniMouse scans. For Brain Treebank, we first averaged the three held-out runs within each participant. Within each dataset, we ranked the five models in Table 1 separately by increasing mean MAE and decreasing mean composite. For the paired IBL comparison, we counted mice for which POCO had lower MAE and Linear had higher composite on the same held-out windows.

## C Validation and robustness analyses

### C.1 Metric definitions and implementation

**Inputs and aggregation.** Reference and prediction tensors must be aligned to identical sequence, time and channel keys, with shape *M T R*. Unaligned samples are not silently truncated. Different components compare different summaries: some pool time and sequences, others first score individual sequences or channels. Let *s_j_* [0, 1] be a finite component similarity. Each family is the arithmetic mean of its available components, and the five families receive the fixed weights in Equation 1. There are four distributional, three temporal-spectral, four relational, two geometric and five state dynamics components. If a component is undefined, its within-family weight is redistributed over the finite components. If an entire family is absent, outer weights are likewise renormalized. Missing values are not zeros and component availability must accompany a result.

The implementation distinguishes a similarity from its associated raw discrepancy. Frequently used transforms are *q*(*d*) = (1 + *d*)^−1^ for *d* 0 and *c*(*a, b*) = clip((1 + corr(*a, b*))*/*2, 0, 1). The latter is an affine, clipped Pearson score, not raw signed correlation or absolute correlation. For vectors, *r*(*a, b*) = *q*(RMSE(*a, b*)*/σ_a_*), falling back from reference SD to mean absolute reference magnitude and then 1 for degenerate scales. A generic matrix comparison applies 0.6*c* + 0.4*r* to strictly upper-triangular entries, with available-term renormalization. These choices define numerical agreement rather than an identified generative likelihood.

**Distributional components.** *Symmetric KL:* shared histogram support is derived from pooled reference and prediction 0.001/0.999 quantiles, with 60 interior bins and two tail bins. Histograms use additive 10^−12^ stabilization. For each sequence/channel pair with at least ten finite samples,

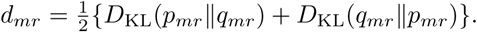

Within a sequence, *q*(*d_mr_*) values are geometrically averaged over channels. The final score is half their sequence mean plus half their tenth percentile. The retained divergence is symmetric KL; the generic implementation label “KL/JSD” does not mean that JSD was used.

*Mean shift:* absolute reference–prediction temporal mean differences are divided by the pooled reference channel IQR. The largest 0.1*R* channel discrepancies are averaged per sequence, then across sequences, before applying *q*. *Tail quantiles:* 99 quantiles at levels 0.01–0.99 are computed, retaining levels at or below 0.10 and at or above 0.90. IQR-scaled absolute differences are averaged over retained levels, then over the largest 0.25*R* channel discrepancies and sequences. The score is *q* of this distance. At least 80 samples per sequence/channel are required; long sequences can be subsampled to 1200 samples with fixed RNG seed 0.

*Moments:* pooled per-channel variance, skewness and excess kurtosis yield

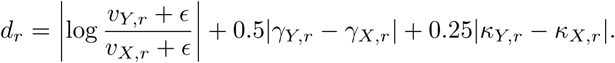

The component averages *q*(*d_r_*) over finite channels, requiring at least 24 samples. Skewness and kurtosis use finite-sample bias correction. The retained pooled-moment score should not be confused with auxiliary weighted-moment visualizations or differentiable training losses.

**Temporal-spectral components.** The temporal family is the equal mean of trajectory, ACF and power-sensitive PSD agreement. *Trajectory* combines occupancy/velocity comparisons (weight 0.6) and path-feature comparisons (0.4) in reference-fitted PCA coordinates. Occupancy uses 12-bin axis histograms; speed and turning are compared through quantiles at 0.1, 0.3, 0.5, 0.7 and 0.9. Path summaries include path length, displacement, radius, displacement-to-path ratio and lag-one speed dependence. The PCA variance target is 85%, with component caps of three for occupancy and four for path features. These are selected trajectory statistics, not an exact path-distribution distance.

*ACF* compares both raw activity and first increments using relative *ℓ*_2_ discrepancies of autocorrelation vectors. Physical delays correspond to 1, 2, 4, 8, 16, 32, 48 */*30 seconds, converted to rounded, unique valid sample lags using each dataset’s sampling frequency. Constant signals can make this component undefined. *PSD* uses within-sequence Welch density estimates with constant detrending and at most 256 samples per segment, excluding DC. With normalized non-DC spectra *p, q* and total powers *P_X_, P_Y_*, its agreement combines spectral overlap and power:

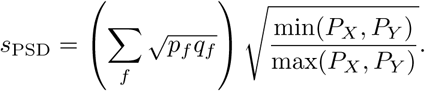

Two zero-power spectra score 1 and a single zero-power spectrum scores 0. ACF and PSD aggregate finite sequence/channel scores with equal weights on their mean and tenth percentile. The separately plotted PSD-shape and directionality diagnostics are not additional retained components. Sampling rates are 30 Hz for widefield/OmniMouse, 16 Hz for Brain Treebank and 50 Hz for IBL; synthetic BOLD controls account for their fourfold downsampling.

**Relational components.** *Correlation graph* uses pooled Pearson correlations with zero diagonal. It combines Jaccard overlap of the strongest 20% of absolute edges, agreement in absolute-weight degree, and binary clustering-coefficient agreement by a geometric mean of the finite terms, clipped to [10^−6^, 1]. Thus this score is not simply elementwise correlation-matrix error. *Cross-region mutual information* uses a five-nearest-neighbor estimator, averages the two regression directions and clips negative estimates to zero, with up to 1500 pooled samples and RNG seed 0. Its matrix comparison is 0.6*c* + 0.4*r*.

*Lagged covariance* averages matrix agreements at lags 1, 2 and 4. *VAR-response proxy* compares ridge VAR(1) coefficient matrices with ridge 0.01. The latter is the operational “impulse response” component: it summarizes a fitted one-step linear response, not a multi-horizon impulse-response identification or causal influence estimate. For these additional matrix metrics, reference and pre-diction are each median/IQR scaled separately, with SD/unit-scale fallbacks. The implementation pools finite rows and applies lagged operations to the flattened data; transitions across sequence boundaries are therefore included in these particular covariance/VAR estimates. Upper-triangular extraction also retains only one orientation of their asymmetric matrices. These components must not be interpreted as complete directed coupling recovery. ACF, PSD and discrete-state transitions use their own sequence-aware computations.

**Geometric components.** *Manifold agreement* combines stratified persistence-lifetime agreement (0.75) with a local contribution (0.25), the latter combining pooled persistence and five-nearest-neighbor distance-profile agreement. Persistence is computed for *H*_0_ and *H*_1_ with a reference PCA cap of three dimensions and deterministic point subsampling; stratified sampling takes two points from up to 48 sequences. The persistence comparison uses lifetime-distribution summaries, not a full optimal matching of diagrams. *Subspace agreement* is the mean cos^2^ *θ_i_* of principal angles between covariance eigenspaces. Each subspace aims to explain 85% of variance, capped at six dimensions, and both are truncated to their smaller rank. Covariances use shrinkage after separate robust scaling. High subspace agreement does not guarantee that all low-variance dimensions, manifold topology or task-relevant information are preserved. This limitation matters because cortical population dimensionality can continue to grow with the number of recorded neurons (Manley et al., 2024). Native-space temporal checks and the Brain Treebank lower-variance perturbation test provide complementary evidence, without establishing that the reduced representations preserve all neural information.

**State-dynamical components.** A reference median/IQR scaler and three-dimensional reference PCA define a common representation. Mini-batch *k*-means centers are fitted only to reference points (at most 10,000 fit points, ten initializations, RNG seed equal to *K*); occupancy evaluation uses at most 30,000 points per dataset. The five components comprise occupancies at *K* = 11, 12 and transition histograms at *K* = 11, lags 1, 2 and 3. Transitions are counted within sequences. Each flattened count matrix is normalized by its total count; it is a joint transition histogram, not a row-normalized conditional transition matrix. For normalized histograms *p, q*, *s*_hist_ = 0.40 min(*p_i_, q_i_*) + 0.25*c*(*p, q*) + 0.35*r*(*p, q*),

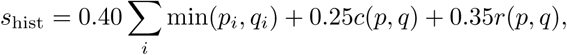

with available-term renormalization. The states are computational partitions, not annotated behavioral or latent biological states. For smooth activity, a transition may simply cross a cluster boundary; its interpretation depends on the embedding, state count and lag. The parameter controls quantify sensitivity to these choices without validating discrete biological states.

**Sidecars and numerical scope.** The displayed pointwise-fidelity sidecar combines the implementation’s error and paired mutual-information scores with weights 0.65 and 0.35. It is excluded from the structural composite and is not a proper scoring rule. The main comparisons additionally use native MAE/MSE where available. The scorer is deterministic conditional on tensors, configuration and fixed subsampling seeds; this does not remove finite-window uncertainty or data-selection effects. Different components have distinct normalization and pooling conventions, so figures illustrating intermediate moments, correlations or spectra are diagnostic explanations rather than new component definitions.

### C.2 Controlled synthetic and empirical validation

**AR-based oracle and perturbations.** The controlled generator uses a latent first-order autoregressive process and a calcium observation transform, with six latent dimensions, 16 observed regions, 100 sequences and 768 retained time bins after a 128-bin burn-in. Generator parameters are fixed by seed 11; the reference uses realization seed 101 and the 20 independent oracle realizations use seeds 501–520. Shared generator parameters do not imply identical latent paths.

The perturbation inventory changes gain/bias; time constants/frequency; phase synchronization; observation kernels; combinations of temporal parameters; coupling; channel loadings; signs; relational combinations; subspace orientation; variance allocation; rank; and geometric combinations. Each of 13 classes is evaluated at five normalized doses using realization seed 701. Dose is normalized within a manipulation and is not a common physical unit across classes. The heatmap shows the mean oracle score minus the maximum-dose perturbed score. Negative differences are retained numerically; an independent oracle is not a strict upper bound. Dose-response bands are SD across the 13 classes, not confidence intervals over seeds. Mean monotonic degradation does not imply that each class decreases monotonically or affects a unique family.

**Metric-level corruption screen.** Figure S3a retains 34 specified corruptions and 16 component readouts, with the reference fixed while transforming a fixed prediction. Rows cover mean/bias, scale/tails, additive noise, temporal rearrangement, region remapping, spectral contamination and smoothing. For component *m*, corruption *c* and severity *α*, the displayed sensitivity is

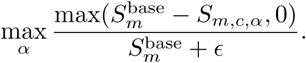

This maximum-over-dose summary describes sensitivity, not specificity, a seed-averaged effect or an inference test. Its temporal column is trajectory agreement. ACF and PSD coverage is evaluated in the separate temporal-control panel, rather than inferred from the 16-column matrix.

**Matched-error and temporal controls.** Five widefield transformations—time-order change, channel shifts, smoothing, a PCA shortcut and state cycling—are amplitude-matched to MSE 0.04956. Family-profile differences are tested with 999 blocked permutations, yielding *p* = 0.001; paired evaluation-sequence bootstraps (200 replicates per corruption) characterize profile uncertainty. The separate 20-shuffle comparison permutes time independently within each region. The contraction control defines

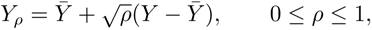

where means are taken over time within region/window. This preserves means while retaining fraction *ρ* of temporal variance. The plotted means and supplied 95% interval boundaries are preserved; they should not be interpreted as training-seed or animal-level uncertainty.

Dedicated temporal controls use 100 held-out widefield sequences of 630 bins and 16 regions, with transformations confined within sequences. Block shuffling, lag jitter, Gaussian smoothing and phase scrambling are evaluated over the full displayed dose grids, with 20 corruption realizations per setting; a fixed-dose time reversal is also retained. Larger blocks preserve more temporal order, so block length is not an increasing severity scale. A signed lagged-covariance directionality diagnostic remains outside the composite. Bands show the 10th–90th percentile over corruption realizations. Across 360 evaluations, removing PSD gives rank correlation 0.998 and mean absolute composite change 0.0046; removing ACF gives 0.983 and 0.0070. On 500 paired evaluation-sequence bootstrap rows from five fixed models, both rank correlations are 0.997, with mean absolute changes 0.0209 and 0.0206. Ablations average the two remaining temporal components equally and retain the temporal family weight of 0.18. Rank stability does not establish component redundancy or unchanged absolute scores.

**Richer generators and the spiking–calcium control.** Figure S24 evaluates nonlinear latent dynamics, switching linear dynamics, Lorenz dynamics, a low-rank recurrent network, and spiking– calcium observations. Five generator seeds (7, 11, 19, 23, 31) each yield eight sequences of 160 bins and 12 channels. The independent oracle is a new realization from the same generator. A noisy-reference control adds Gaussian noise with SD 0.05; it is not an empirical split-half comparison. All 18 retained components are finite, including trajectory, ACF and power-sensitive PSD.

The spiking–calcium comparison distinguishes an independent oracle, a pure shuffle, and an MSE-matched reference–shuffle mixture. For a channel-wise shuffle *π*(*X*) across pooled sequence/time samples,

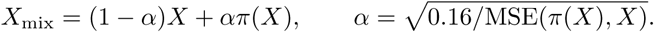

The implementation’s clipping limits are inactive here. Across seeds, *α* = 0.2821–0.2846, retaining 71.54–71.79% reference coefficient. Mean composites are 0.73005 for the mixture, 0.66640 for the oracle and 0.50360 for the pure shuffle, with MSE 0.160, 1.770 and 1.993. The mixture exceeds the oracle in distributional, temporal, relational and geometric agreement; the oracle exceeds it in state dynamics. Relational and geometric gains contribute 0.04900 to the total mixture-minus-oracle difference of 0.06365. The oracle exceeds the pure shuffle in all five seeds. This comparison explains the reference-retaining control’s advantage; it does not identify a finite-sample mechanism or establish that a pointwise-close mixture is a viable forecast.

**Synthetic BOLD observations.** Nonlinear, switching and low-rank recurrent systems undergo hemodynamic-response convolution and fourfold downsampling. Five seeds, three systems and five controls yield 75 scores. Controls comprise repeated observations of the same neural realization with independent observation noise/drift, a noisy neural reference, an independent oracle, an MSE-matched transition mixture before the observation transform, and a pure BOLD time shuffle. Each seed is averaged equally over the three systems, and figures show the five seed means. The 40-bin sequences fail the quantile component’s 80-sample threshold, so these scores use 17/18 components, redistributing only the within-distribution weight. All three temporal components remain included and the sampling-frequency parameter changes from 30 to 7.5. Mean composites are respectively 0.848, 0.848, 0.745, 0.778 and 0.500. Neither ordering nor its differences from the neural-domain panel isolate a causal effect of hemodynamic smoothing. These controls are synthetic observation-model tests, not validation on real fMRI.

#### C.2.1 Comparison with angular DSA

We compared angular Dynamical Similarity Analysis (DSA) (Ostrow et al., 2023) with Nethobench’s temporal–spectral (T) and state-dynamical (S) families on synthetic oscillations and widefield calcium forecasts. Using the 100-sequence synthetic realization underlying Figure 2, we separately halved or doubled oscillation frequency and amplitude. Both frequency changes increased angular DSA distance and reduced T and S agreement. Amplitude scaling distinguished the measures. Angular DSA distance remained at 0.00352 for the unchanged, half-amplitude, and double-amplitude conditions, whereas T fell from 0.840 to 0.583 and 0.557, respectively, and S fell from 0.965 to 0.390 and 0.424. For each of the eight widefield forecasters, we then averaged scores over 200 sequences and computed Spearman’s *ρ* across the eight model means, using negative angular DSA distance so that positive correlations indicate agreement in model rankings. The correlations were 0.810 for T, 0.714 for S, and 0.833 for their mean. The corresponding correlation was −0.048 for the mean of the distributional, relational, and geometric families and 0.429 for the full composite. Thus, angular DSA ranked these forecasters similarly to the two dynamics families, while its ranking gave little indication of which forecasters best preserved the other three properties on average.

### C.3 Metric-selection and robustness analyses

**Candidate inventory.** The candidate audit contains 45 metric candidates and six criteria: oracle agreement, targeted-corruption selectivity, model utility, nonredundancy, coverage and interpretability. Eighteen candidates belong to the retained structural specification; complete criterion records are available for 16, while ACF and PSD have dedicated controls rather than complete screening records. Twenty-five candidates have an unmeasured nonredundancy entry that used a neutral 0.5 fallback in the rubric. These entries are masked, not treated as measured evidence, in Figure S5. Group summaries use available entries with criterion-specific denominators. Interpretability is a design judgment, and utility on a selection panel is not independent validation on that panel. Predictive MAE/correlation proxies address different targets and are not declared inferior because of a structural-screening rubric.

**Parameter and family-weight sensitivity.** Twelve alternative PCA, state-count and lag settings are compared on exactly 45 matched seed/checkpoint observations (15 training seeds at epochs 10, 30 and 100). Rank correlations always compare the same condition set. Separately, 5000 shared family-weight draws use RNG seed 20260918. Their mixture is 70% Dirichlet(80*w*), 10% equal family weights, 10% uniformly selected single-family omission, and 10% uniformly selected single-family weight doubling, with renormalization; *w* = (0.22, 0.18, 0.24, 0.18, 0.18). Each draw applies the same weights to every condition. Components, within-family averaging and score transforms are fixed.

The violin distributions in Figure S1 show, for each draw, the mean of 15 paired seed contrasts. Their central 95% ranges describe specification sensitivity, not sampling confidence intervals. Mean contrasts 100–10, 30–10 and 30–100 are positive for all draws. The first two are positive in every seed for all draws, but 30–100 is positive in every seed for only 0.0088 of draws. Thus stable mean ordering coexists with individual reversals and nonmonotonic training trajectories.

**Dependence and omission.** Family correlations are computed separately on the 45 training observations and 360 temporal-control observations. Both contain dependent repeated evaluations and support descriptive correlations rather than independent-replicate inference. Average-linkage clustering uses distance 1 *ρ*_Spearman_ among the 18 components. For a full composite *C* and a composite *C*_−_*_f_* omitting one family with renormalized weights, the loss of corruption sensitivity is

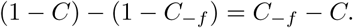

A positive value means omission makes the corruption appear less severe. This measures an aggregation change, not causal unique information, semantic decoding or incremental downstream utility.

**Five-checkpoint panel and PCA choices.** The fixed-checkpoint analysis aligns five Transformer forecasts to a common 100-sequence, 630-bin, 16-region tensor. This panel is distinct from the principal eight-model, forecast-only experiment. At state-representation PCA dimensions 2, 3, 5 and 8, rank correlations with the default are 0.90, 1.00, 0.90 and 1.00. The TF distributional-loss variant remains first, while 1-step and Sequifier exchange order at dimensions 2 and 5. Retained reference variance after median/IQR scaling is 81.0%, 88.4%, 94.9% and 98.4%. These values do not quantify retained task information. Replacing the temporal family by native-space (ACF + PSD)*/*2 yields pooled *ρ* = 0.9761 over 500 rows from 100 paired sequence bootstraps and five models. These are evaluation resamples, not independent fits.

**Empirical references and subsets.** Twenty random 50/50 partitions of the same 100-sequence pool provide real–real references. Twenty transformations each provide independent within-region time permutation, joint time–region permutation, and regional marginal resampling with replacement. Reference scores compare 50 versus 50 sequences, whereas the five forecast markers score all 100; these are descriptive reference points rather than sample-size-matched normalized performances. Repeated partitions are not independent subjects.

All 31 nonempty family subsets retain relative default weights and renormalize. Across 100 paired sequence bootstraps, subset and full scores are compared on the same five models, producing 3100 subset/bootstrap rows. Pairwise agreement averages over all ten model pairs; a tie in only one ordering counts as disagreement. The temporal family alone has median rank correlation 1.00, central bootstrap range [0.90, 1.00], and mean pairwise agreement 0.971. One family can therefore recover coarse ordering on this panel. Multiple family profiles still distinguish structural failure types; the subset experiment establishes neither a universal minimum number of families nor individual indispensability. Best subsets are selected descriptively within this panel, without an independent selection/test split.

### C.4 Replicated endpoints and training trajectories

The source-domain generator combines latent network dynamics and state switching, regional spiking with refractory effects and inter-region coupling, AR(2) calcium rise/decay filtering, and neuropil, observation-noise and baseline-drift terms. “Biophysical” names this artificial data-generating process, not the forecasting architecture. The forecasters use the Sequifier Transformer architecture. CalciumGAN is a WaveGAN-based model with a Wasserstein training objective for synthesizing population calcium traces (Li et al., 2020); here its generated traces supply the transfer domain. The endpoint comparison uses five paired runs (seeds 2001–2005) of a low-capacity model (approximately 11,000 parameters, 16-bin context, one layer, one epoch) and a higher-capacity model (approximately 1.58 million parameters, 128-bin context, six layers, validation-selected weights and last logged epoch labels 65–69). Capacity, context, depth and training co-vary. Both forecasts are rescored on exactly matching targets within domain: 96 source sequences at bins 128–1039 (912 generated bins) and 128 CalciumGAN-transfer sequences at bins 128–1023 (896 generated bins), retaining 16 regions. Context is excluded, although forecast origins differ with context length. CalciumGAN is a generated target domain, not another animal cohort.

Mean higher-minus-lower composite differences are 0.1317 in the source domain and 0.1206 in transfer. Unadjusted paired Student-*t* 95% intervals with four degrees of freedom are [0.0074, 0.2707] and [0.0212, 0.2199]; both exact two-sided sign-flip tests give *p* = 0.125. We do not infer uniform family improvement or isolate the effect of an individual design variable.

A separate 15-seed training experiment evaluates epochs 10, 30 and 100 on 45 validation sequences of 810 generated bins (Figure S25e). Mean composites are 0.4742, 0.6163 and 0.5680, respectively. Epoch 100 exceeds epoch 10 in all seeds, while epoch 30 exceeds epoch 100 in 14/15. The latter exact paired sign-flip *p* = 0.000122 becomes *p* = 0.000183 after Holm adjustment over the three checkpoint contrasts. Thus, structural agreement changes nonmonotonically during training; “peak” refers only to the sampled checkpoints.

A separate fixed-epoch comparison evaluates six forecasting configurations - TF, TF QL KL, AR, 1-step, RNN and SSM - with 3 training seeds per configuration (Figure S25f-g). All configurations are evaluated at epoch 150 on the same validation sequences. Panel f reports the Nethobench composite and panel g the normalized forecast MSE. Points represent individual training seeds, lines connect matching seed identifiers across configurations, and bars show the seed mean SEM. These fixed-epoch comparisons are distinct from the validation-selected checkpoints used in the principal widefield comparison. Model capacity and compute are not matched, and seed SEM describes training variability on the fixed evaluation set rather than animal or session uncertainty.

### C.5 Proper scores and statistical conventions

Before probabilistic scoring, targets and draws are transformed by the same per-region target median and 1.4826 MAD (SD/unit-scale fallbacks). This evaluation normaliza_√_ti<u>on i</u>s shared across methods and does not fit a forecasting model. Energy distances are divided by *TR*, with the pairwise draw term including the zero diagonal. The lag-restricted variogram uses *p* = 0.5, lags *L* = {1, 2, 4, 8, 16} and normalized weights *w_ℓ_* ∝ 1*/ℓ*:

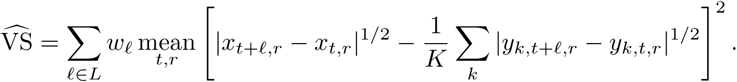

Scores are averaged over contexts. We report the empirical finite-ensemble implementation rather than claiming an unbiased estimator of an infinite-ensemble score. Neither energy nor a lag-restricted variogram alone localizes every structural failure.

All uncertainty units are stated locally: training-seed SD after averaging evaluation groups for the principal widefield comparison; split SEM for OmniMouse; seed variation for replicated fits; corruption-realization ranges for perturbations; specification ranges for weight draws; and participant-level inference for Brain Treebank. Independent synthetic realizations are not biological replicates. Exact paired sign-flip tests enumerate the available seed/participant signs when feasible, and adjusted results specify the contrast family. We avoid interpreting raw differences between different metric scales as effect sizes or pooling the distinct experiments into a global significance claim.

### C.6 Within-sequence relational and conditional-transition sensitivity

The primary figures and Tables 9-10 use the implementation described in Appendix C.1. We separately evaluate within-sequence, directed relational comparisons on fixed synthetic controls (Figure S26). These variants were evaluated in supplementary sensitivity analyses; the principal empirical figures and tables retain the original implementation.

**Sequence-safe directed relational comparisons.** Reference and forecast are median/IQR scaled separately as in the pinned implementation. For lag *ℓ*, define valid within-sequence pairs

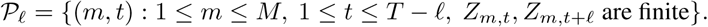

Let *A_ℓ_* and *B_ℓ_* contain the corresponding earlier and later row vectors. The corrected lagged covariance is

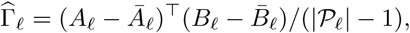

using lags 1, 2, 4. The one-step VAR-response proxy is

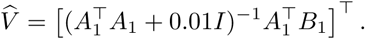

Both comparisons use every directed off-diagonal entry, vec*_i_*_̸=*j*_(), with the same 0.6*c* + 0.4*r* similarity as the pinned component. No last-to-first transition connects different sequences. Tests verify pair counts, sequence-permutation invariance, channel-permutation invariance, and sensitivity to a change confined to the previously omitted triangle. The two old components were first reproduced from every saved tensor before their differences were computed.

The corrected relational variant changes the composite by at most 0.00485 in the neural controls and 0.01059 in simulated BOLD. It preserves candidate ordering in all 40 system-by-seed groups (Figure S26). These are finite-panel sensitivity results; they do not establish unchanged rankings for the principal widefield models or other datasets.

Rescoring frozen forecasts further preserves the mean composite ordering of the three tested widefield models and all Brain Treebank models, including the latter’s structural-accuracy reversal (Figure S19).

**Conditional state transitions.** For each of the same reference-fitted *K* = 11 partitions and lags 1, 2, 3, let *N_ij_* count transitions within sequences. The pinned score normalizes all entries by *_ij_ N_ij_* and therefore combines occupancy and transition information. The alternative compares conditional rows, *P_ij_* = *N_ij_/ _j_ N_ij_*, on origin states with positive reference counts. It averages the existing histogram similarity uniformly over these states, retaining its overlap/correlation/RMSE terms. An origin state visited in the reference but never left in the prediction receives zero agreement; absent reference origin states are excluded. The two occupancy components are unchanged. Thus, the alternative reduces weighting by how often an origin state occurs, while still penalizing unsupported predicted states.

Conditional-row scoring changes neural composites by at most 0.01327 and BOLD composites by at most 0.01134, preserving ordering in 38 of 40 system-by-seed groups. Applying both variants yields maximum absolute changes 0.01375 and 0.00972, respectively. These checks expose a modeling choice rather than establish uniquely correct state partitions or independent families. All variant component scores, composites, support masks and ordering comparisons are retained with the analysis.

## D MODELS

We compare linear, recurrent, Transformer, and stochastic latent state-space forecasters. We first define the shared widefield protocol, then specify the models and their training objectives. The two-photon extension uses a separate model and observation protocol.

### D.1 Shared widefield protocol

Let *x_t_* ∈ R*^N^* denote neural activity, *X* = *x_t_*_−_*_T_* _+1:*t*_ the context, and *Y* = *x_t_*_+1:*t*+*H*_ the training target. The principal comparison uses *N* = 16 regions sampled at 30 Hz and *T*_in_ = 90 (3 s). The training horizon is *H* = 90, except for the next-step Transformers, which use *H* = 1 (Table 7). Data preparation and partitions are specified in the experimental-setup appendix.

**Autonomous evaluation.** Only the initial 90-step context is observed. Each subsequent prediction is conditioned on generated activity. Multi-step Transformers, RNN, and SSM generate 90-step blocks and re-encode the latest 90 generated observations at each boundary; Transformer positions and attention caches, recurrent states, and SSM initial-state distributions are reset there. Within a block, states are retained. VAR and next-step Transformers update a sliding 90-step context after every prediction. The observed context is excluded from scoring. The 3-, 12-, and 24-s analyses score the first 90, 360, and 720 forecast steps: these are cumulative windows, so changes in scores also reflect changes in the amount of data available for metric estimation.

### D.2 Deterministic forecasters

**Linear autoregression (VAR).** The VAR baseline predicts *x*^*_t_*_+1_ = *W* vec(*X*) + *b*, allowing every output region to depend linearly on every region and lag in the context (Lütkepohl, 2005). Training applies this map to ground-truth sliding windows across the target block; inference recursively substitutes predictions for unavailable observations.

**Recurrent model (RNN).** A unidirectional GRU encodes *X* from a zero hidden state. The first prediction is an affine readout of the layer-normalized top-layer state. The GRU then consumes the preceding target observation during training, or its own preceding prediction during evaluation, before applying the same readout. Dropout acts between recurrent layers. Training minimizes MAE over all *H* future steps.

**Shared Transformer backbone.** The 1 step, TF, AR, and TF_QL KL_ variants use an affine observation embedding *ϕ*, learned positional embeddings, and a learned future-start token *s*. Each causal decoder block contains pre-layer-normalized self-attention and a GELU feed-forward network with residual connections. A final layer normalization and affine readout produce the observations. Positions are relative to the current context–forecast block, with *T*_in_ + *H* learned positions.

**Training regimes.** For teacher forcing (TF), the input tokens are [*ϕ*(*X*)*, s, ϕ*(*Y*_1_)*, . . ., ϕ*(*Y_H_*_−1_)]; a causal mask prevents each prediction from accessing its own or later targets. All future outputs are computed in one pass. AR instead recursively feeds back its predictions during training. Its attention history grows throughout the block, and gradients propagate through predictions and cached states without detachment. The 1 step baseline uses the same backbone with *H* = 1, so only the context and start token are supplied. TF and AR minimize MAE; TF_QL KL_ uses TF with the augmented objective below. Schematics are provided on Figure S9.

**Sequifier.** Sequifier (Luithlen, 2025) is an additional causal Transformer implementation trained on the next-step *ℓ*_1_ objective (*H* = 1) and applied recursively at evaluation. Its dimensions and optimization settings are given in Tables 7 and 8. Operationally, Sequifier encodes input regions separately and predicts the next activity vector from the final observed position; our 1 step model jointly embeds the activity vector and predicts from an appended learned future-start token. Both feed predictions back one step at a time during evaluation.

### D.3 Distribution-augmented Transformer objective

For a batch of *B* examples, let 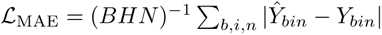. The augmented objective is

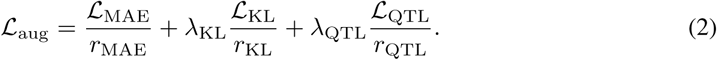

Each *r_j_* is the detached component value on the first training batch plus 10^−8^, held fixed thereafter. The MAE-only Transformer objective uses zero distributional weights. For TF_QL KL_, the weights are *λ*_KL_ = 0.02 and *λ*_QTL_ = 0.08. Both added losses compare within-window marginals; neither directly constrains temporal order or cross-region dependence.

**Soft-histogram loss.** For each region, 33 Gaussian bins span the 0.001–0.999 quantiles of pooled target and detached prediction values across batch and time. The Gaussian width is half the bin width.

Soft counts are summed over time, stabilized by 10^−8^, and normalized separately for each sequence and region. If *p_bn_* and *q_bn_* are the resulting target and prediction histograms, respectively, define

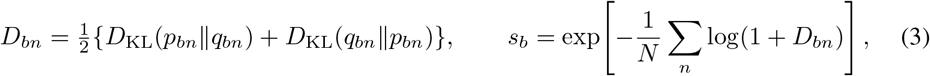

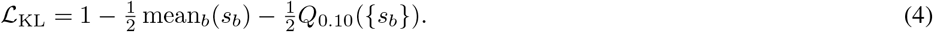

Here *Q_q_* is the empirical quantile. Histogram support is detached, but soft assignments remain differentiable. The divergence is symmetric KL.

**Tail-quantile loss.** Temporal quantiles are computed at 99 levels from 0.01 to 0.99, retaining *q* 0.10 or *q* 0.90. For each sequence and region, the mean absolute prediction–target difference over these tail quantiles is divided by the target region’s interquartile range pooled across batch and time (minimum 10^−8^). Average the largest ⌈*N/*4⌉ regional discrepancies per sequence and then across sequences to obtain *D*_QTL_. The loss is L_QTL_ = 1 − (1 + *D*_QTL_)^−1^.

### D.4 Conditional latent state-space model (SSM)

**Generative model.** A context GRU parameterizes the diagonal Gaussian initial state *p*(*z*_0_ *X*). The conditional model is

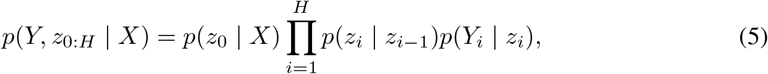

with *p*(*z_i_* | *z_i_*_−1_) = N(*m*(*z_i_*_−1_), diag(*σ_z_*^2^)) and *p*(*Y_i_* | *z_i_*) = N(*d*(*z_i_*), diag(*σ_x_*^2^)). The transition combines linear and nonlinear dynamics,

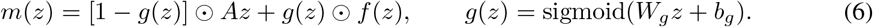

The linear matrix is normalized to spectral norm at most 0.99; this bound does not constrain the full nonlinear transition. The transition MLP *f* has tanh activations after both affine layers, while decoder *d* has tanh hidden layers and a linear output. Transition and observation variances are learned, diagonal, and state-independent. Log variances are clipped to [12, 8]; transition scales use softplus plus 10^−4^. Table 7 gives the layer dimensions. Initial-state and transition standard deviations start near 0.1; the posterior mean correction starts at zero and the gate bias at 4, initially favoring the linear transition.

**Variational training.** A second GRU, initialized from the context encoder, processes the target prefix *Y*_1:*i*_. Its state and *z_i_*_−1_ enter a 108 → 96 → 24 MLP with a tanh hidden layer to parameterize

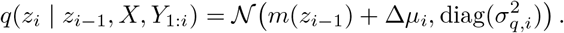

The objective uses reparameterized latent samples:

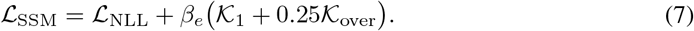

The Gaussian negative log-likelihood sums over time and channels and averages over examples. _1_ likewise sums latent KL terms over time and dimensions, applying a free-bits floor of 0.05 nats to each example–time–dimension term before averaging across examples. *β_e_* increases linearly from zero at epoch 1 to 0.2 at epoch 45 and then remains fixed. The initial state is sampled from *p*(*z*_0_ *X*); there is no separate initial-state KL.

Overshooting compares target posterior distributions with endpoint transition Gaussians after 5, 15, and 30 autonomous steps from four evenly spaced posterior anchors per horizon. Intermediate latents are sampled; anchors and target posterior parameters are detached. For each horizon, *D*_KL_(*q*_target_ *p*_endpoint_) uses the same per-dimension floor, sums over latent dimensions, averages over anchors and examples, and is multiplied by *H*; K_over_ averages these values over horizons.

**Forecasts used for evaluation.** Evaluation uses only the context-conditioned generative model, never the target-informed posterior. Initial states and transitions are sampled, but the output is the decoder mean *d*(*z_i_*) without observation noise. The configured evaluation draws one stochastic latent trajectory per context; this is the primary prediction. Thus, structural scores evaluate sampled SSM forecasts, whereas the other forecasters produce deterministic outputs. Each SSM forecast represents latent-process variability, not a full observation sample, and supplies its own context at subsequent block boundaries. Averaging multiple forecasts from the same context instead approximates the model’s conditional mean, attenuating fluctuations with variable timing while retaining predictable temporal structure (Figure S10b). The separately exported deterministic trajectory recursively propagates latent means; because the dynamics are nonlinear, it generally differs from the Monte Carlo mean used for checkpoint selection.

### D.5 Two-photon extension: population-conditioned MLP

The separate 88-neuron calcium example uses a shared temporal MLP to encode each neuron’s 90-step history, adds learned neuron embeddings, and conditions these features on a population summary. The summary concatenates each neuron’s last value, temporal mean, maximum, and change over the last five steps. A population encoder produces feature-wise scale and shift terms. The prediction head emits *H*_MLP_ residuals *r_in_* per neuron and adds a learned persistence skip:

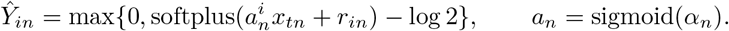

The local and population widths are 64 and 128, with GELU activations and dropout 0.05. The training configuration uses *H*_MLP_ = 16, *a_n_* = 0.965 at initialization, AdamW with learning rate 10^−3^ and weight decay 10^−4^, batch size 1024, 30 epochs, and gradient clipping at norm 1. Training and checkpoint selection use weighted smooth-*ℓ*_1_ loss, with unit smooth-*ℓ*_1_ transition threshold and weights 1+2 min 10, max(0*, Y_in_/s*) . Here *s* is the 95th percentile of strictly positive training-target values, bounded below by 10^−6^ (or 1 if none are positive).

The inference implementation supports direct block prediction followed by resets to recorded context. These forecasts receive intermittent observations and are not equivalent to the autonomous widefield rollouts above.

## E Code, Data and Reproducibility

**Overview.** We release the processed widefield-calcium forecasting dataset (Appendix B.1), the anonymized code used to train models, generate rollouts, compute Nethobench scores, and reproduce the main figures, and the widefield model suite checkpoints. The two-photon dataset (Appendix B.5) will be released on Zenodo upon publication. Dataset construction, splitting, normalization, and long-rollout generation are described in Appendix B. Model architectures and training details are described in Appendix D, and metric definitions are provided in Appendix C.

**Dataset.** The processed benchmark dataset is available at:

https://huggingface.co/datasets/Nethobench/nethobench-widefield-v1

**Code.** The code is provided in two anonymized repositories for review, released under MIT License. The Nethobench repository contains the final metric implementations, family-level and composite scoring code, documentation, and examples intended for public reuse.

https://github.com/andrestrocyte/nethobench

The forecasting repository contains the model training and evaluation pipeline for widefield and two-photon calcium imaging, including rollout-generation scripts and Nethobench-evaluation workflows:

https://github.com/blanchemarion/sequential_neural_pred

**Checkpoints.** The widefield models suite checkpoints are available at:

https://github.com/blanchemarion/submission_checkpoints

**Reproducibility workflow.** To reproduce the main benchmark analyses, users should download the processed dataset, train or load a forecasting model, generate closed-loop rollouts from the provided context windows, and evaluate the generated trajectories against held-out target windows using the Nethobench scoring code. The reported outputs are the five family-level structural-realism scores, the final composite score, and pointwise-fidelity sidecar metrics.

### E.1 Evidence roadmap

The roadmap separates support, availability and units of variation. Sequences, model seeds, corruption draws and participants are not interchangeable independent units. Absolute scores should only be compared on matched supports and definitions.

Evaluation questions and evidence.

| Question | Evidence and interpretation boundary |
|---|---|
| Temporal order | Figures S3, S4, S21: shuffles, blocks, jitter, ACF/PSD ablations. Sensitivity is not complete dynamical identification. |
| Under-dispersion | Figure S2: mean-preserving variance contraction; Figure S22: proper scores and coverage. Structural sensitivity and conditional calibration are separate. |
| Empirical anchors | Figure S2: real–real references, structured shuffle or resampling controls, and variance contraction; no universal ceiling or floor. |
| Parameter sensitivity | Figures S1, S7, S26: weights, PCA, states, lags and explicit implementation variants. Fixed-panel stability does not certify new supports. |
| Family dependence | Figures S6 and S8: correlated scores, omissions and all subsets. Diagnostic complementarity does not require all families for ranking. |
| Stochastic validity | Figures S24 and S22: correct-process samples versus predictive-distribution scoring. Structural agreement does not imply conditional calibration. |
| Forecast representation | Figure S10: representative model rollouts and sampled SSM forecasts versus their pointwise mean. Averaging attenuates variability; architectural and sampling effects are not isolated. |
| Cross-modality scope | Figures S15, S24, S23, S21: distinct recordings and observation transforms; scores are not directly comparable across them. |
| Human protocol | Appendix B.6: training-only contact selection and exact analog-library exclusions; offline transforms and the changed contact fold remain limitations. |

## F SUPPLEMENTARY FIGURES

**Figure S1:**
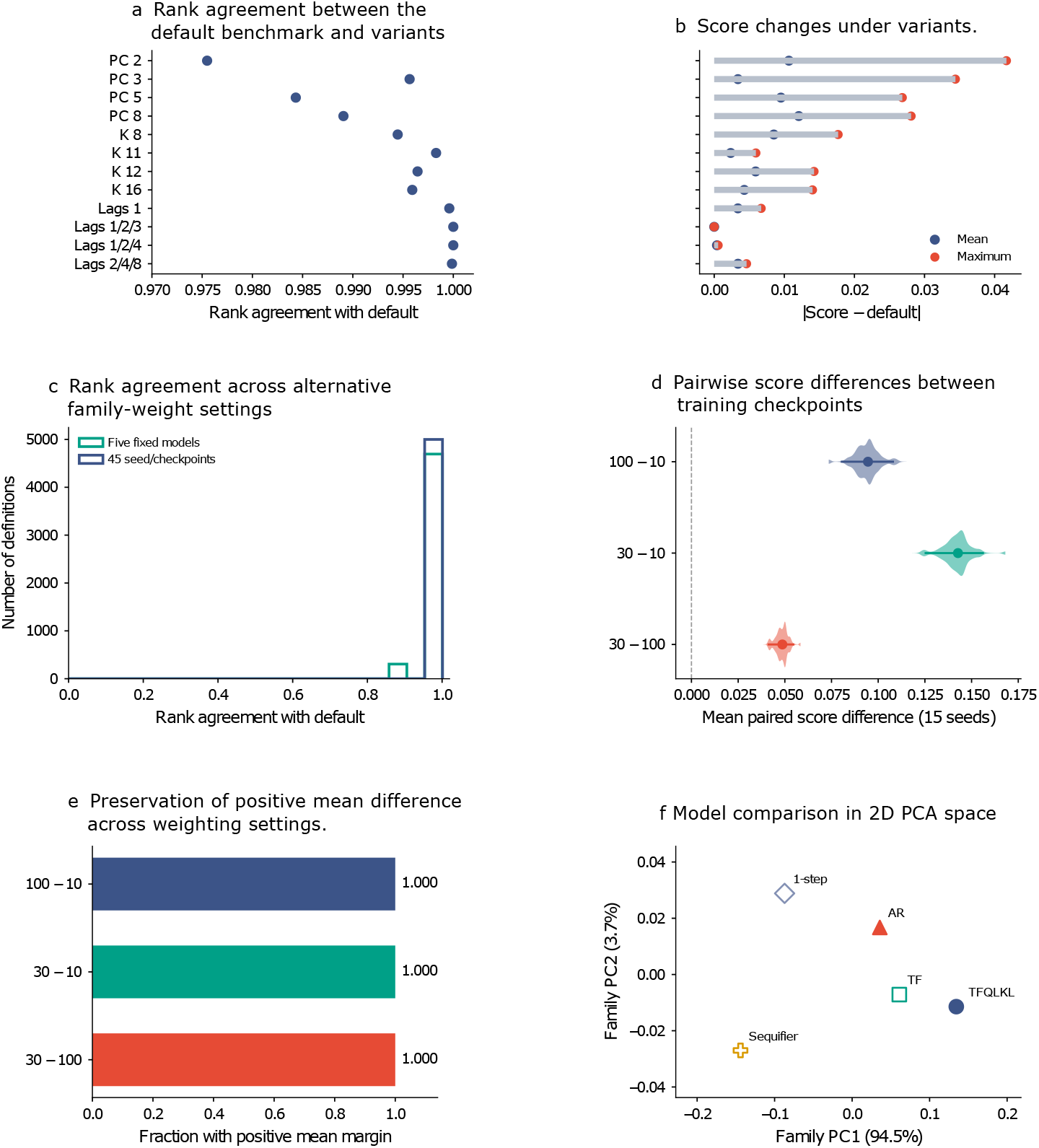
Robustness to benchmark definition and family weighting. (a) Rank agreement between the default benchmark and variants using different PCA dimensions, latent-state counts (*K*), or transition lags across 45 matched seed-checkpoint conditions. (b) Mean and maximum absolute score changes under the same variants. Rankings remain stable although absolute scores shift slightly. (a-b) 15 seeds, 3 checkpoints. (c) Rank agreement across 5,000 alternative family-weight settings for five models and 45 seed-checkpoint conditions. The ensemble mixes Dirichlet(80*w*) draws (70%), equal weights (10%), single-family omission (10%) and single-weight doubling (10%), with renormalization. (d) Mean paired score differences between training checkpoints across 15 seeds under each weighting. Intervals describe specification sensitivity. (e) Fraction of weighting settings preserving a positive mean difference. All three contrasts remain positive across all settings. (f) PCA visualization of the five-dimensional family-score profiles. Filled markers indicate Pareto-nondominated models. Further parameter- and weighting-sensitivity details are specified in Appendix C.3.

**Figure S2:**
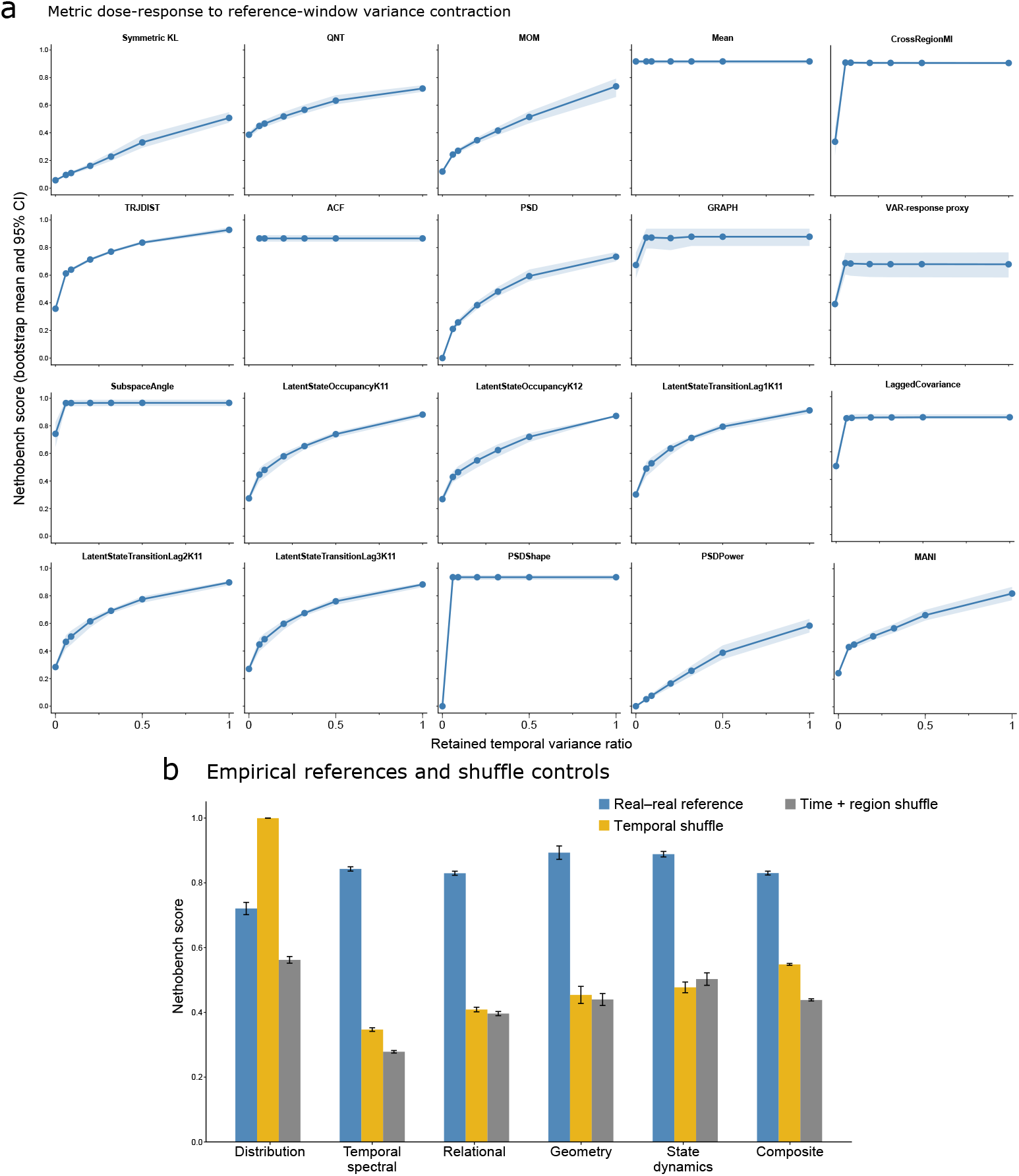
Empirical references and sensitivity to temporal-variance contraction in widefield neural activity. **(a)** Metric-level responses to progressively contracting the temporal variance of the second window in each split-half pair. For each region, the contracted activity was defined as 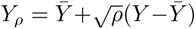, thereby retaining a fraction *ρ* of the original temporal variance while preserving the temporal mean. Thus, *ρ* = 1 corresponds to the unmodified real window, whereas *ρ* = 0 produces a temporally constant signal at each region’s mean. Lines show bootstrap-mean Nethobench scores, and shaded bands indicate 95% confidence intervals obtained by paired resampling of joint batch scores. **(b)** Family-level and composite scores for matched empirical references, using the same 222 evaluation targets and 720-timestep forecast-only windows as the principal model comparison. For each target, the real-real reference samples a non-overlapping 720-timestep companion window from the same recording. Temporal shuffling independently permutes time within each region, preserving regional marginals while disrupting temporal structure and inter-region coordination. Time-and- region shuffling permutes all entries within each target sequence, additionally disrupting regional identity and regional marginals. Each of 20 random draws is scored within four fixed evaluation groups of 56/56/55/55 sequences. Bars show means across draws after equally averaging the four group scores within each draw; error bars show the Monte Carlo sample SD of these draw means. Further details on the widefield evaluation and temporal controls are specified in Appendices B.1 and C.2.

**Figure S3:**
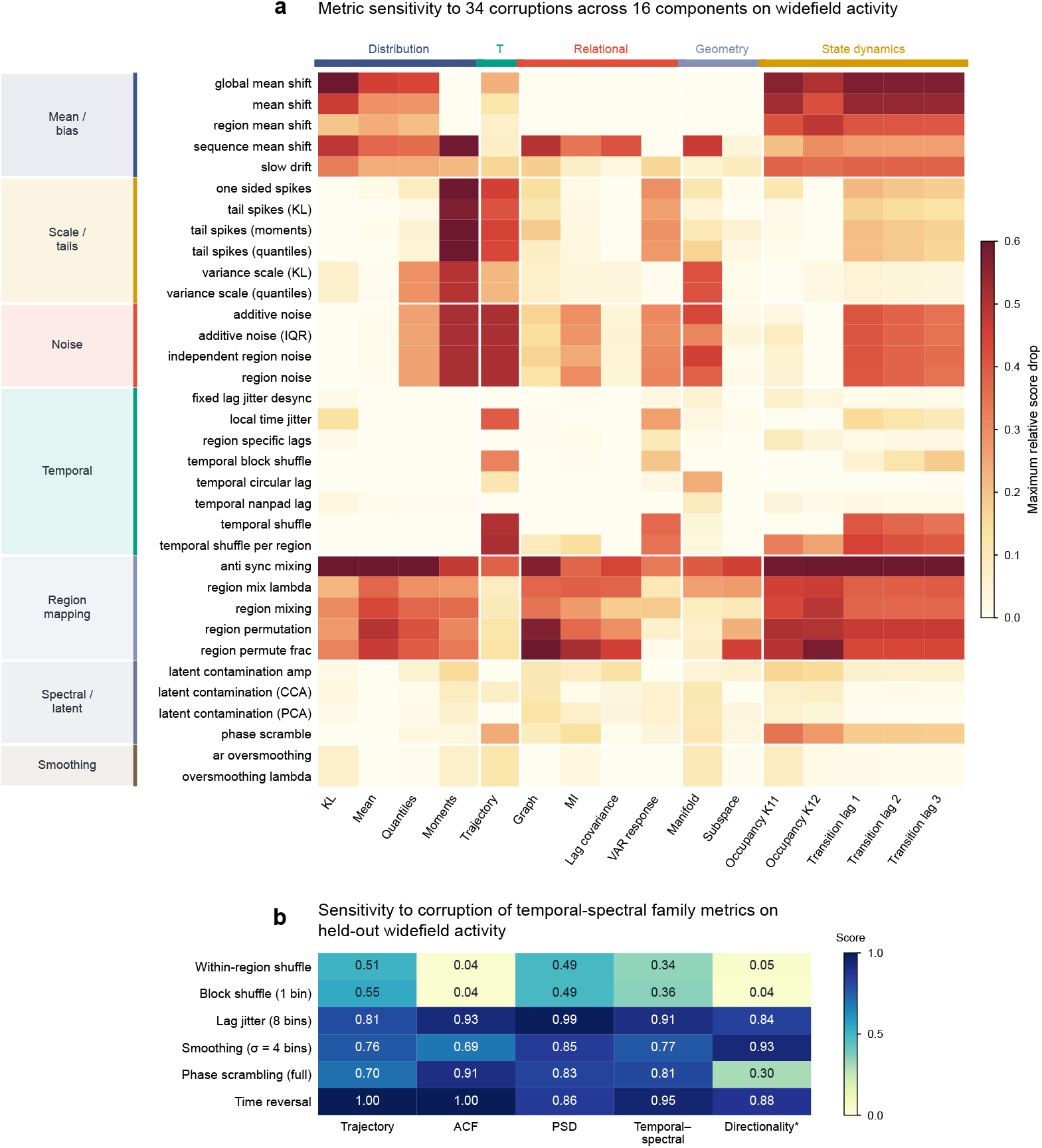
Metric-level responses to specified corruptions. (a) Maximum relative score drops over tested severities for 34 corruptions and 16 components; the reference is fixed. Labeled row blocks distinguish corruption classes. (b) Dedicated trajectory, ACF, PSD and directionality controls on held-out widefield activity; cells average 20 corruption realizations at the displayed dose. Directionality is a signed lagged-covariance diagnostic excluded from the composite. Replicates are transformations of a fixed 100 630 16 tensor. Overlapping responses do not imply exclusive family selectivity. Further details on the controlled corruptions are specified in Appendix C.2.

**Figure S4:**
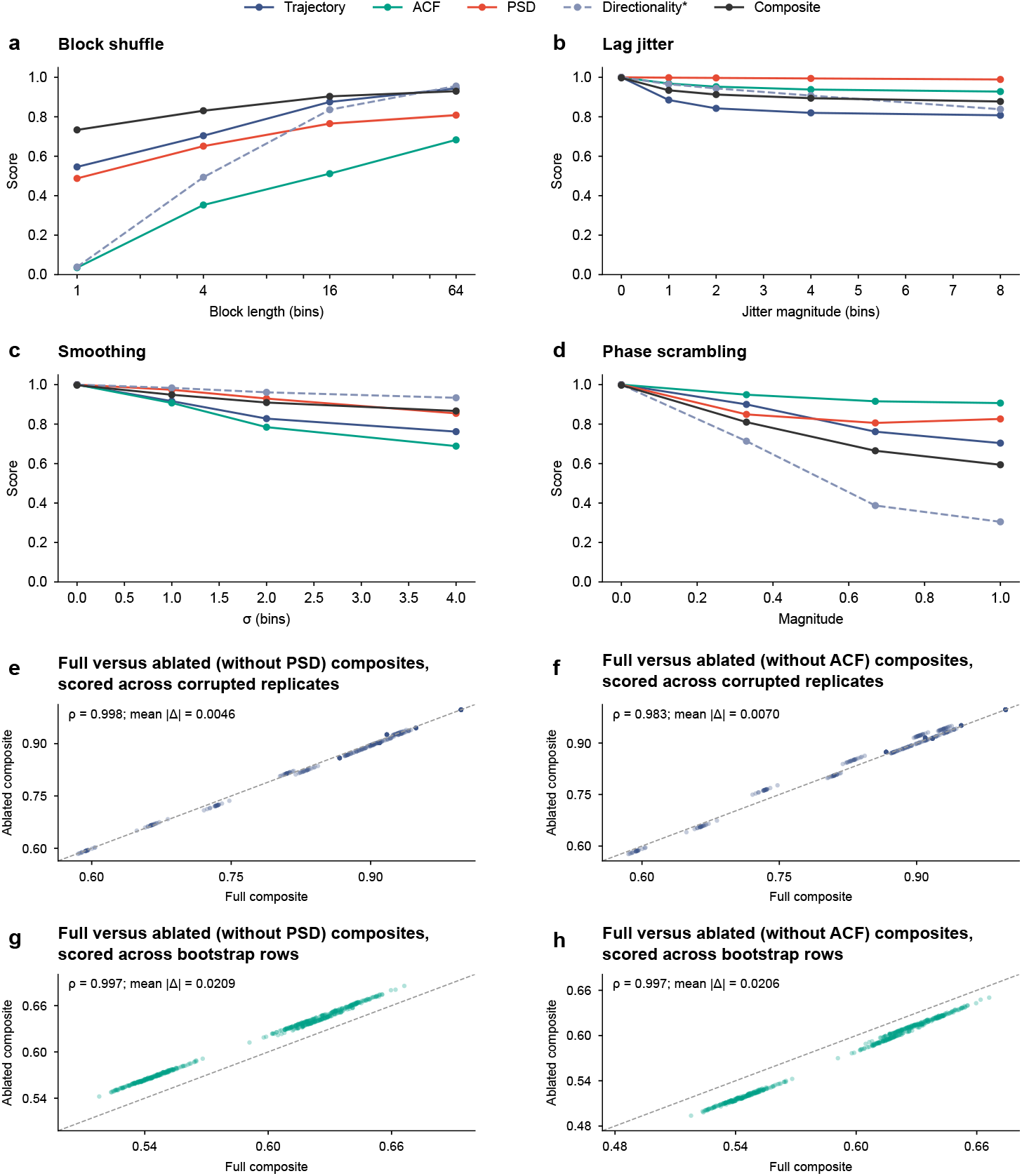
Temporal sensitivity and component ablations. (a–d) All doses of block shuffling, jitter, smoothing and phase scrambling. Lines are means and bands span the 10th-90th percentile over 20 corruption realizations. Longer blocks preserve more temporal order. (e–f) Full versus ablated composites, scored across 360 temporal controls, including time reversal. (g–h) Analogous comparisons, scored across 500 evaluation-bootstrap rows (100 paired sequence resamples, five fixed models). Removing ACF or PSD averages the remaining two temporal components equally while retaining family weight 0.18. Directionality remains outside the composite. Further temporal-control and ablation details are specified in Appendix C.2.

**Figure S5:**
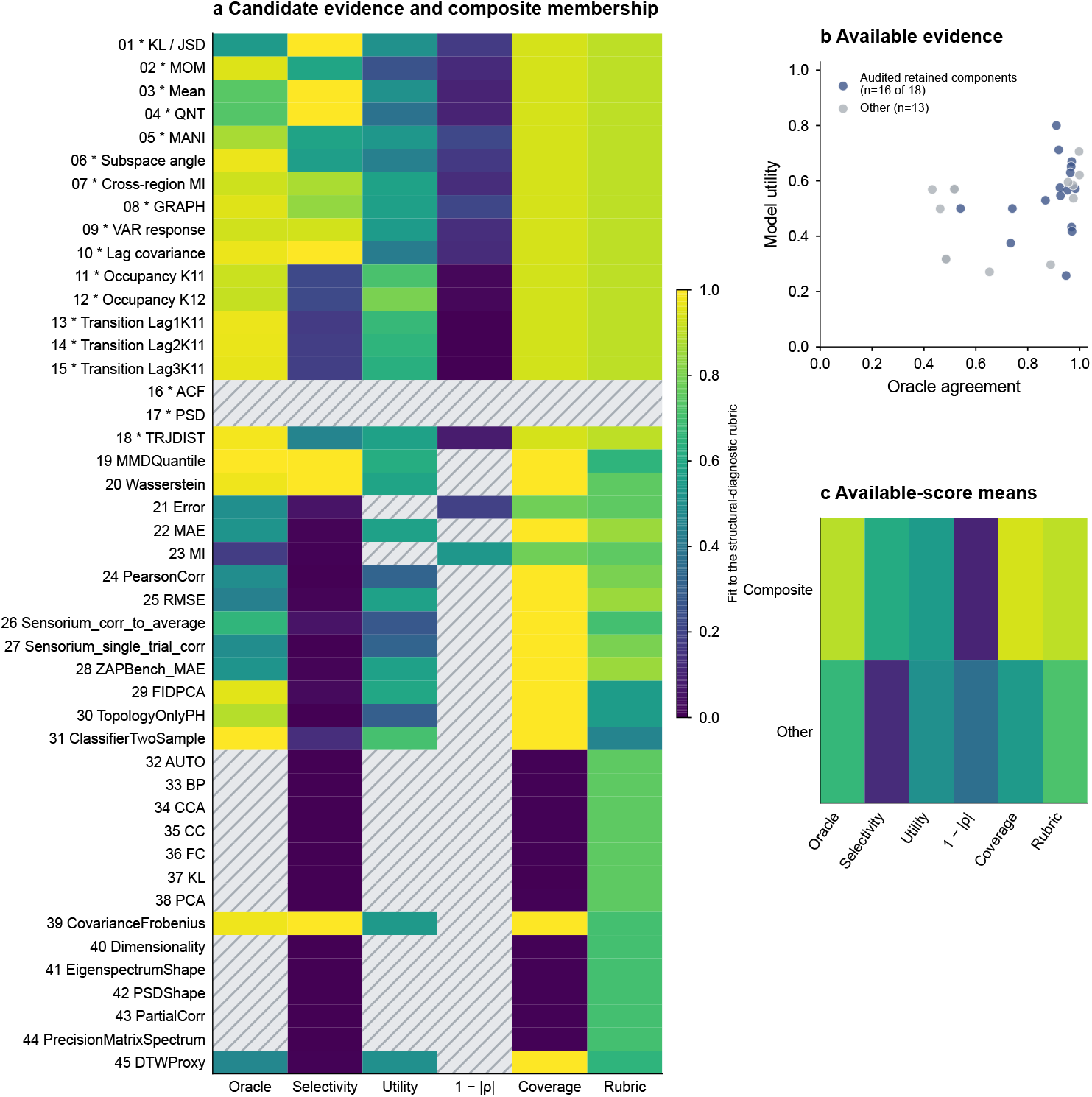
Metric construction and retrospective design evidence. (a) Six screening criteria for 45 candidates; asterisks mark 18 retained components. Hatched cells indicate unavailable measurements. (b) Oracle agreement versus utility for components with both available (16 of 18 retained). (c) Available-entry criterion means. Utility on a selection panel is not independent validation. Predictive sidecars address different targets. Dedicated temporal controls support ACF/PSD coverage without supplying missing screening records. Further metric-selection and robustness details are specified in Appendix C.3.

**Figure S6:**
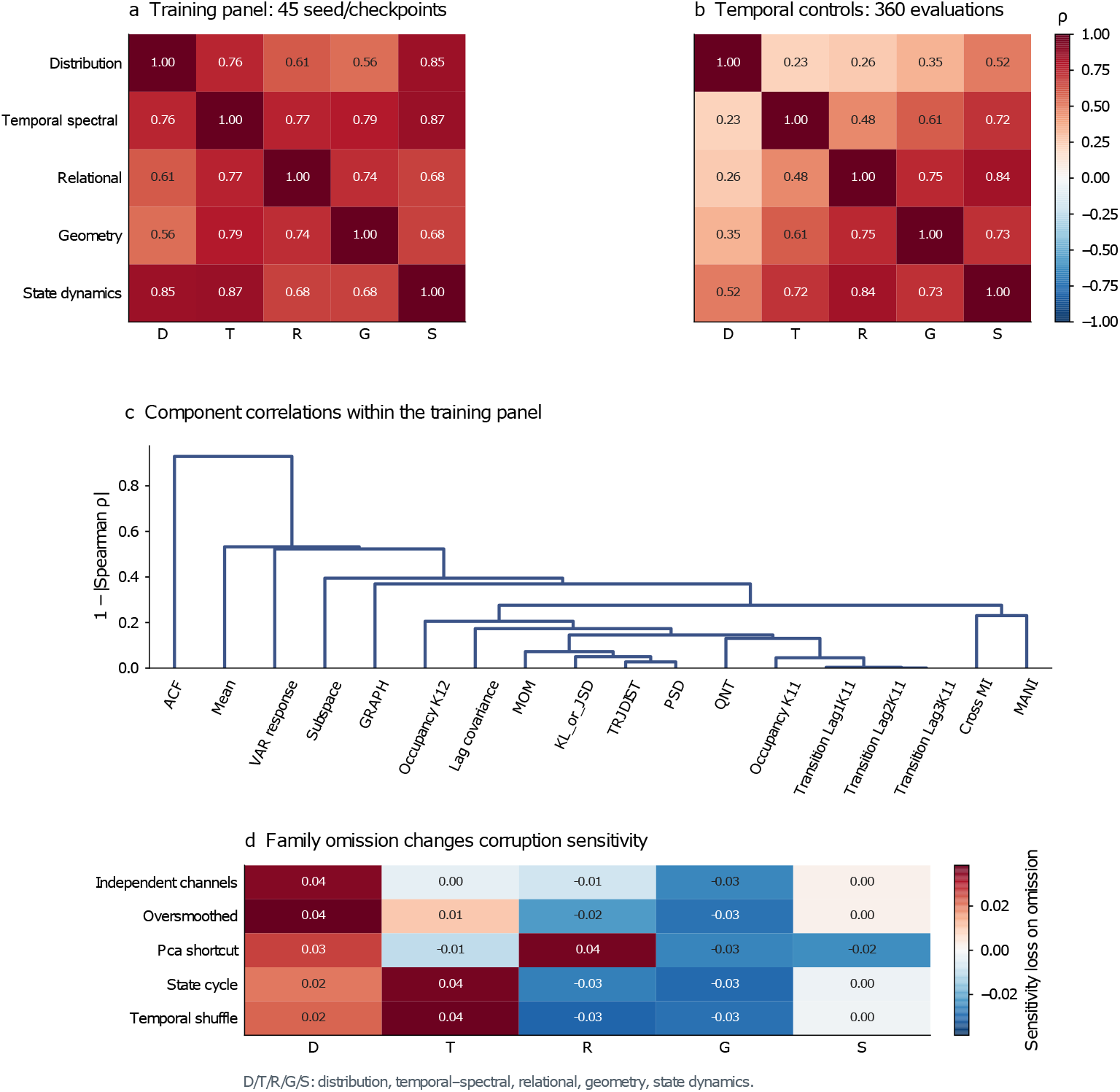
Family dependence and diagnostic complementarity. (a–b) Descriptive family correlations across 45 training evaluations and, separately, 360 temporal controls. Repeated evaluations. (c) Average-linkage clustering of 18 components using 1 *ρ* . (d) Sensitivity loss on omission, *C*_−_*_f_ C*, with remaining family weights renormalized: positive values mean that omission reduces sensitivity to the specified corruption. The heatmap shows point estimates; 200 paired sequence bootstrap records per corruption accompany the analysis. Correlation and omission effects do not establish causal unique information or downstream utility. Multiple profiles localize distinct failures even when one family reproduces a coarse model ordering. Further details on family-dependence and omission analyses are specified in Appendix C.3.

**Figure S7:**
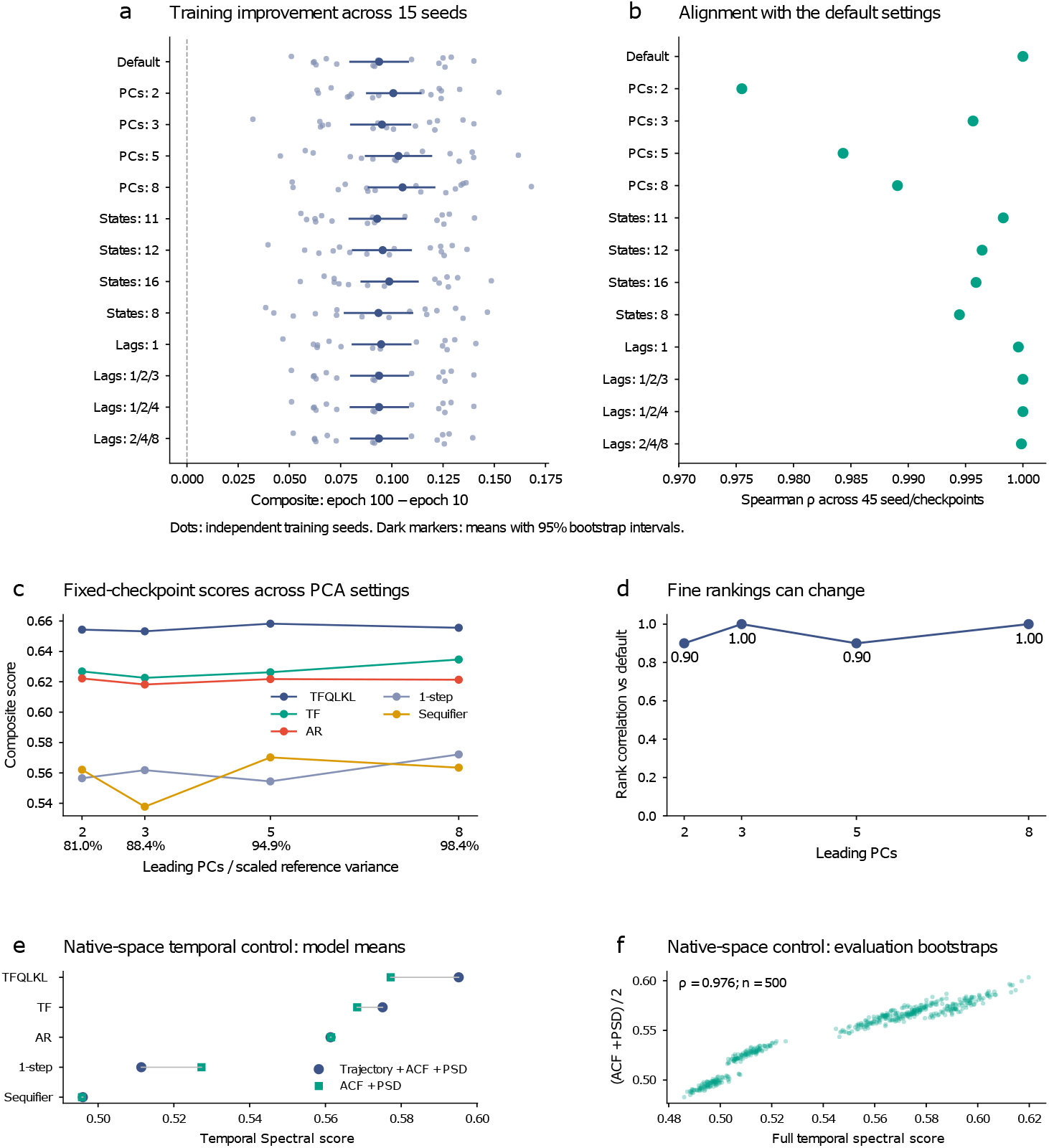
Sensitivity to benchmark parameters and temporal components. (a–b) 15 indepen-dently initialized teacher-forced, one-step Transformers (8 layers, 8 heads, width 64), trained with MAE on 16-region widefield activity and evaluated on a fixed split of 45 held-out sequences, scoring 810 forecast bins after 90 context bins. Across the default and 12 alternative PCA, state-count, and lag settings, all seeds improve from epoch 10 to 100; intervals are seed-bootstrap 95% intervals. Rank agreement with default settings is assessed across 45 seed/checkpoint evaluations (15 seeds, epochs 10, 30, and 100). (c–d) Five fixed Transformer checkpoints evaluated on one aligned 100 630 16 tensor. Varying PCA dimensionality preserves the top-ranked model but swaps 1-step and Sequifier at 2 and 5 PCs (Spearman *ρ* = 0.90–1.00). (e–f) Removing the trajectory component preserves the five-model temporal–spectral ordering. Full and ACF/PSD-only scores correlate strongly (*ρ* = 0.976) across 100 paired sequence bootstraps per model (500 evaluations). Further parameter-sensitivity details are specified in Appendix C.3.

**Figure S8:**
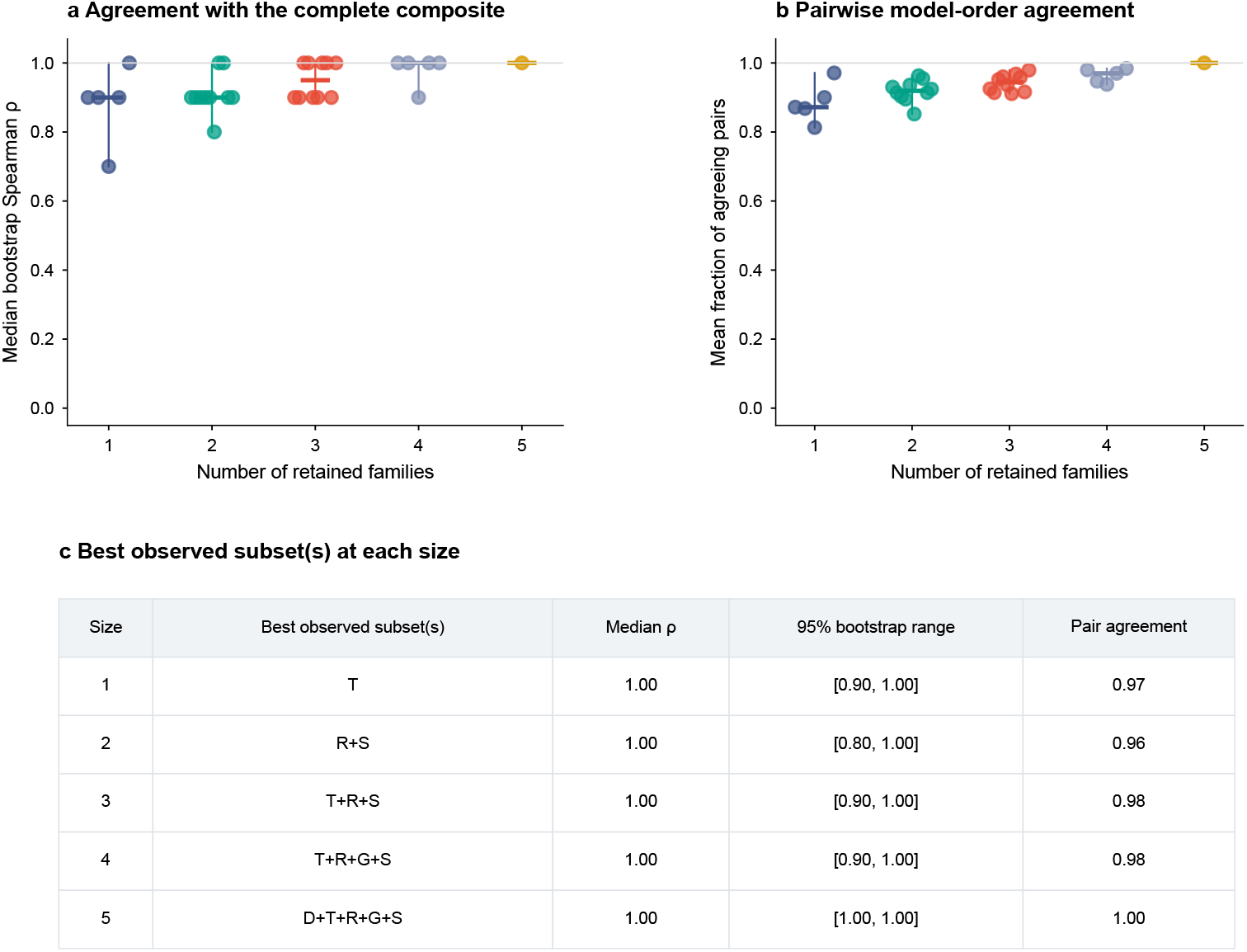
Family-subset agreement on five fixed forecasting models. All 31 nonempty subsets retain relative default weights and renormalize. Each is compared with the full composite on 100 paired evaluation-sequence bootstraps. (a) Median bootstrap rank agreement per subset. (b) Mean agreement over all ten model pairs; ties in only one ordering count as disagreement. (c) Best observed subsets selected descriptively; brackets give central 95% bootstrap ranges. The temporal family alone attains median *ρ* = 1.00 (range [0.90, 1.00]), with pairwise agreement 0.971. One family can recover coarse ordering here, while multiple family profiles remain useful for diagnostic localization. No universal minimum family count follows. Further family-subset analysis details are specified in Appendix C.3.

**Figure S9:**
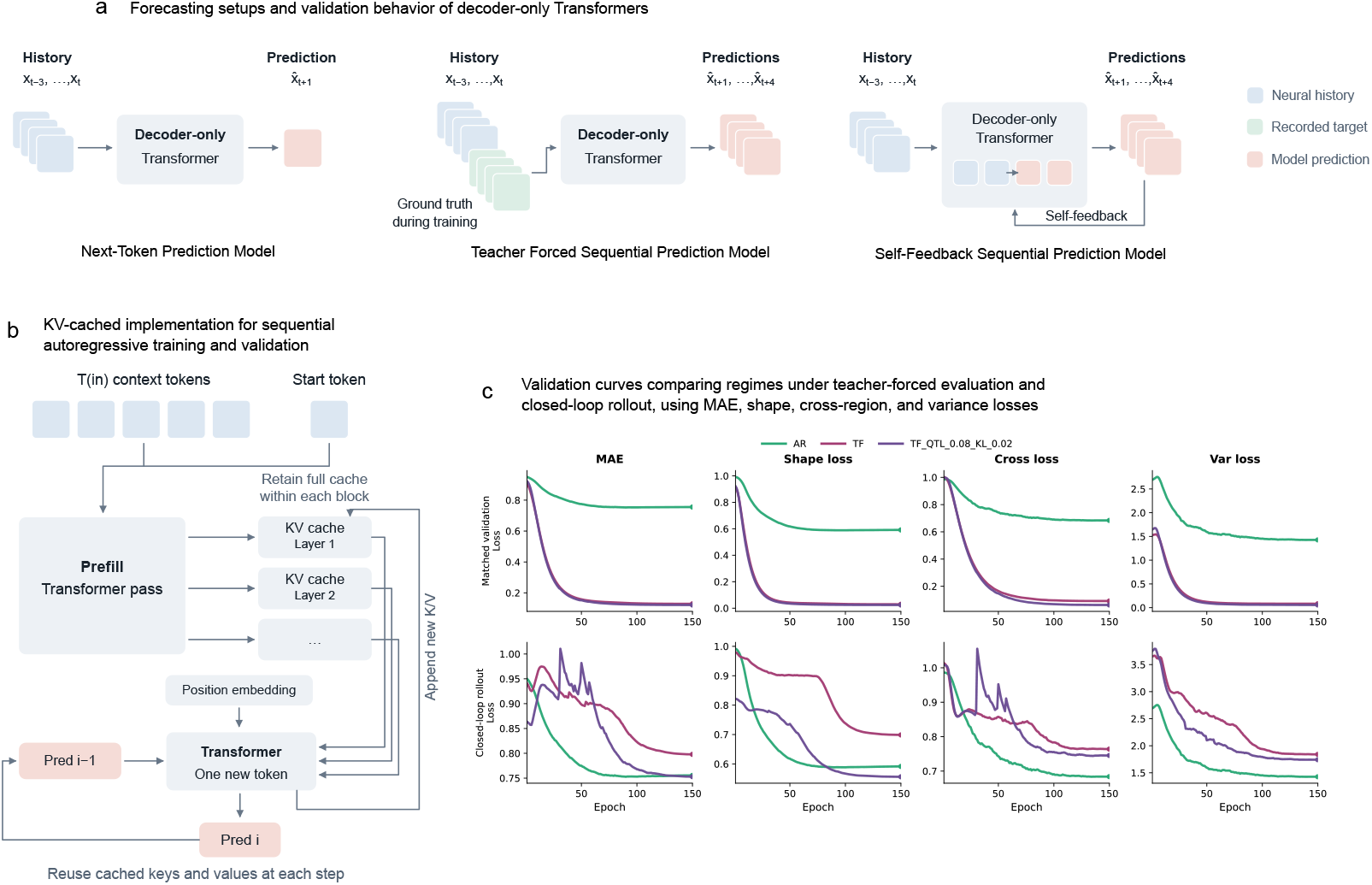
Forecasting formulations, autoregressive implementation, and validation behavior of decoder-only Transformers. **(a)** Comparison of next-token, teacher-forced sequential, and self-feedback sequential prediction. The next-token model predicts a single future timestep from the observed history; the teacher-forced model predicts a future sequence conditioned on shifted ground-truth targets; and the self-feedback model generates the sequence autoregressively by conditioning on its own previous predictions. **(b)** KV-cached implementation of self-feedback prediction. A causal prefill pass over the context and start token initializes the layer-wise key–value caches, after which only the newly generated token is processed at each step. The principal implementation retains the growing cache throughout a block (Appendix D.2). The cache resets at block boundaries. **(c)** Validation losses across training for self-feedback (AR), teacher-forced (TF), and distribution-augmented teacher-forced (TF-QL-KL) models. Rows show training-regime-matched validation (top) and common closed-loop rollout (bottom); columns report mean absolute error (MAE), temporal-shape, cross-region, and variance losses. These auxiliary diagnostics differ from the retained benchmark components. Further model and training details are specified in Appendix D.

**Figure S10:**
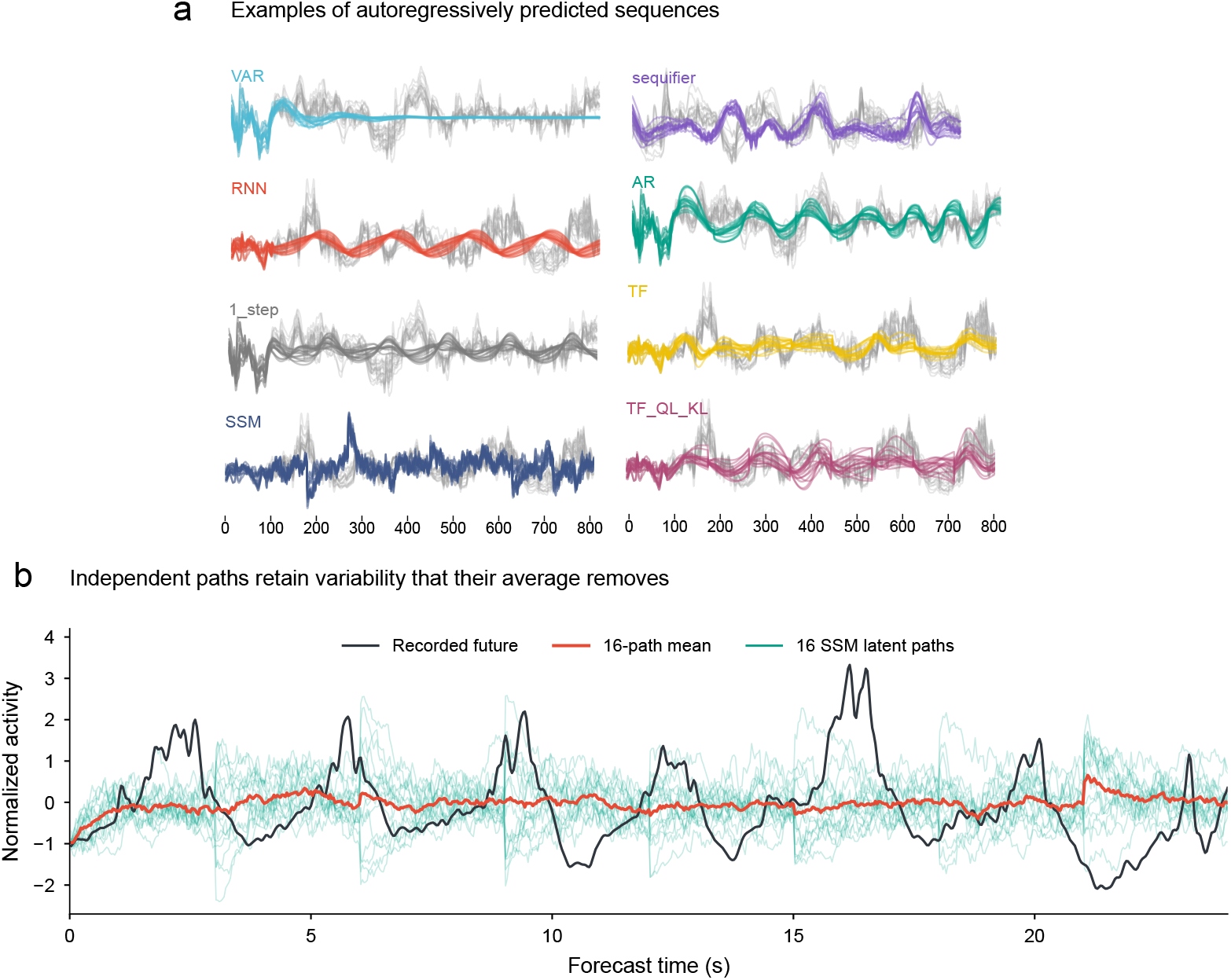
Representative forecasts and the effect of averaging stochastic trajectories. **(a)** Autoregressive rollouts from the eight forecasting models on widefield calcium activity. Model-generated regional traces are overlaid on recorded activity (light gray). Each sequence comprises 90 context samples (3 s) followed by 720 forecast samples (24 s); the horizontal axis denotes sample index. Examples illustrate differences in fluctuation amplitude, temporal regularity, and persistence across models. **(b)** Sixteen independently sampled SSM latent trajectories conditioned on the same observed history (teal), their pointwise mean (red), and the recorded future (dark gray) for one region. Individual trajectories exhibit fluctuations that occur at different times and are attenuated by averaging, yielding a smoother mean forecast. Further details on model rollouts and SSM forecasts are specified in Appendix D.

**Figure S11:**
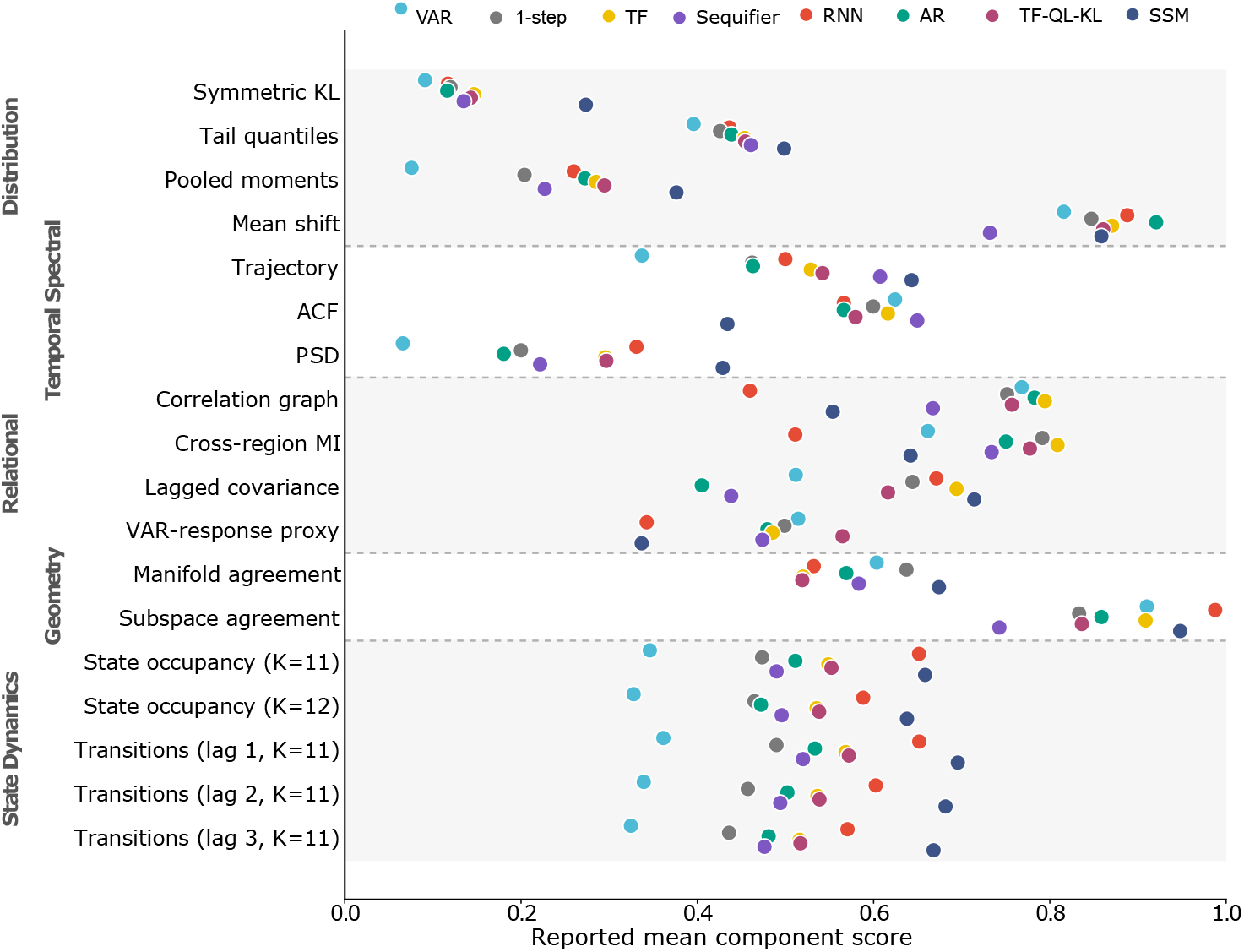
**Component-level audit of the principal widefield model panel.**Points are the reported model means; components are grouped by the five families. Further details on the widefield model comparison are specified in Appendix B.1.

**Figure S12:**
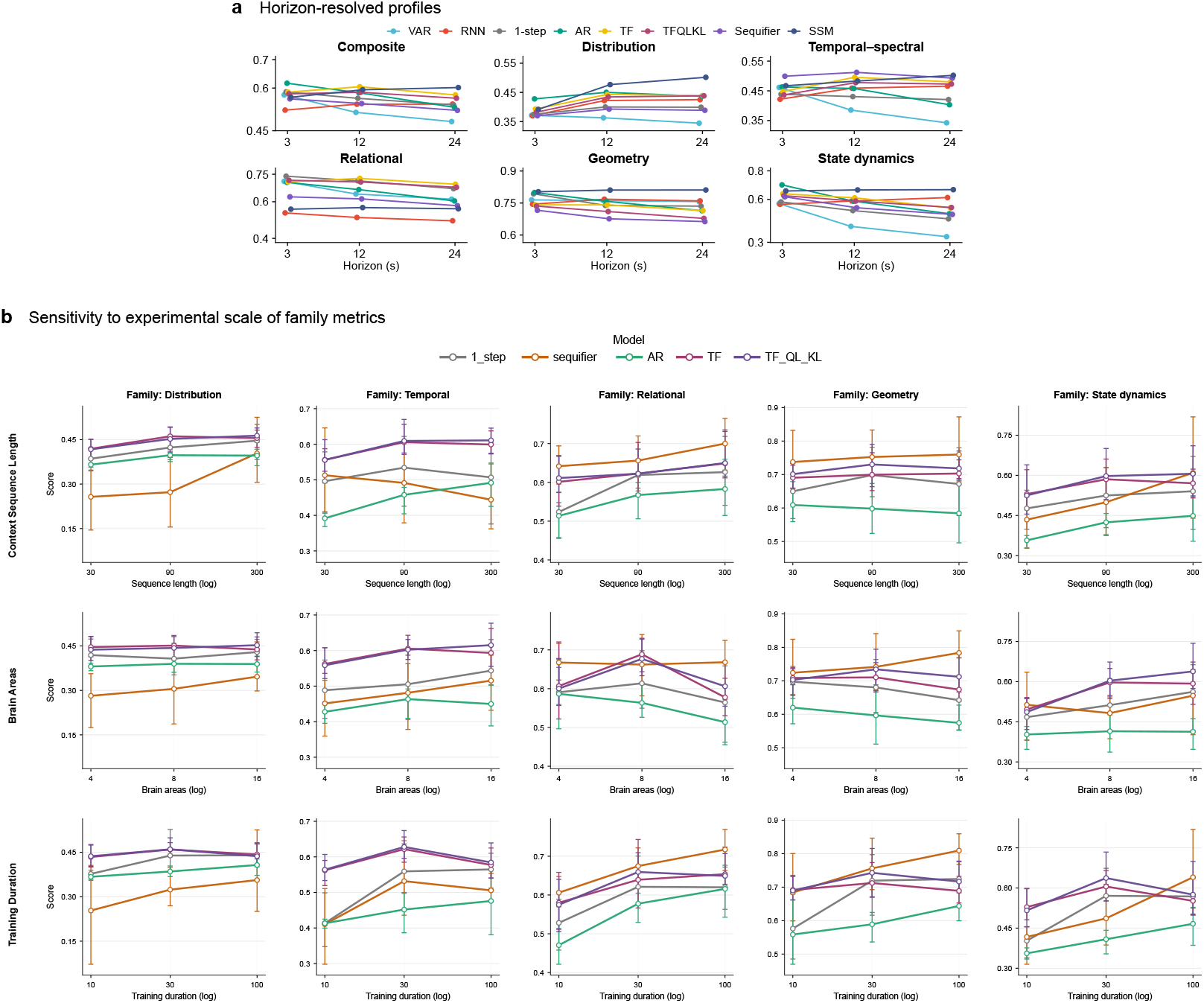
Family-level sensitivity to forecast horizon and experimental scale of the Widefield model suite. (a) Mean profiles at three forecast horizons. (b) Nethobench family scores for five Transformer configurations as functions of context length (30, 90 and 300 samples), number of areas (4, 8 and 16), and training duration (10, 30 and 100 epochs). These are descriptive trends within the evaluated configurations. Changes with increased scale are family-specific and model-dependent: some patterns are stable across axes, while others sharpen or weaken with larger temporal context, system dimensionality, or training budget. Further details on the widefield forecast horizons and evaluation support are specified in Appendix B.1.

**Figure S13:**
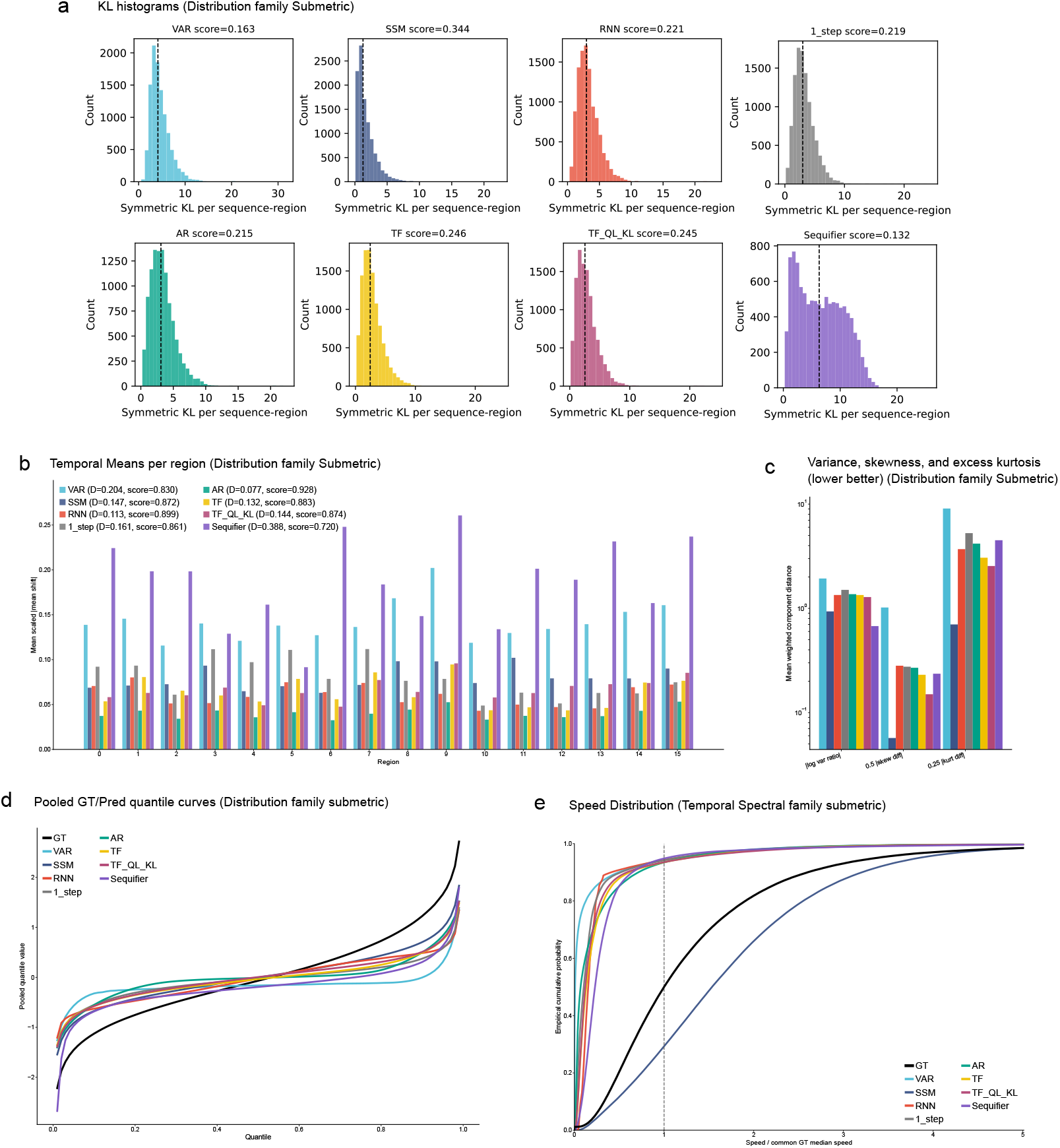
Model-specific diagnostics underlying selected Nethobench submetrics (Distribution and Temporal-spectral families). Each panel compares model predictions with matched ground-truth (GT) activity and displays component-related diagnostics. Panel annotations are diagnostic summaries and need not equal the retained aggregations in Table 10. **(a)** Distributions of symmetric KL divergence across sequences and regions. **(b)** IQR-scaled absolute temporal-mean shifts across regions. **(c)** Weighted diagnostic discrepancies in variance, skewness, and excess kurtosis; the retained pooled-moment score is defined in Appendix C.1. **(d)** Pooled GT and predicted quantile functions. **(e)** Empirical cumulative distributions of speed in the GT-defined latent space, normalized by the common GT median. Together, these diagnostics localize the statistical and dynamical differences summarized by the Nethobench Distribution and Temporal-spectral families. Further model and metric details are specified in Appendices D and C.1.

**Figure S14:**
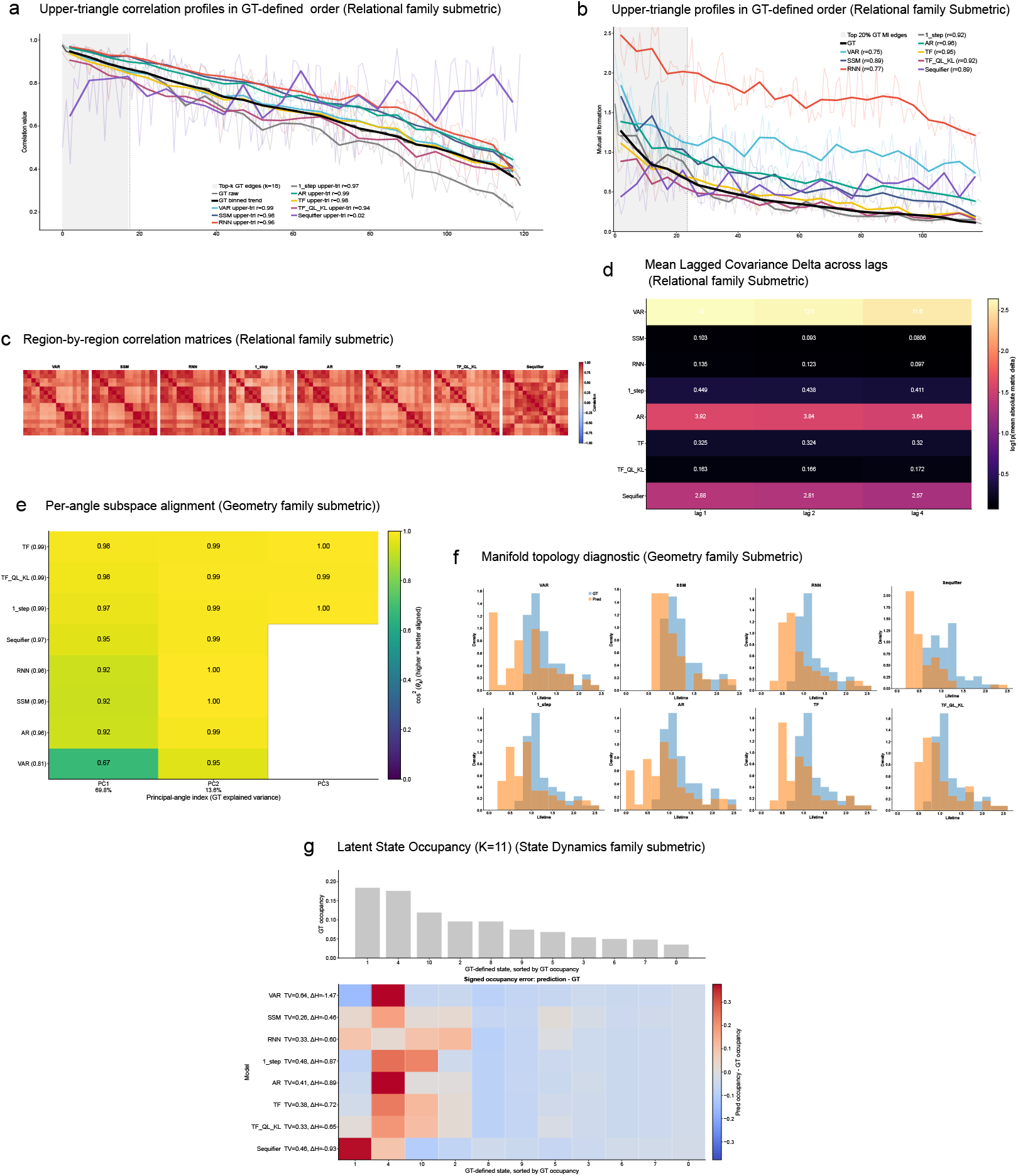
Model-specific diagnostics underlying selected Nethobench submetrics (Relational, Geometry and State dynamics families). Each panel compares model predictions with matched ground-truth (GT) activity and displays component-related diagnostics. Panel annotations are diagnostic summaries and need not equal the retained aggregations in Table 10. **(a)** Region-pair correlations ordered by GT edge strength. **(b)** Mutual-information profiles for the top 20% of GT edges, shown in GT-defined order. **(c)** Region-by-region correlation matrices. **(d)** Mean absolute lagged-covariance discrepancies at lags 1, 2, and 4. **(e)** Per-dimension subspace alignment, measured by the squared cosine of each principal angle. **(f)** Persistence-lifetime distributions for GT and predicted manifolds. **(g)** GT occupancy across *K* = 11 latent states and signed model occupancy errors in the same GT-defined state space. Together, these diagnostics localize the statistical and dynamical differences summarized by the Nethobench Relational, Geometry and State dynamics families. Further model and metric details are specified in Appendices D and C.1.

**Figure S15:**
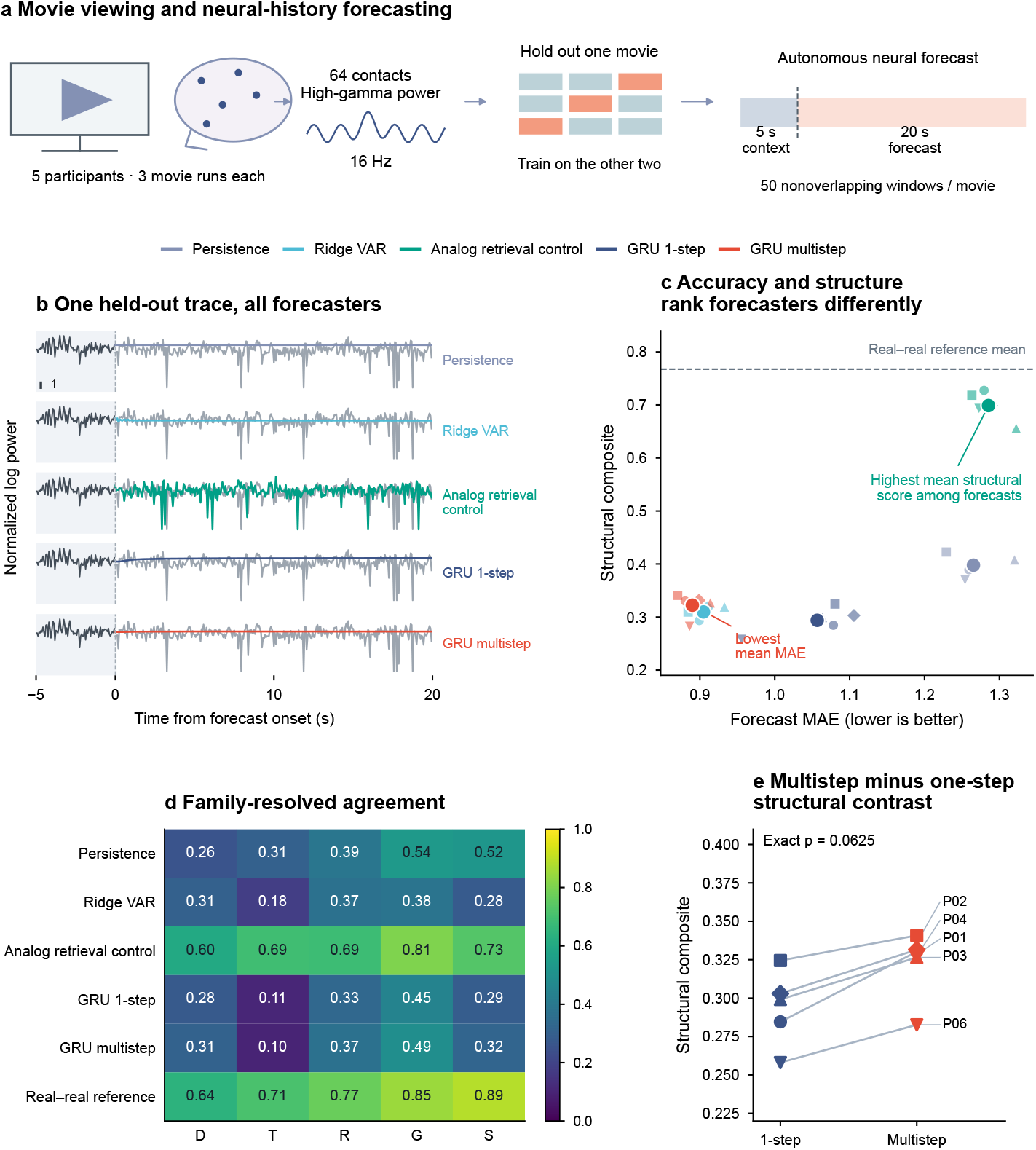
Brain Treebank human neural forecasts separate structural agreement from pointwise accuracy. (a) Five participants, high-gamma activity during naturalistic movie viewing (three movie runs per participant) and 64 contacts per participant, 5-s context and 20-s forecast. The task/contact drawings are illustrative. (b) A fixed first-participant/run/window/contact example, without smoothing. (c–d) Participant means and family profiles: training-only analog retrieval leads structural agreement, whereas multistep GRU has the lowest mean MAE. The real–real line is an empirical reference. (e) Paired GRU participant means: Δ*C* = 0.0285, hierarchical 95% interval [0.0180, 0.0415], exact two-sided *p* = 0.0625. Structural advantage does not establish conditional correctness, mechanisms or downstream utility. Retrieval exclusions and preprocessing boundaries are detailed in Appendices B.3 and B.6.

**Figure S16:**
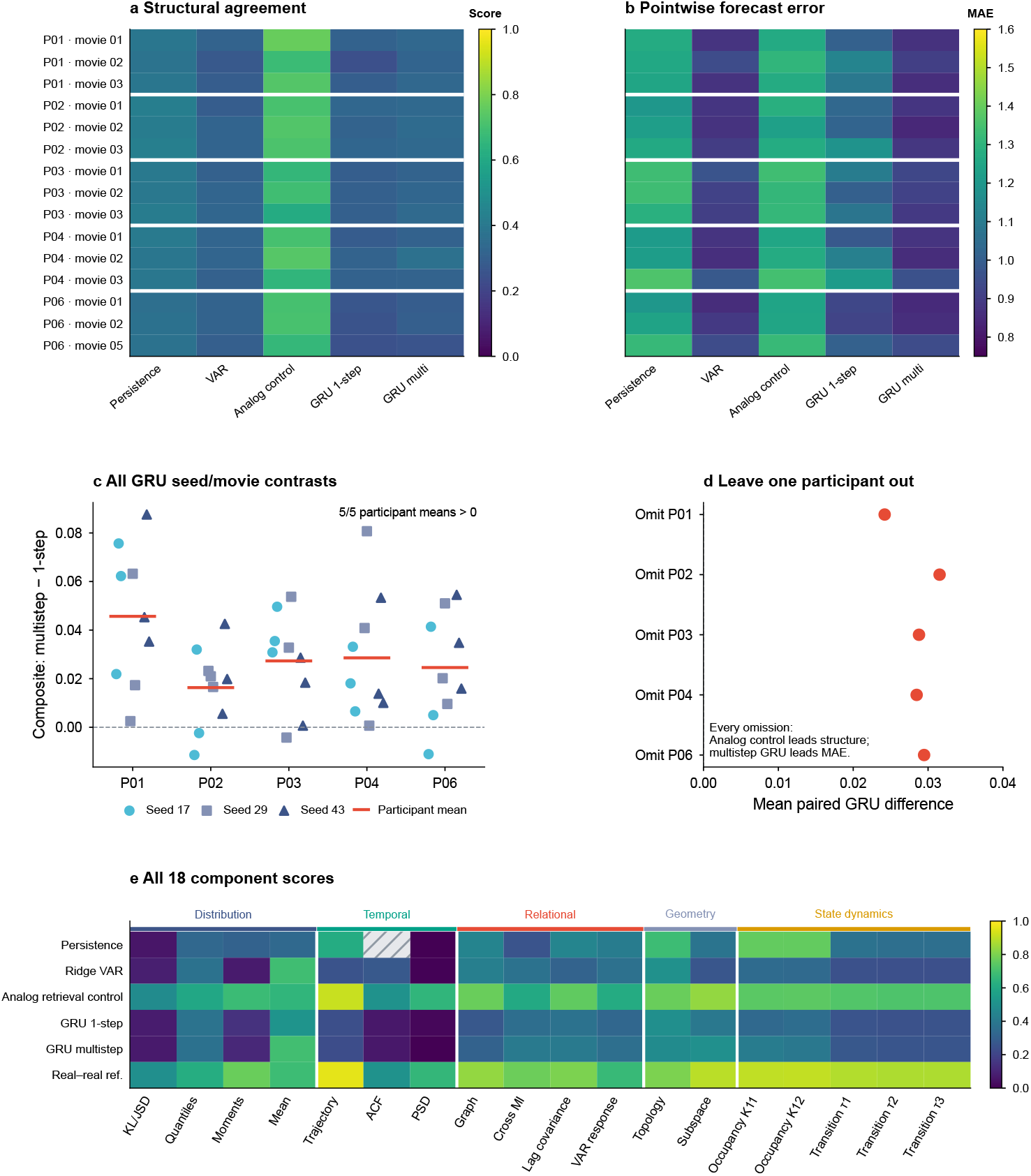
Human forecast robustness by movie, seed, participant and component (Brain Treebank). (a–b) All 15 held-out folds, averaging GRU seeds; 135 model/fold/seed evaluations underlie the summaries. Analog retrieval leads structure in every fold; multistep GRU leads MAE in 13 and VAR in two. (c) All 45 paired GRU seed/movie contrasts, including negative values. (d) Leave-one-participant-out contrasts. (e) All 18 components and the real-real reference. Persistence ACF is undefined; excluding ACF from every method preserves ordering and a positive mean GRU contrast (0.0280). The independent sample size is five participants, not 15 folds or 45 contrasts. Further details on Brain Treebank splits, scoring, and uncertainty are specified in Appendix B.3.

**Figure S17:**
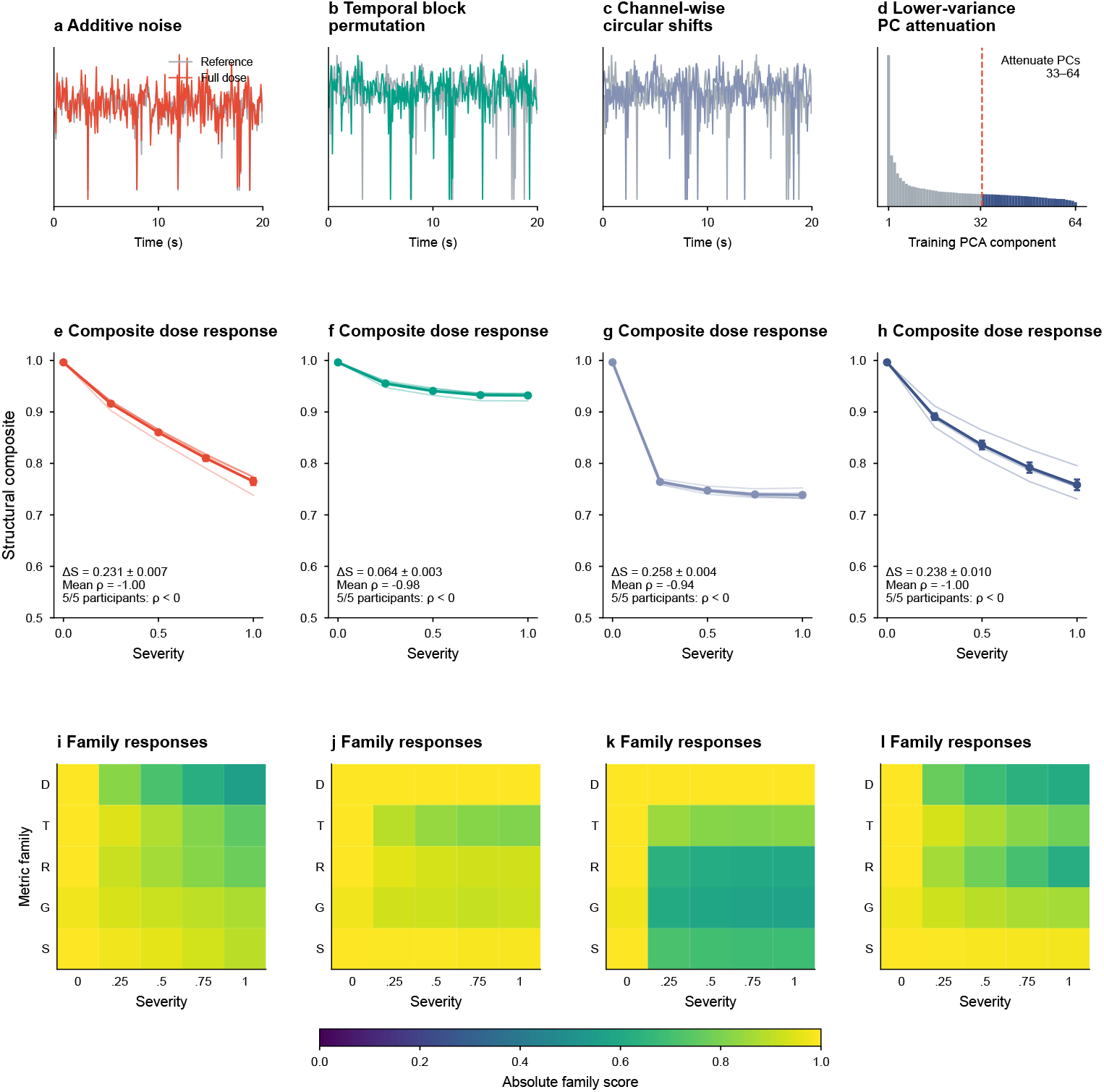
Controlled perturbations of human neural activity from Brain Treebank. (a–c) Full-dose noise, 2-s block permutation and channel shifts applied to a fixed held-out trace. (d) Training-PC spectrum; attenuation affects PCs 33–64, carrying 21.3–40.3% of training variance across folds. (e–h) Participant dose curves and mean participant SEM across all 300 evaluations. Full-dose drops are 0.231, 0.064, 0.258 and 0.238. All 20 participant/type dose correlations are negative, without implying strict monotonicity at every step. (i–l) Family profiles on common scales. Block permutation preserves samples and within-block paths. These controls do not establish preservation of arbitrarily small-variance or task-specific information. Further details on Brain Treebank perturbations and scoring are specified in Appendix B.3.

**Figure S18:**
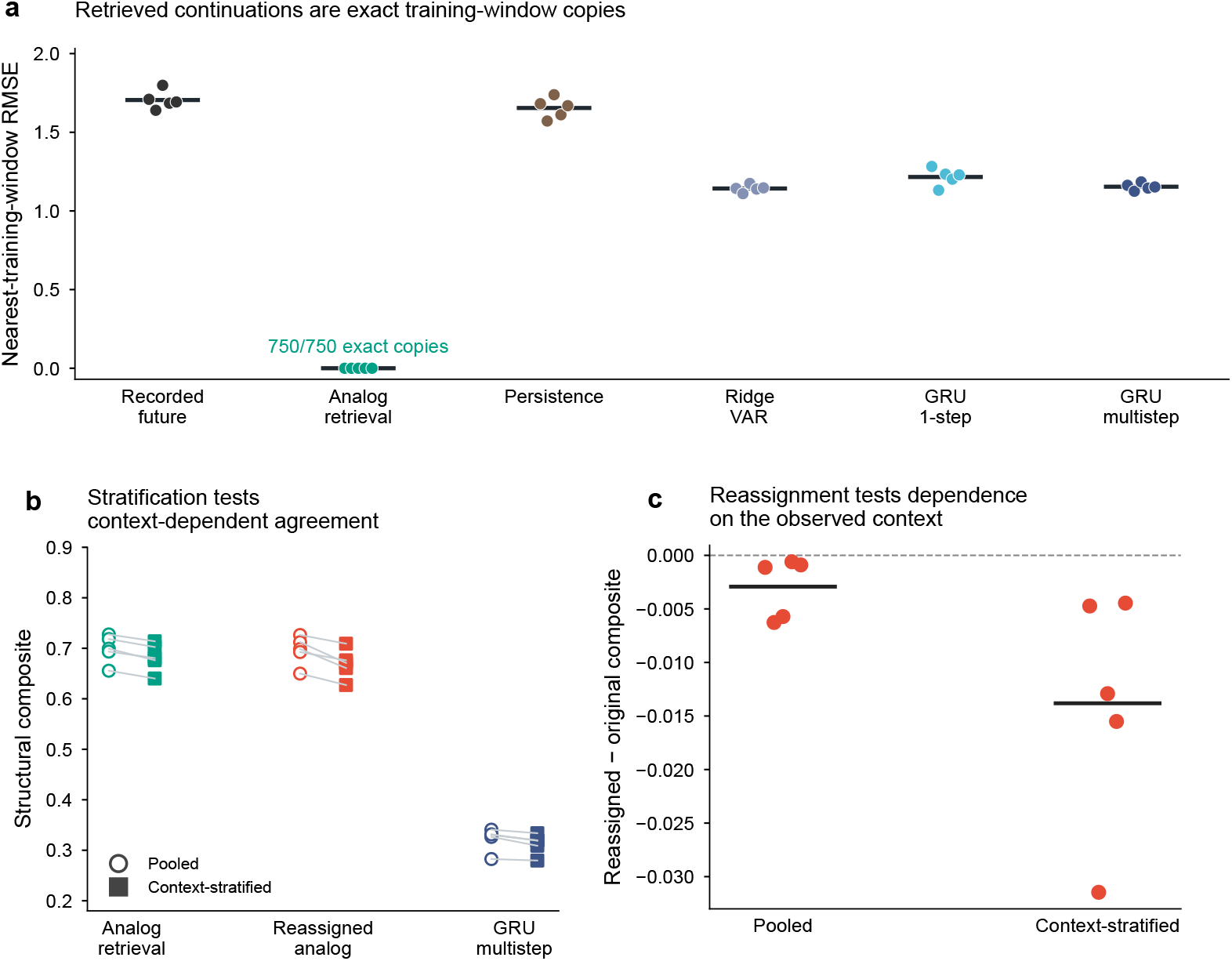
Training-window provenance and context-stratified structural evaluation. **(a)** Nearest-training-continuation RMSE, computed over the full forecast window without alignment or amplitude fitting. All 750 analog-retrieval outputs exactly reproduce eligible training continuations; no other evaluated forecast or recorded future does. **(b)** Structural composites scored over all windows together (pooled; circles) or separately within two observed-context groups and then averaged (context-stratified; squares). Groups contain 25 windows each, sorted by the first PC of per-contact context means and log standard deviations; standardization and PCA are fitted on training contexts only. Reassigned retrieval swaps the same continuations between groups by shifting this order by 25 positions. **(c)** Change in retrieval composite after reassignment, corresponding to the reassigned-minus-original differences in (b). Mean decreases are 0.003 for pooled and 0.014 for context-stratified scoring. Points represent participants, averaging seeds within movies and movies within participants. Black bars indicate participant means; lines in (b) connect paired values. Further implementation details are given in Appendix B.6.

**Figure S19:**
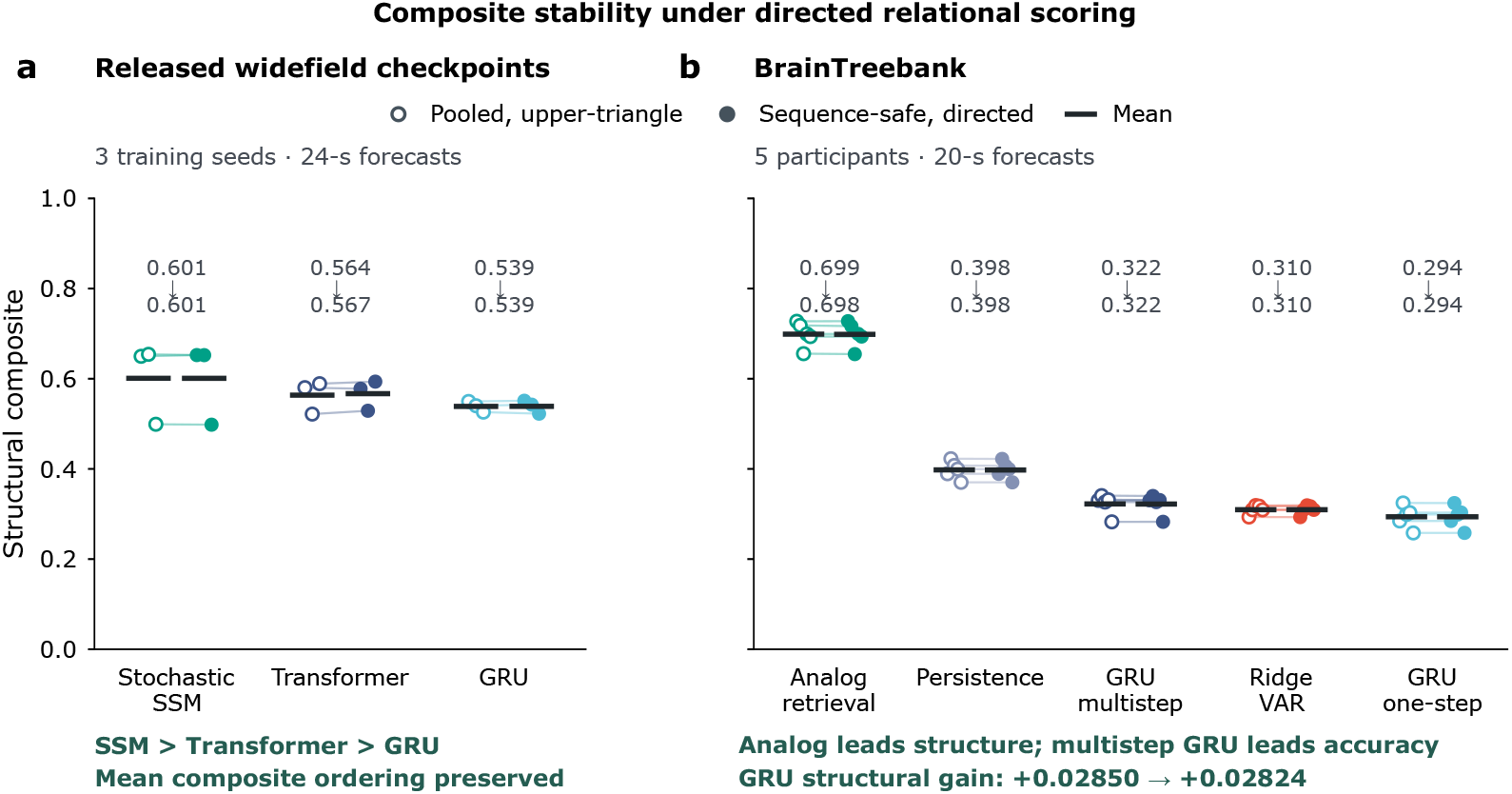
Composite findings persist under sequence-safe, directed relational scoring. Frozen forecasts were rescored using lag pairs restricted to their original sequences and both directed off-diagonal orientations in the lagged-covariance and ridge VAR(1) comparisons. Scaling, similarity transforms, all other components and aggregation weights were unchanged. Open circles show pooled, upper-triangle scoring, which flattens sequences and compares one off-diagonal matrix triangle; filled circles show sequence-safe, directed scoring, which retains within-sequence lag pairs and both off-diagonal orientations; lines pair the same evaluation unit, and black bars mark means. Numerical annotations give the pooled, upper-triangle mean followed by the sequence-safe, directed mean. ** (a) Released widefield checkpoints: three training seeds per model, evaluated on 58 validation sequences with 3-s context and 24-s forecasts. Each SSM checkpoint averages the scores of four fixed latent draws. Mean composite ordering remains SSM *>* Transformer *>* GRU: 0.60097 → 0.60094, 0.56362 → 0.56682 and 0.53856 → 0.53873, respectively. (b) BrainTreebank: five participant means, averaging training seeds within held-out movie and then three movies within participant (135 model/fold/seed evaluations; 5-s context and 20-s forecasts). The complete mean composite ordering is preserved. Analog retrieval remains highest in structural agreement (0.69878 → 0.69837); multistep GRU retains the lowest MAE (0.88990), which is unaffected by rescoring. Its composite advantage over one-step GRU changes from 0.02850 to 0.02824 and remains positive in all five participants. The structural and accuracy winners also persist in every leave-one-participant-out summary. Points represent training seeds in (a) and participants in (b); bars are means, not uncertainty intervals. These findings establish stability of the reported composite conclusions in the tested panels. Further details on the relational-scoring implementation are specified in Appendix C.6.

**Figure S20:**
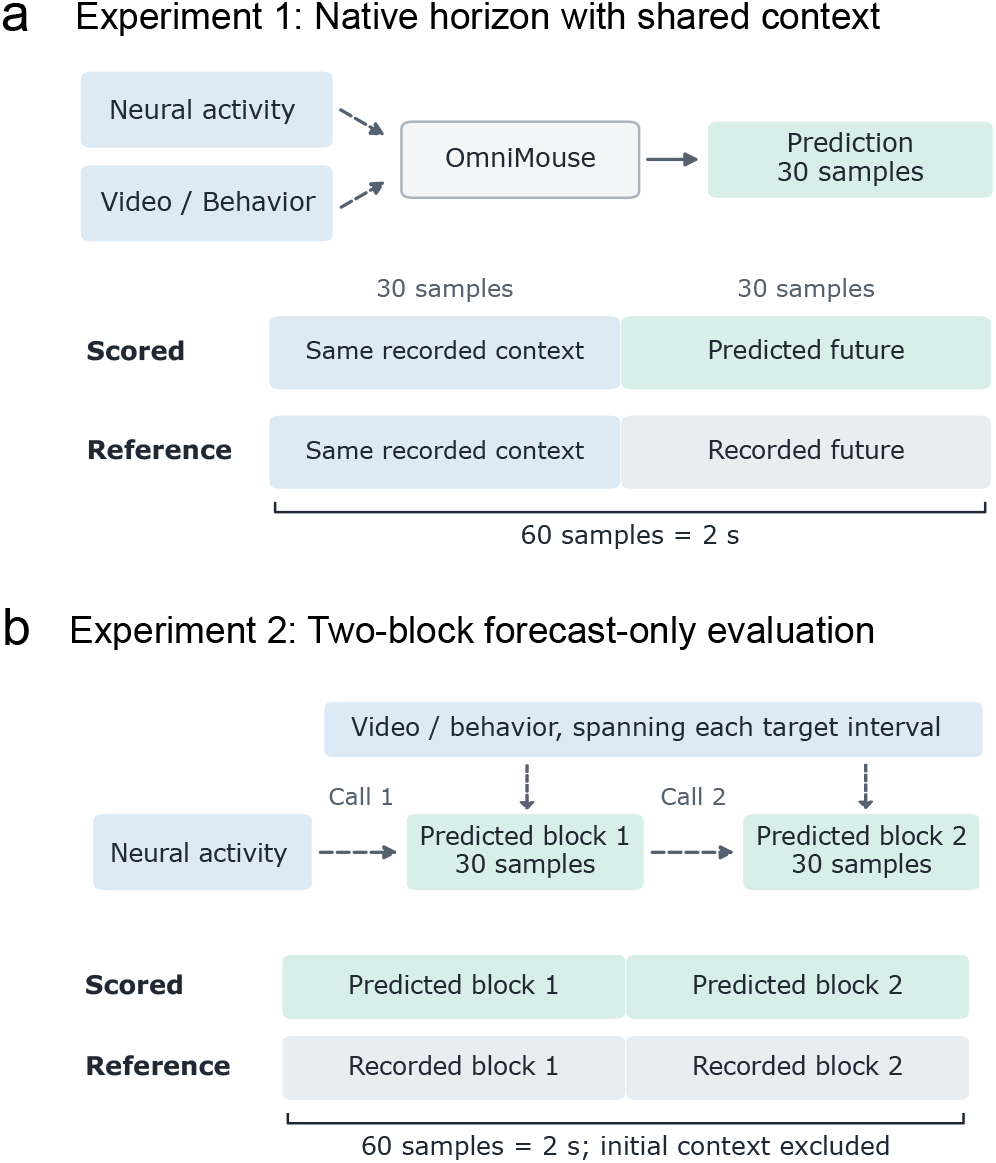
OmniMouse input and scoring protocols. (a) Native-horizon evaluation concatenates 30 recorded context samples with 30 predicted neural samples and compares this sequence with the same recorded context followed by the recorded target. This shared context contributes to absolute agreement. Configurations without neural history receive no neural response values as model input. (b) Prediction-only evaluation scores two successive 30-sample predicted blocks against the corresponding recorded activity, excluding initial recorded context. For neural-history configurations, the second call encodes the first 25 predictions from the preceding block. In both protocols, enabled video and behavioral inputs include observations from the target interval. The complete video clip enters the encoder before any strategy-specific feature selection. Further details on input timing, covariates, and scoring support are specified in Appendix B.2.

**Figure S21:**
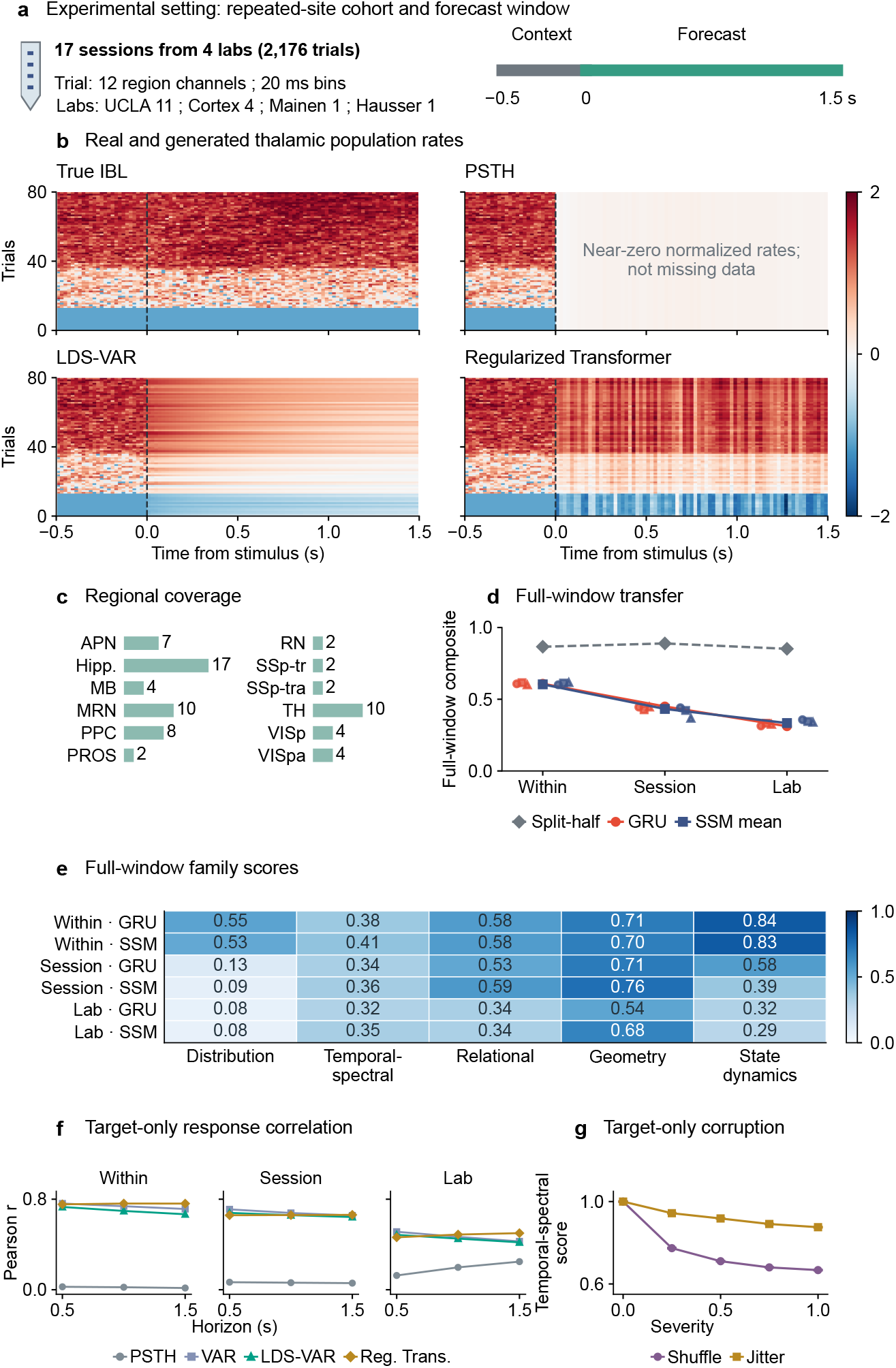
IBL Neuropixels population forecasts and transfer: structural agreement declines under transfer. (a) Seventeen sessions, four laboratories and 2176 trials. (b) Identically ordered regional-rate maps; white PSTH entries are finite near-zero normalized rates. (c) Regional coverage requires five quality-filtered units; missing channels are zero-filled before normalization. (d–e) Full-window transfer composites and profiles, including context. Small markers show three training seeds; connected points score ensemble-mean predictions. (f) Target-only Pearson correlations by horizon. (g) Target-only order controls on 436 trials at 50 Hz; bars show SD over three corruption realizations. Shuffle severity controls selected positions, and maximum channel jitter is nine bins. Sampling supports, controls and model identities are detailed in Appendix B.4.2.

**Figure S22:**
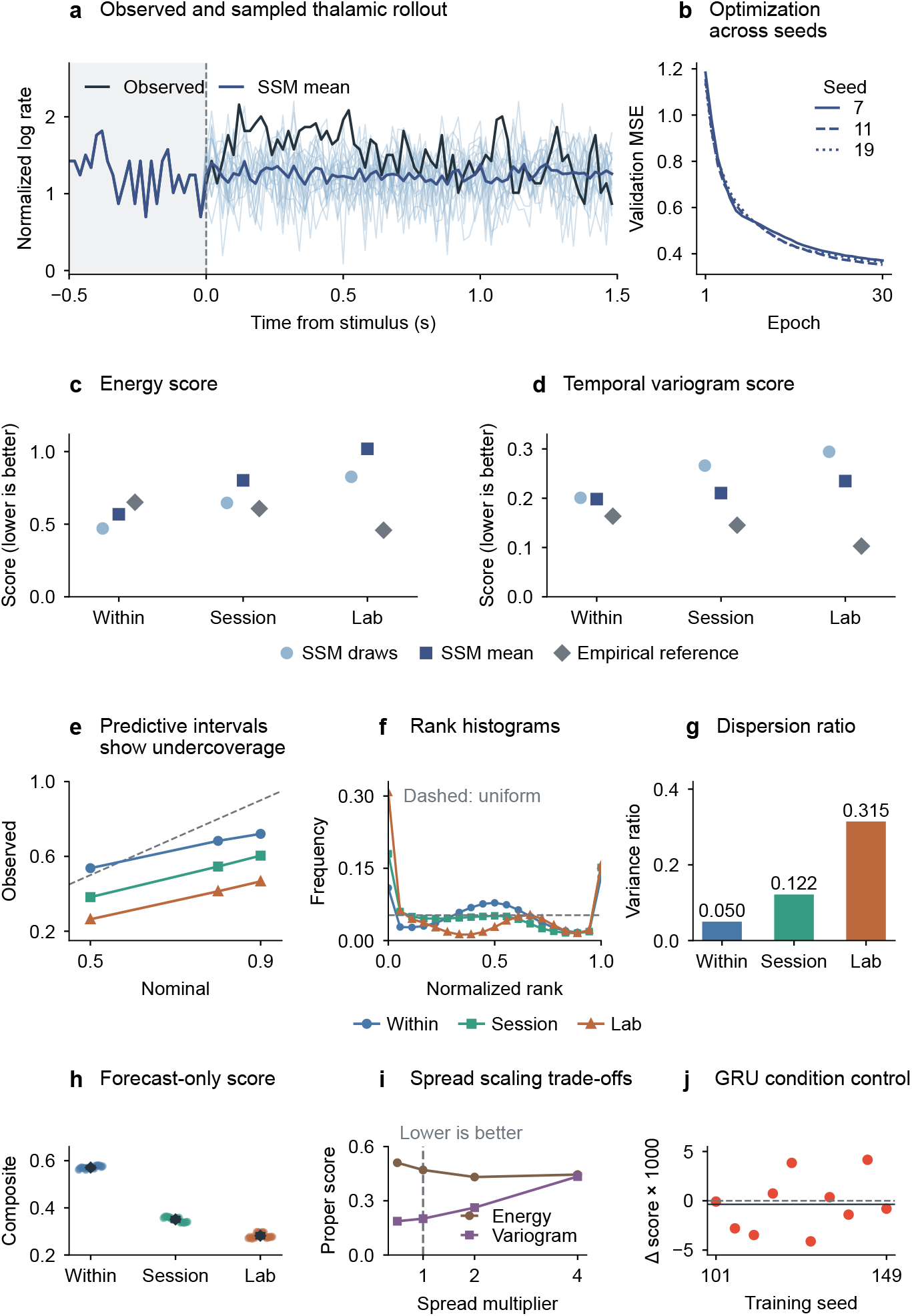
Stochastic IBL Neuropixels forecasts, calibration and conditioning controls. (a–b) First stored example and three SSM training histories (75 target bins 12 regions). (c–d) Target-only energy and temporal variogram scores for 18 draws, their mean, and other-real-target reference ensembles. Draws improve energy versus the mean; variogram errors exceed the reference. (e–g) Coverage, ensemble ranks and variance ratios; pooled target variance includes between-context variation. (h) Eighteen structural Monte Carlo replicates (mean SD). (i) Spread sensitivity in each metric’s own units. (j) Ten paired GRU conditioning seeds: correct-minus-shuffled mean composite 0.000358, exact *p* = 0.6914. These covariates and models do not establish a general absence of conditioning benefits. Further details on probabilistic evaluation and the conditioning control are specified in Appendix B.4.2.

**Figure S23:**
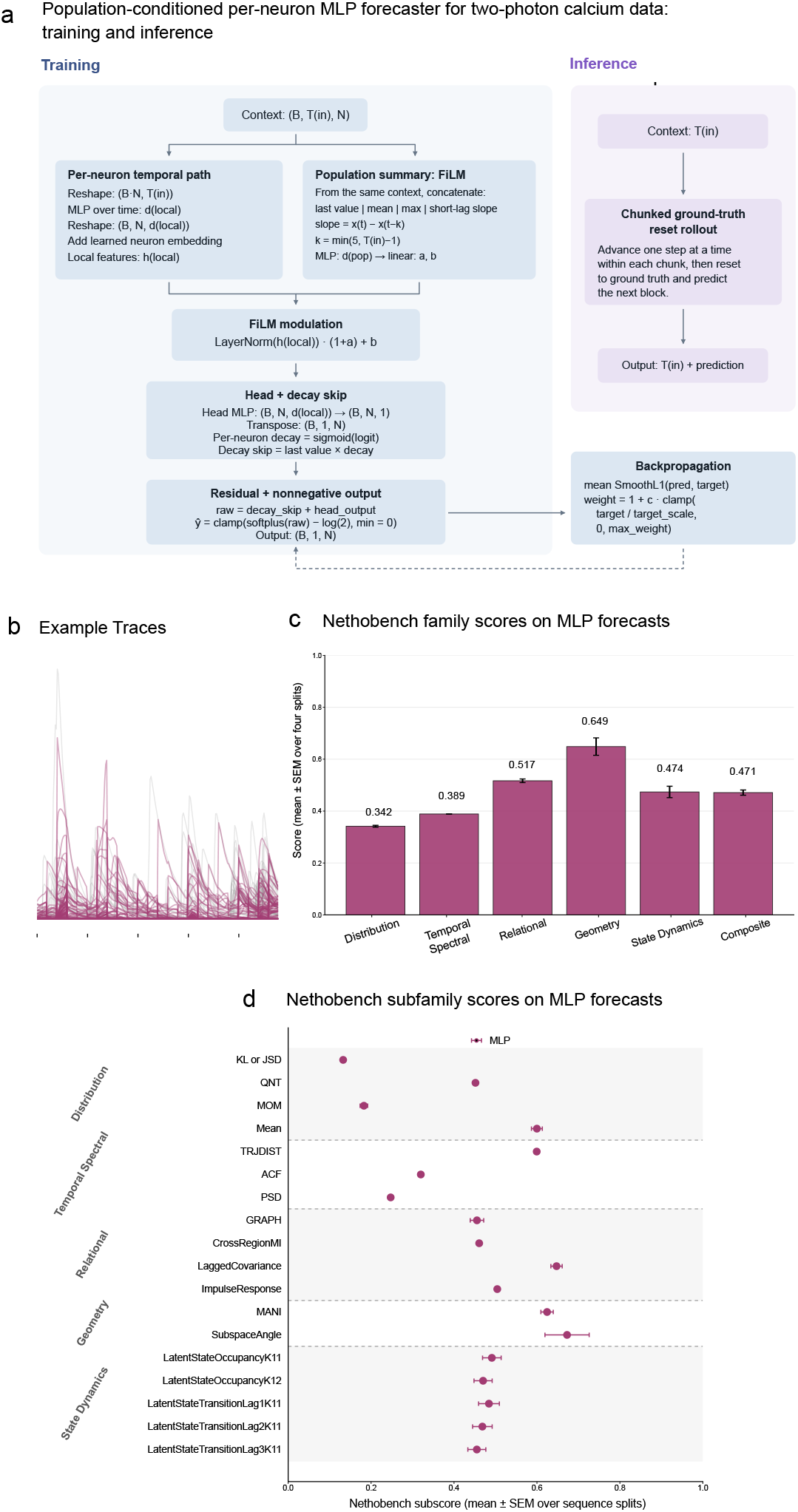
Two-photon MLP baseline and Nethobench evaluation. **(a)** Schematic of the MLP forecaster, which combines per-neuron temporal features with a population-level context embedding and predicts the next calcium-activity sample through a residual head plus learned persistence skip. Chunked inference: the one-step predictor advances autoregressively within each chunk, then the context is reset to recorded activity before the next chunk. This is not a continuously autonomous long-horizon forecast. **(b)** Example calcium-activity traces. **(c)** Family-level Nethobench scores and composite (0.471), with mean SEM across four sequence splits. Geometry (0.649) exceeds distributional (0.342) and temporal-spectral (0.389) agreement. **(d)** Nethobench submetric profile across all five families. Split uncertainty is within one mouse; it does not estimate between-animal variation. Further details on the two-photon recording and model protocol are specified in Appendix B.5.

**Figure S24:**
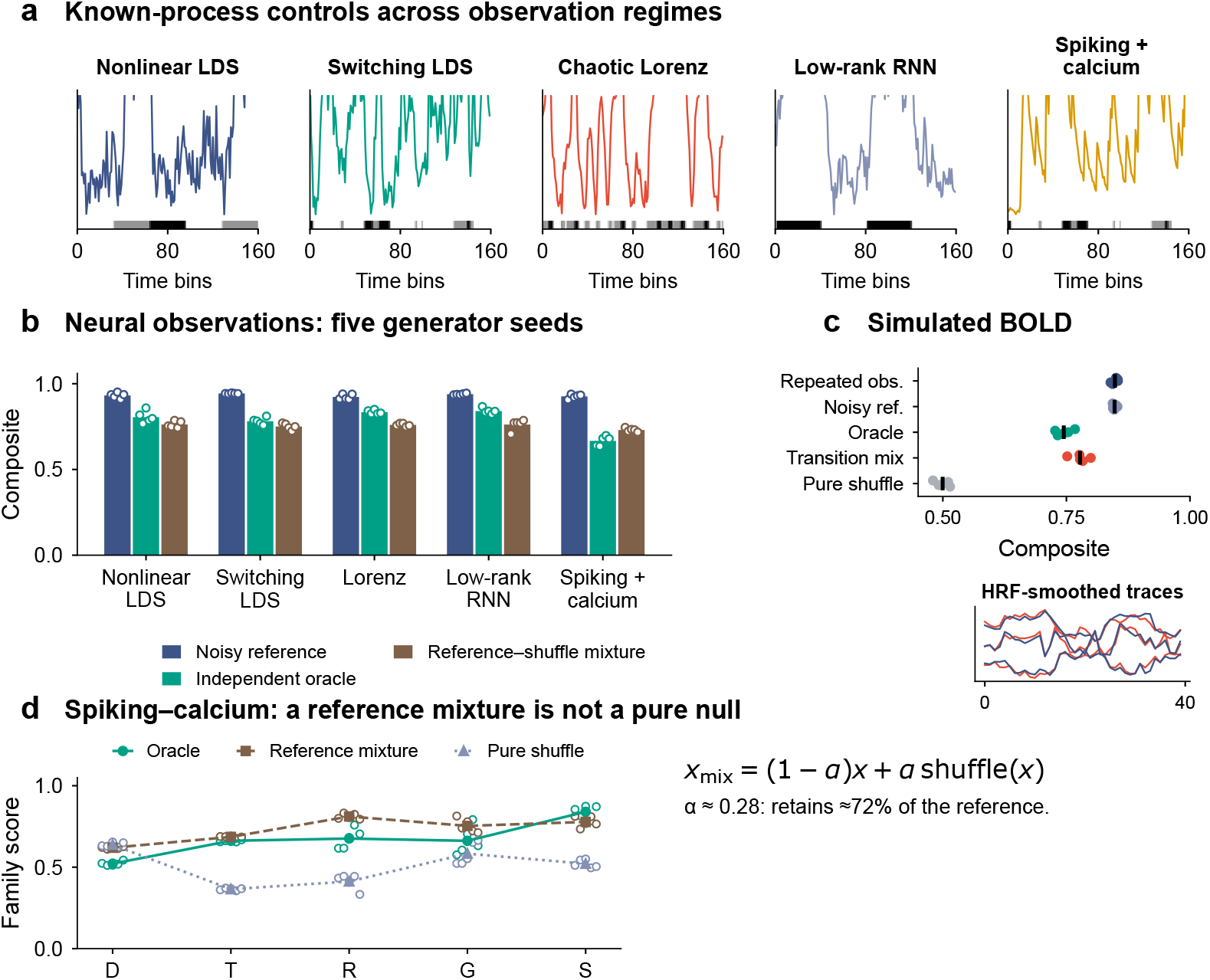
Structural validation across richer synthetic systems and simulated BOLD. (a) Display-scaled available trace segments and latent/task-state strips from five known-process systems. (b) Five generator seeds, all 18 retained components: noisy reference, independent oracle, and reference–shuffle mixture. Bars are means; points retain individual seeds. The noisy reference is not an independent real–real half-split. (c) Simulated BOLD controls average three systems within each seed; 40-bin support provides 17/18 components because quantiles are unavailable. The inset retains the reference-like (blue) and corrupted (red) HRF-smoothed trace examples. (d) Spiking–calcium profiles distinguish a mixture retaining approximately 72% of the reference from a pure shuffle. The independent oracle exceeds the pure shuffle in all five seeds. D/T/R/G/S denote distributional, temporal-spectral, relational, geometric and state dynamics families. The mixture’s closeness to the reference and its family profile explain its higher composite. Controls do not establish mechanistic validity or real-fMRI performance. Human calibration is reported separately in Figure S17. Further details on the synthetic and simulated-BOLD validation protocols are specified in Appendix C.2.

**Figure S25:**
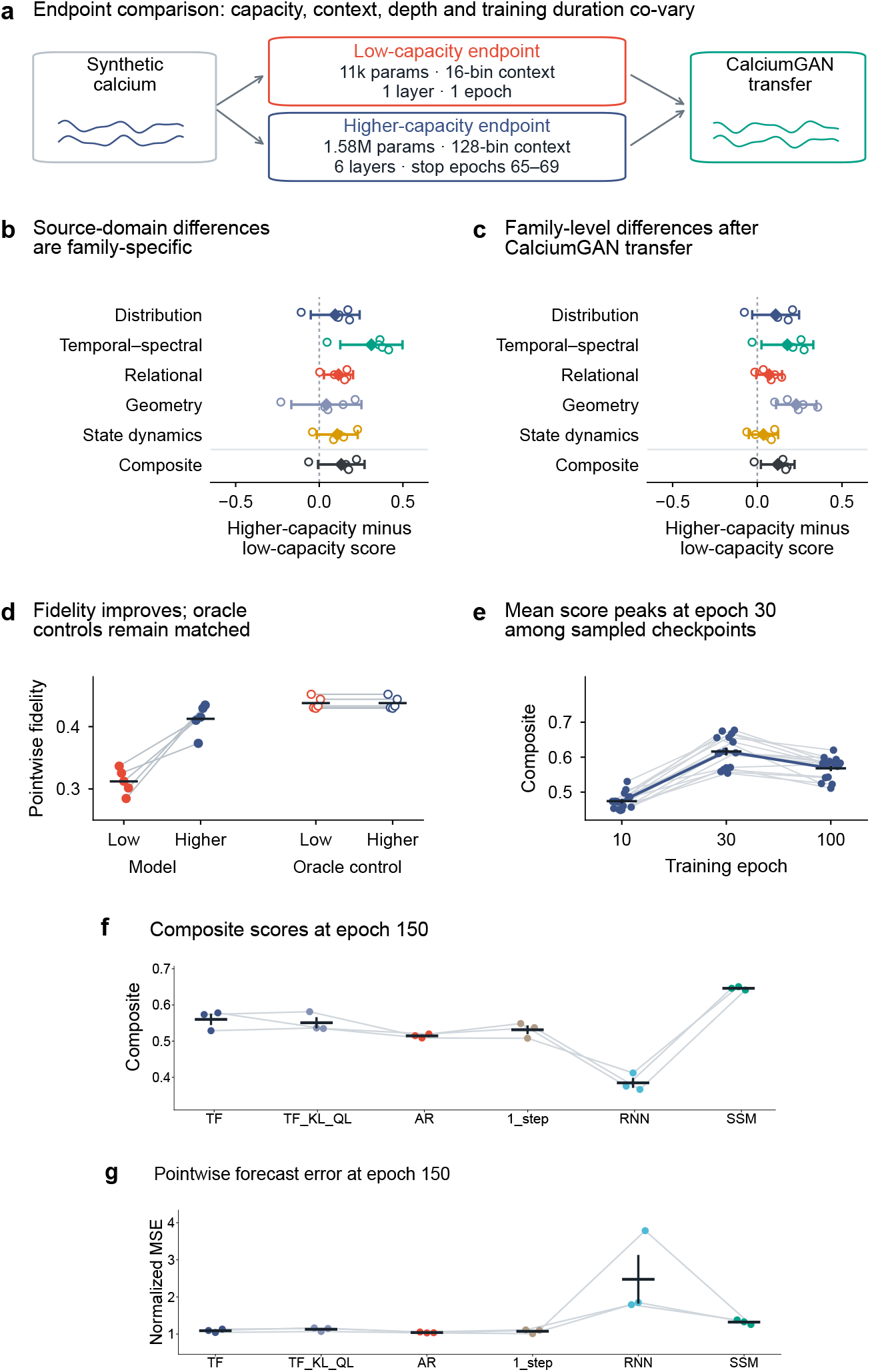
Nethobench scores differ across model endpoints, training stages and model families. (a) Endpoint settings co-vary in capacity, context, depth and training; traces are schematic. (b–c) Five paired source/CalciumGAN-transfer differences on matched generated targets. (d) Source-domain pointwise-fidelity controls (the same five paired seeds as b–c; filled/open markers show model/oracle scores). (e) Fifteen paired training seeds: epoch 100 exceeds 10 in all seeds, whereas 30 exceeds 100 in 14/15 (Holm-adjusted *p* = 0.000183). (f–g) Six model families evaluated at epoch 150 on the same validation sequences: (f) composite score and (g) normalized forecast MSE. Points show three training seeds, lines connect matched seeds, and bars show seed mean SEM. Model capacity and compute are not matched; CalciumGAN is a generated domain. Further details on endpoint comparisons and training trajectories are specified in Appendix C.4.

**Figure S26:**
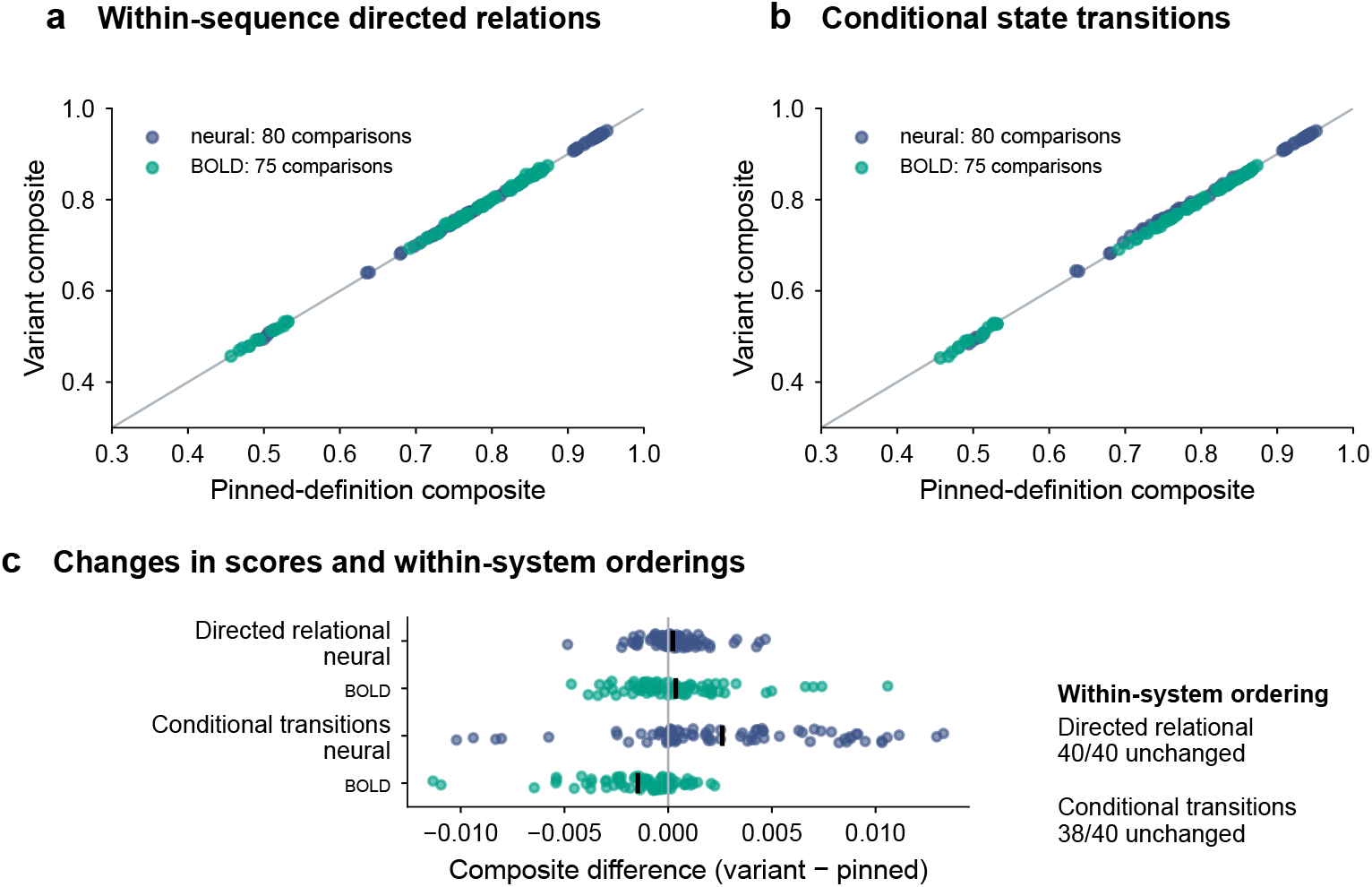
Implementation sensitivity on fixed known-process controls. (a) Sequence-safe lagged covariance/VAR comparisons using all directed off-diagonal entries. (b) Row-conditional state-transition comparisons with unchanged occupancy components. Each point is one fixed control comparison, not an independent biological replicate: 80 neural and 75 simulated-BOLD comparisons. (c) Composite changes for the two variants; vertical marks are means and points retain every comparison. The directed-relational variant preserves candidate ordering in all 40 system-by-seed groups; the conditional-transition variant preserves 38/40. These finite-panel checks do not establish unchanged principal-model rankings. Definitions, missing-origin handling and verification tests appear in Appendix C.6.

**Table R1:**
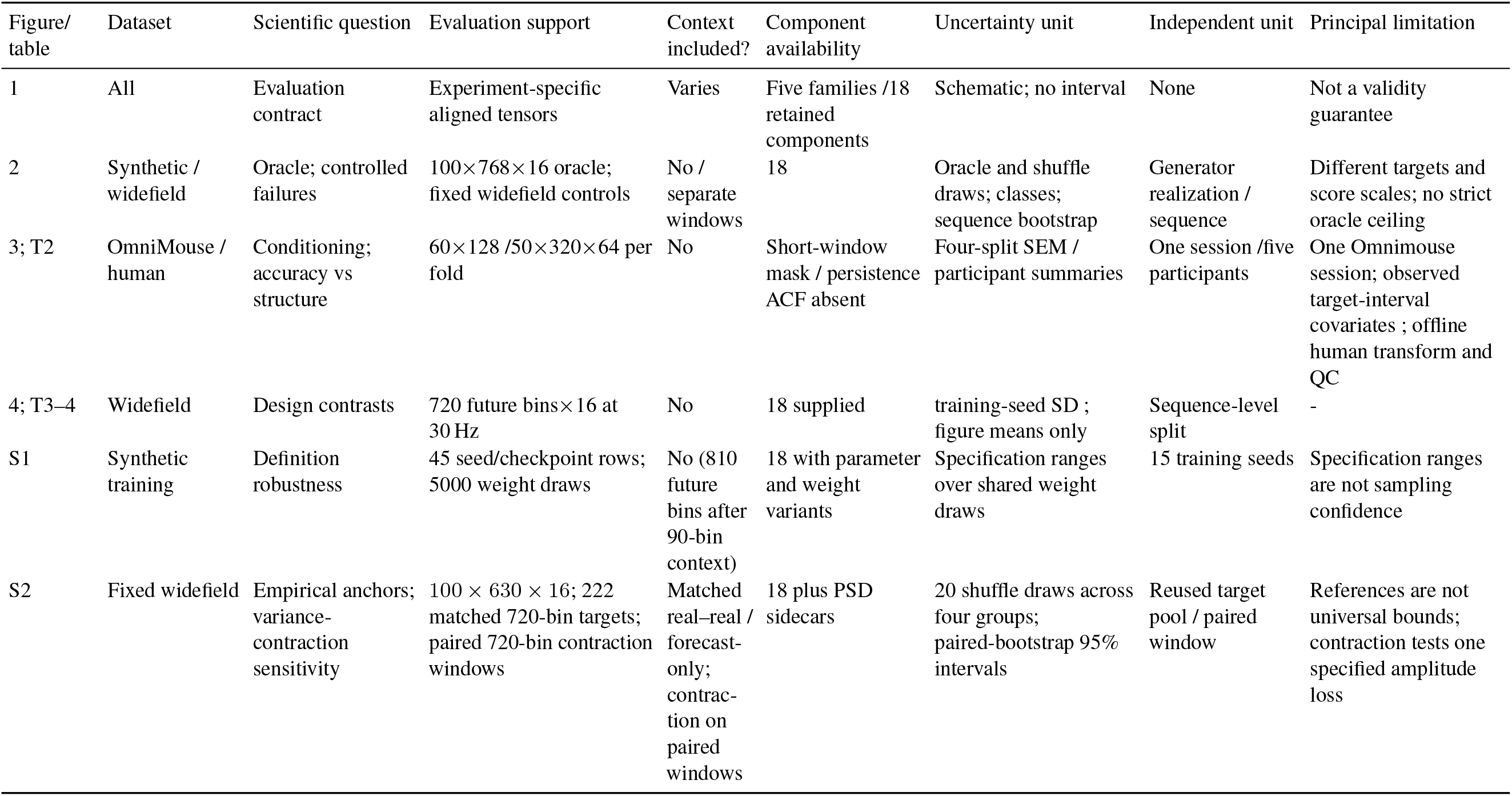
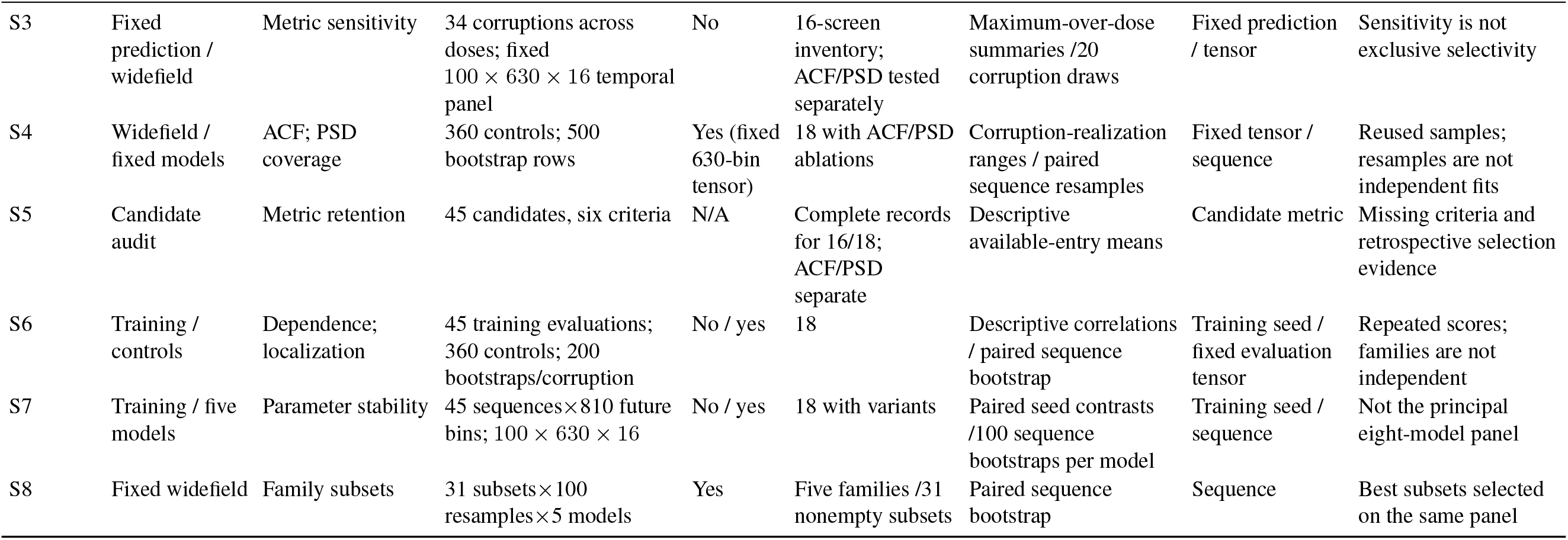
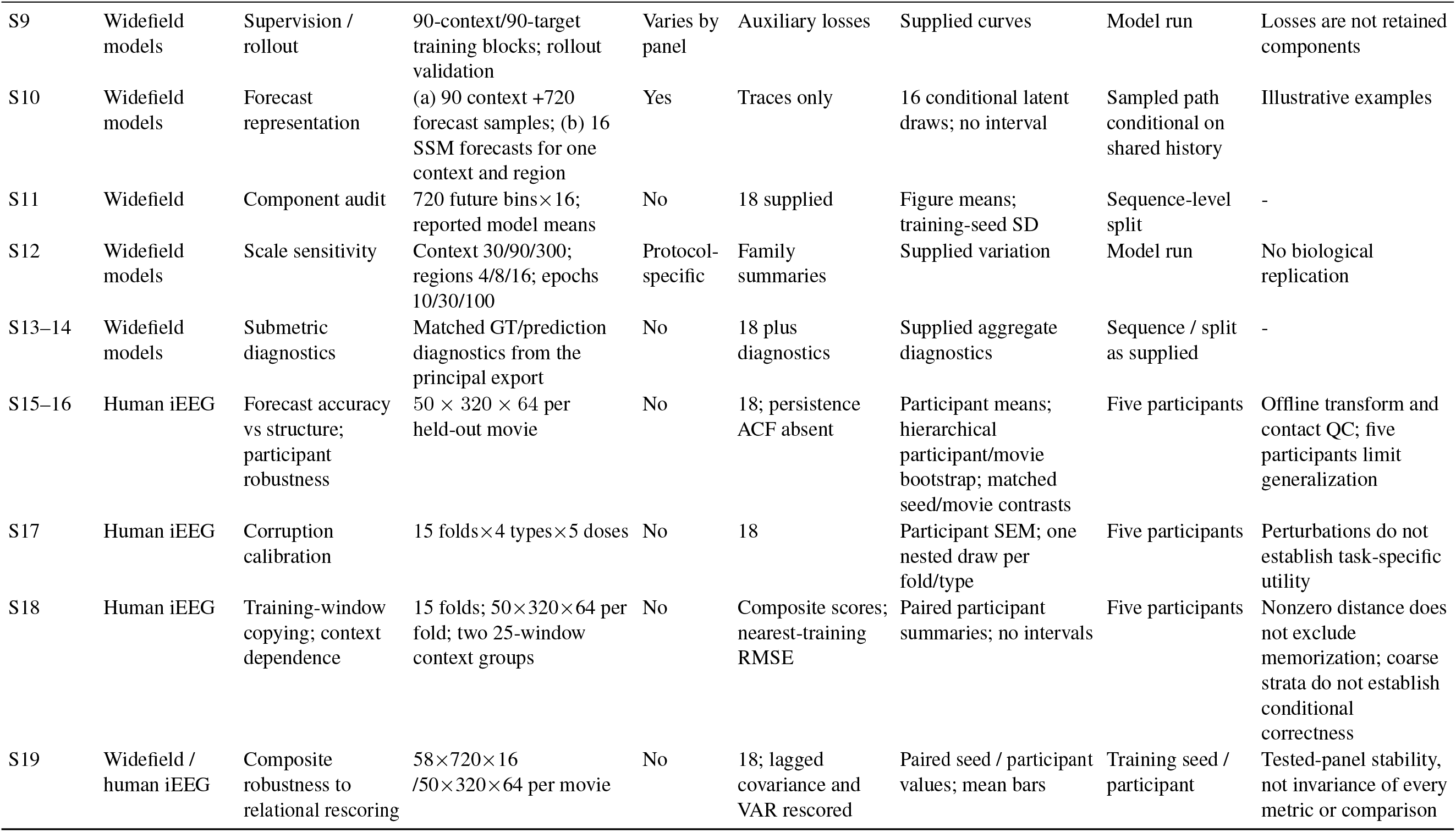
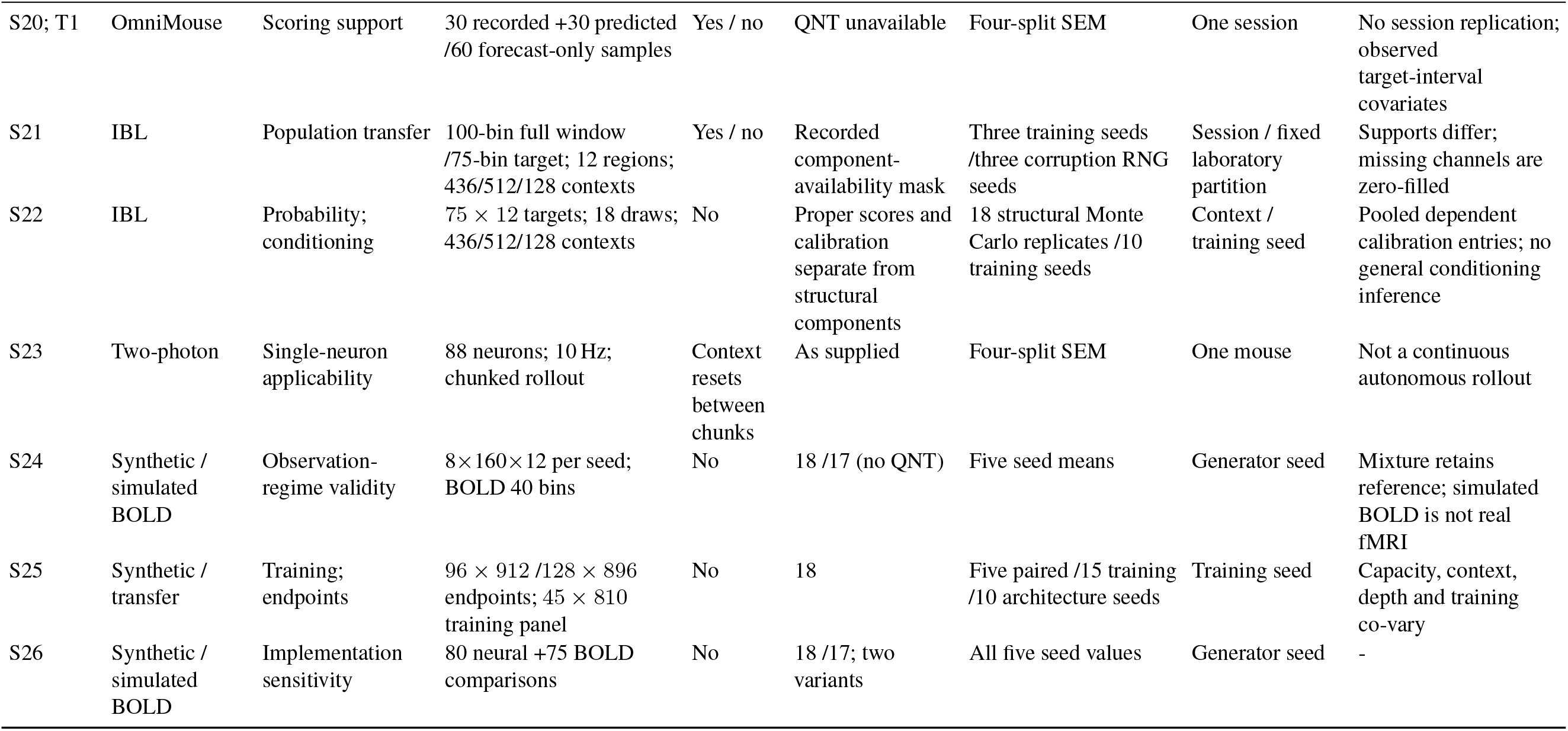
Evaluation supports, units of variation and interpretation limits. T1–T4 denote supplementary tables.

| Figure/table | Dataset | Scientific question | Evaluation support | Context included? | Component availability | Uncertainty unit | Independent unit | Principal limitation |
|---|---|---|---|---|---|---|---|---|
| 1 | All | Evaluation contract | Experiment-specific aligned tensors | Varies | Five families /18 retained components | Schematic; no interval | None | Not a validity guarantee |
| 2 | Synthetic / widefield | Oracle; controlled failures | 100×768×16 oracle; fixed widefield controls | No / separate windows | 18 | Oracle and shuffle draws; classes; sequence bootstrap | Generator realization / sequence | Different targets and score scales; no strict oracle ceiling |
| 3; T2 | OmniMouse / human | Conditioning; accuracy vs structure | 60×128 /50×320×64 per fold | No | Short-window mask / persistence ACF absent | Four-split SEM / participant summaries | One session /five participants | One Omnimouse session; observed target-interval covariates ; offline human transform and QC |
| 4; T3–4 | Widefield | Design contrasts | 720 future bins×16 at 30 Hz | No | 18 supplied | training-seed SD ; figure means only | Sequence-level split | - |
| S1 | Synthetic training | Definition robustness | 45 seed/checkpoint rows; 5000 weight draws | No (810 future bins after 90-bin context) | 18 with parameter and weight variants | Specification ranges over shared weight draws | 15 training seeds | Specification ranges are not sampling confidence |
| S2 | Fixed widefield | Empirical anchors; variance-contraction sensitivity | 100 × 630 × 16; 222 matched 720-bin targets; paired 720-bin contraction windows | Matched real–real / forecast-only; contrac-tion on paired windows | 18 plus PSD sidecars | 20 shuffle draws across four groups; paired-bootstrap 95% intervals | Reused target pool / paired window | References are not universal bounds; contraction tests one specified amplitude loss |
| S3 | Fixed prediction / widefield | Metric sensitivity | 34 corruptions across doses; fixed 100 × 630 × 16 temporal panel | No | 16-screen inventory; ACF/PSD tested separately | Maximum-over-dose summaries /20 corruption draws | Fixed prediction / tensor | Sensitivity is not exclusive selectivity |
| S4 | Widefield / fixed models | ACF; PSD coverage | 360 controls; 500 bootstrap rows | Yes (fixed 630-bin tensor) | 18 with ACF/PSD ablations | Corruption-realization ranges / paired sequence resamples | Fixed tensor / sequence | Reused samples; resamples are not independent fits |
| S5 | Candidate audit | Metric retention | 45 candidates, six criteria | N/A | Complete records for 16/18; ACF/PSD separate | Descriptive available-entry means | Candidate metric | Missing criteria and retrospective selection evidence |
| S6 | Training / controls | Dependence; localization | 45 training evaluations; 360 controls; 200 bootstraps/corruption | No / yes | 18 | Descriptive correlations / paired sequence bootstrap | Training seed / fixed evaluation tensor | Repeated scores; families are not independent |
| S7 | Training / five models | Parameter stability | 45 sequences×810 future bins; 100 × 630 × 16 | No / yes | 18 with variants | Paired seed contrasts /100 sequence bootstraps per model | Training seed / sequence | Not the principal eight-model panel |
| S8 | Fixed widefield | Family subsets | 31 subsets×100 resamples×5 models | Yes | Five families /31 nonempty subsets | Paired sequence bootstrap | Sequence | Best subsets selected on the same panel |
| S9 | Widefield models | Supervision / rollout | 90-context/90-target training blocks; rollout validation | Varies by panel | Auxiliary losses | Supplied curves | Model run | Losses are not retained components |
| S10 | Widefield models | Forecast representation | (a) 90 context +720 forecast samples; (b) 16 SSM forecasts for one context and region | Yes | Traces only | 16 conditional latent draws; no interval | Sampled path conditional on shared history | Illustrative examples |
| S11 | Widefield | Component audit | 720 future bins×16; reported model means | No | 18 supplied | Figure means; training-seed SD | Sequence-level split | - |
| S12 | Widefield models | Scale sensitivity | Context 30/90/300; regions 4/8/16; epochs 10/30/100 | Protocol-specific | Family summaries | Supplied variation | Model run | No biological replication |
| S13–14 | Widefield models | Submetric diagnostics | Matched GT/prediction diagnostics from the principal export | No | 18 plus diagnostics | Supplied aggregate diagnostics | Sequence / split as supplied | - |
| S15–16 | Human iEEG | Forecast accuracy vs structure; participant robustness | 50 × 320 × 64 per held-out movie | No | 18; persistence ACF absent | Participant means; hierarchical participant/movie bootstrap; matched seed/movie contrasts | Five participants | Offline transform and contact QC; five participants limit generalization |
| S17 | Human iEEG | Corruption calibration | 15 folds×4 types×5 doses | No | 18 | Participant SEM; one nested draw per fold/type | Five participants | Perturbations do not establish task-specific utility |
| S18 | Human iEEG | Training-window copying; context dependence | 15 folds; 50×320×64 per fold; two 25-window context groups | No | Composite scores; nearest-training RMSE | Paired participant summaries; no intervals | Five participants | Nonzero distance does not exclude memorization; coarse strata do not establish conditional correctness |
| S19 | Widefield / human iEEG | Composite robustness to relational rescoring | 58×720×16 /50×320×64 per movie | No | 18; lagged covariance and VAR rescored | Paired seed / participant values; mean bars | Training seed / participant | Tested-panel stability, not invariance of every metric or comparison |
| S20; T1 | OmniMouse | Scoring support | 30 recorded +30 predicted /60 forecast-only samples | Yes / no | QNT unavailable | Four-split SEM | One session | No session replication; observed target-interval covariates |
| S21 | IBL | Population transfer | 100-bin full window /75-bin target; 12 regions; 436/512/128 contexts | Yes / no | Recorded component-availability mask | Three training seeds /three corruption RNG seeds | Session / fixed laboratory partition | Supports differ; missing channels are zero-filled |
| S22 | IBL | Probability; conditioning | 75 × 12 targets; 18 draws; 436/512/128 contexts | No | Proper scores and calibration separate from structural components | 18 structural Monte Carlo replicates /10 training seeds | Context / training seed | Pooled dependent calibration entries; no general conditioning inference |
| S23 | Two-photon | Single-neuron applicability | 88 neurons; 10 Hz; chunked rollout | Context resets between chunks | As supplied | Four-split SEM | One mouse | Not a continuous autonomous rollout |
| S24 | Synthetic / simulated BOLD | Observation-regime validity | 8×160×12 per seed; BOLD 40 bins | No | 18 /17 (no QNT) | Five seed means | Generator seed | Mixture retains reference; simulated BOLD is not real fMRI |
| S25 | Synthetic / transfer | Training; endpoints | 96 × 912 /128 × 896 endpoints; 45 × 810 training panel | No | 18 | Five paired /15 training /10 architecture seeds | Training seed | Capacity, context, depth and training co-vary |
| S26 | Synthetic / simulated BOLD | Implementation sensitivity | 80 neural +75 BOLD comparisons | No | 18 /17; two variants | All five seed values | Generator seed | - |

## Notes

https://huggingface.co/datasets/Nethobench/nethobench-widefield-v1

### Competing Interest Statement

The authors have declared no competing interest.

### Summary of Updates

- New validation experiments conducted and reported - Revised text

